# Cryo-EM structure and functional landscape of an RNA polymerase ribozyme

**DOI:** 10.1101/2022.08.23.504927

**Authors:** Ewan K.S. McRae, Christopher J.K. Wan, Emil L. Kristoffersen, Kalinka Hansen, Edoardo Gianni, Isaac Gallego, Joseph F. Curran, James Attwater, Philipp Holliger, Ebbe S. Andersen

## Abstract

The emergence of an RNA replicase capable of self-replication is considered an important stage in the origin of life. RNA polymerase ribozymes (PR) including a variant that uses trinucleotide triphosphates (triplets) as substrates have been created by *in vitro* evolution and are the closest functional analogues of the replicase but the structural basis for their function is poorly understood. Here, we leverage single-particle cryo-EM and high-throughput mutation analysis to obtain the structure of a triplet polymerase ribozyme (TPR) apoenzyme and map its functional landscape. The TPR cryo-EM structure at 5-Å resolution reveals an RNA heterodimer comprising a catalytic and an inactive accessory subunit, where the complex resembles a left hand with thumb and fingers at a 70° angle. The two subunits are connected by two distinct kissing-loop (KL) interactions that are essential for polymerase function. Our combined structural and functional data suggest a model for templated RNA synthesis by the TPR holoenzyme whereby heterodimer formation and KL interactions preorganize the TPR for optimal template binding and templated RNA synthesis activity. These results provide a foundation for a better understanding RNA’s potential for self-replication.

## Introduction

RNA catalysts (ribozymes) occupy central structural and catalytic roles in the function of modern cells including tRNA processing (RNaseP), mRNA splicing (spliceosome, group I / II self-splicing introns) and translation (ribosome peptidyl transferase center) (1). In addition, a much wider variety of ribozyme activities not found in nature has been discovered by *in vitro* evolution, including polymerase ribozymes (PR) that are capable of synthesizing a complementary strand on an RNA template (2-10). Their capacity for RNA-catalyzed RNA-templated synthesis and replication may give rise to a replicase activity enabling RNA self-replication, a process postulated as a central pillar of life’s first genetic system (11, 12).

The earliest examples of nascent PR activity were found in self-splicing intron (SSI) ribozymes, in particular a variant of the *sunY* SSI ribozyme, which allowed single nucleotide triphosphate (NTP) extension (13) or the iterative ligation of RNA oligonucleotides on a complementary strand (14) including assembly of one of its subunits from RNA oligonucleotides (15, 16). The same *sunY* SSI ribozyme was also shown to incorporate short RNA trinucleotide substrates (17), but with relatively low fidelity.

The *in vitro* evolution of the class I ligase (cIL) ribozyme (18) led to a more fully developed PR activity (19), which after further optimization could incorporate up to 14 NTPs in a template-dependent manner (2). The polymerase activity of this first “true” PR was progressively improved by *in vitro* evolution to enable the synthesis of long RNAs (100-200 nts on some RNA templates) (4, 7) as well as the synthesis of functional RNAs including a hammerhead ribozyme (3), tRNA (5), Broccoli fluorescent aptamer (10) and the progenitor cIL ribozyme itself (8). Recently, a variant utilizing trinucleotide triphosphates (triplets) as substrates (a triplet polymerase ribozyme (TPR)) emerged as a heterodimer from *in vitro* evolution (10). This TPR displayed a remarkable ability to copy structured RNA templates including segments of its own sequence (10) as well as circular RNA templates by rolling circle synthesis (20).

However, our understanding of PR function is encumbered by a lack of structural information beyond the progenitor cIL ribozyme (18, 21, 22). While the cIL crystal structures provided insights into the mechanism of phosphodiester bond formation and cIL interaction with the RNA substrate, it is unclear to what extend these features are retained in PRs, which diverge from the cIL not only by a number of mutations in the ribozyme core, but also by 5’- and 3’-extension sequences. A better understanding of how PRs perform accurate substrate selection, general RNA template interaction, and templated RNA synthesis, would therefore benefit from the structure of an active PR.

Here, we report the cryo-EM structure of the complete, heterodimeric TPR apoenzyme determined at its optimal functional magnesium concentration of 100 mM, together with a comprehensive fitness landscape of TPR function. Our results reveal the molecular anatomy of the two polymerase subunits and the geometry and functional importance of their mutualistic association. Our fitness landscape analysis provides a fine-grained mapping of nucleotides important for polymerase function, which together with the structural data define the functional core domains of the TPR and provide the foundation for a model for the TPR holoenzyme consistent with all functional data.

### Cryo-EM structure of optimized TPR heterodimer

In order to improve activity and stability of the original t5+1 TPR (10), we executed further rounds of *in vitro* evolution using an adaptation of a previously described tethered template selection scheme (10) (fig. S1a). Two mutations in t5 (ΔU38 and C110U) were identified and combined with 3 more t5 mutations (U117C, U132C, U148A) discovered in separate selection experiments (to be described elsewhere) (fig. S1b). The resulting t5 variant, 5TU (t5: ΔU38, C110U, U117C, U132C, U148A) exhibited superior triplet polymerase activity compared to t5 (fig. S1c) and remained receptive to activity enhancement by the t1 accessory subunit to copy longer templates (Fig. 1a,b).

**Fig. 1.**
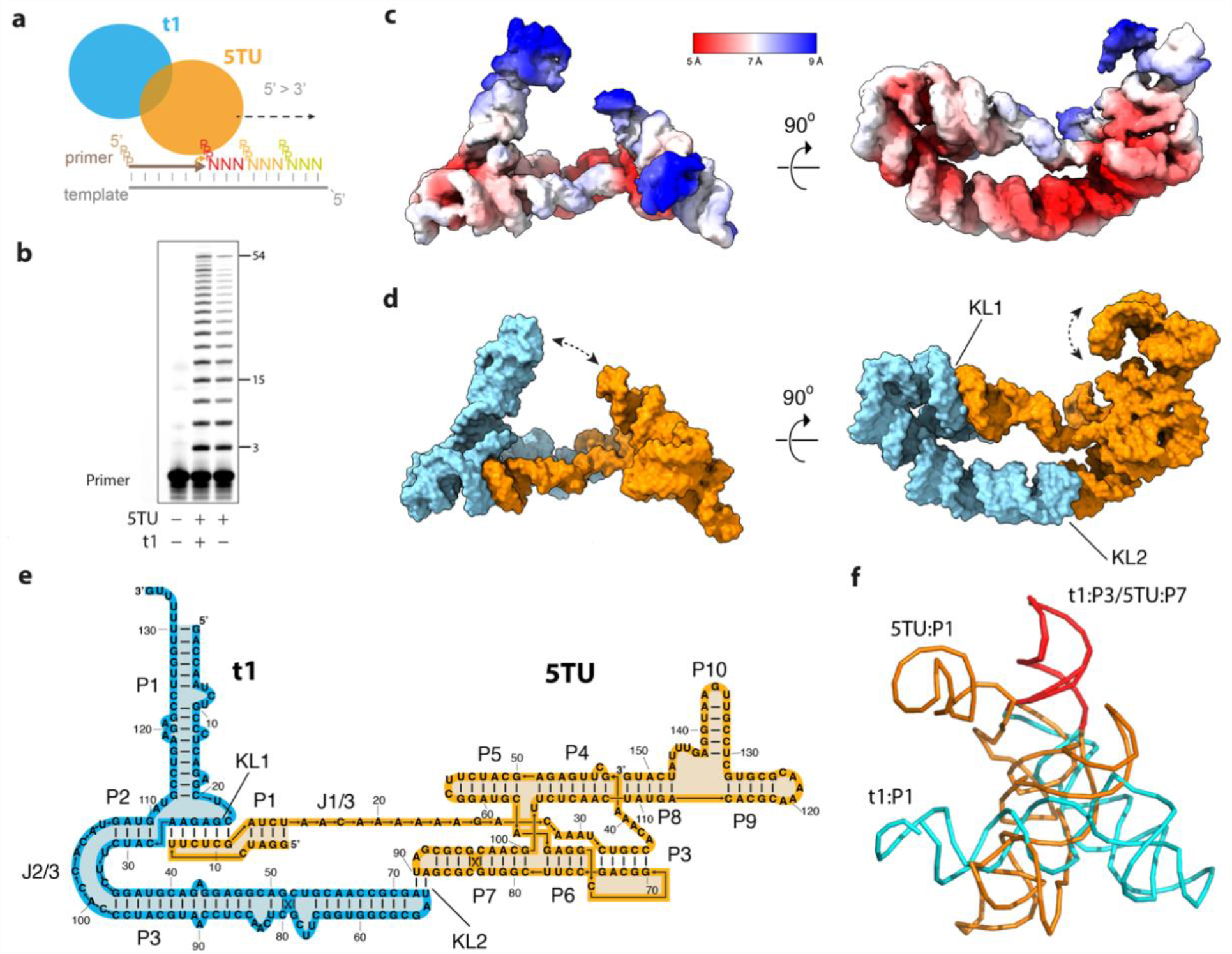
Structure of the Triplet Polymerase Ribozyme (TPR). (a) Schematics of the TPR heterodimer consisting of the catalytic subunit 5TU and scaffolding subuint t1 acting on primer, template and triplet substrates. (b) Polymerization activity of 5TU alone or in combination with t1 in copying a template encoding for (GAA)_18_ after 15 hours. (c) Cryo-EM reconstruction at 5 Å global resolution shown in two perpendicular views colored by local resolution estimates. (d) Atomic model in surface representation shown in two perpendicular views colored by subunit: 5TU (orange), t1 (cyan). Main modes of movement are indicated by double-headed dashed arrows. (e) Secondary structure diagram for the TPR heterodimer consisting of subunits 5TU (orange) and t1 (cyan). Helix domains (P), kissing loops (KL) and longer joining regions (J) are annotated. (f) Structural alignment of t1 P3 and 5TU P7 stems shows major structural divergence between the two subunits.

Cryo-EM was used to investigate the structure of the TPR heterodimer 5TU+t1 in its active form at an optimal Mg^2+^ concentration of 100 mM (see Methods and Table S1). A heat-annealed sample was investigated initially providing a 5.9 Å map (fig. S2-3) but a cotranscriptionally folded sample was later found to provide a 5.0 Å map (fig. S4-6). The map had local resolution carrying from 5-9 Å showing that some domains are highly flexible (Fig. 1c), which was verified by 3D variability analysis (3DVA) (fig. S7-8). The 5TU and t1 subunits were found to have distinct shapes, which were surprising since they are derived from the same core domain. Secondary structure prediction revealed an alternative fold of the t1 subunit (fig. S9-10), which allows for unambiguous placement of all double helices (P) in the cryo-EM map. The remaining joining (J) and loop (L) regions were assembled *de novo* using DRRAFTER (23) (fig. S11). The final model of the heterodimer was refined using molecular dynamics and energy minimizations (see Methods) and reached a map-to-model cross correlation of 7.3 Å at FSC=0.5 and 5.6 Å at FSC=0.143 (Fig. 1d, fig. S13). The model can be described in a secondary structure diagram that shows helix regions and kissing-loop (KL) interactions (Fig. 1e) and a map including tertiary interactions (fig. S14).

The model revealed the overall structural anatomy of the TPR to resemble an upturned left hand, with the thumb formed by the t1 subunit and fingers formed by the 5TU subunit at an approximate angle of 70^°^, with the palm formed by a bipartite interaction of the subunits through two distinct kissing loops (KL1, KL2) (Fig. 1d). The 5TU subunit comprises the catalytic core domains P3-7, the template binding strand J1/3, and peripheral domains P1+P8-10. In contrast, the non-catalytic accessory subunit t1 adopts an extended secondary structure that contains only three main stem domains P1-3. Thus, the structure of the 5TU and t1 subunits has diverged radically, which is surprising since both subunits are derived from the same starting sequence and have only diverged by 7 mutations in the core region (fig. S10a). The only common structure left is a 22-bp segment corresponding to the apical hairpins of the 5TU:P7 and t1:P3 domains, which is involved in the symmetrical KL2 interaction (Fig. 1f, fig. S15).

Further analysis of the cryo-EM data shows that local refinement of the two subunits does not lead to improvements in the resolution (fig. S5-6) suggesting that the bipartite KL interaction is rigid or that the subunits have similar internal flexibility. Two major conformations of the t1 P1 stem were isolated during particle classification (fig S8) and 3D variability analysis (3DVA) revealed a directional movement of both t1:P1 and 5TU P10 domains towards the active site (arrows in Fig. 1d, Movie S1). These movements may have relevance for the mechanism of primer extension by the TPR and will be discussed later. We further investigated the progenitor t5+1 ribozyme (10) at lower (25 mM) Mg^2+^ concentrations and obtained an independently determined 8 Å resolution map that show the same general shape and conformation (fig. S16) suggesting that the 5TU+t1 structure had not significantly diverged from t5+1 and that Mg^2+^ concentration did not affect the general shape.

### Functional landscape of TPR heterodimer

To connect structural features in our heterodimer model to TPR function, we performed a comprehensive fitness landscape (24, 25) analysis in triplicate (Methods, Fig. 2 and figs S17-22, Movie S2) by quantification of changes in genotype abundance pre- and post-selection; we define ribozyme “fitness” as the log-transformed enrichment of a given genotype relative to the wild-type (wt) 5TU or t1 sequence. After filtering, we obtained relative fitness values for 128,708 ribozyme variants, comprising 79,702 5TU and 49,006 t1 genotypes, providing fitness estimates of all t1-as well as 99.6% of 5TU-point mutants. For both subunits, calculated fitness was strongly correlated across replicates (Pearson coefficient R= 0.89 (5TU) / 0.95 (t1), and R = 0.97 (5TU) / 0.95 (t1) if only single and double mutants are considered) (Fig. 2a, fig. S17b).

**Fig. 2.**
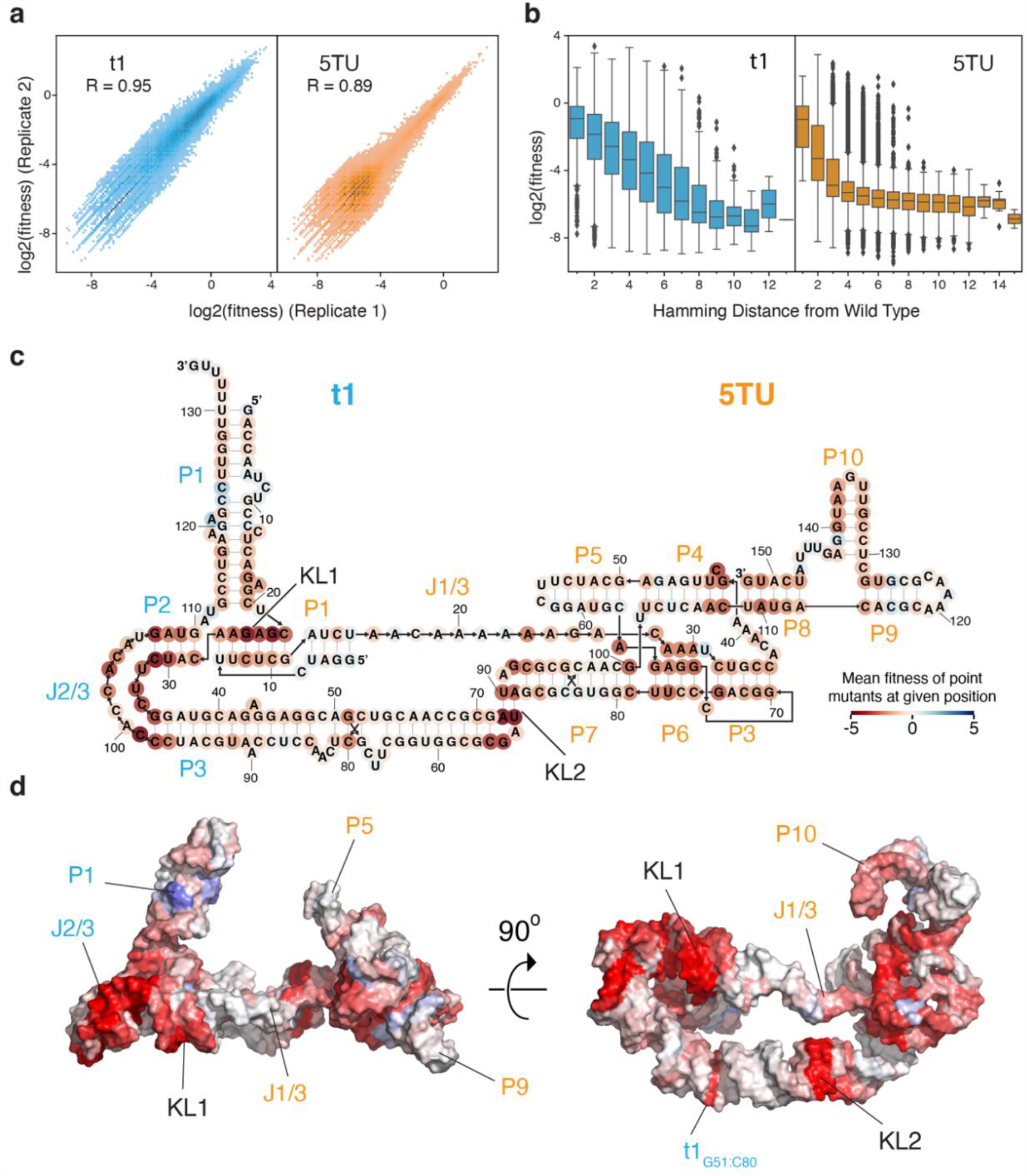
Fitness landscape of the TPR. (a) Reproducibility of fitness values between two replicates of t1 (cyan) and 5TU (orange) (for other replicates see fig. S17). (b) Fitness values as a function of Hamming mutational distance for t1 (cyan) and 5TU (orange) (n=3). (c,d) Average fitness values for a given nucleotide position in TPR secondary structure (c) and tertiary structure (d) (see Movie S2).

Next, we analyzed the dataset for global properties and concordance with established TPR function (Fig. 2a). While mean fitness of both 5TU and t1 mutants was negatively correlated with Hamming distance from wt sequences (Fig. 2b), the fitness decline was noticeably steeper for 5TU than t1. Furthermore, while the majority of 5TU genotypes showed a much-reduced fitness compared to wt, the t1 fitness distribution - while also negatively skewed - was considerably flatter (fig. S17a). These results are consistent with the highly evolved catalytic 5TU subunit occupying a steeper fitness peak (in a more rugged adaptive landscape) compared to the more recently evolved t1 accessory subunit.

Fitness landscape analysis further revealed the functional relevance of both known structural features of functional importance in cIL (21) and novel structural features that are unique to the TPR (Fig. 2c,d, fig. S18). Known structural features include: the template-binding nucleotides in J1/3 (positions 23-24), the active site cytidine in P4 (position 43), and the P6 triple helix-forming adenosines (positions 28-30). Novel features of importance to TPR function include: the P10 stem (positions 137-140), the kissing-loop interactions (KL1 and KL2) between the two subunits, as well as the internal loop region of t1 J2/3,J3/2 (positions 99-106, 32-34) and a G-C base pair (bp) in t1 P3 (position 51 and 80). These will be discussed below in relation to the structural analysis.

Analyzing double mutants, we found that significant epistatic interactions in both 5TU and t1 were negatively biased (fig. S19-20) and rarer in t1 than in 5TU. Moreover, as the physical distance between residues increased (as calculated from our structural model), both the proportion of significant epistatic interactions and the magnitude of epistasis, decreased in both subunits (fig. S21b). Finally, we found that the average epistatic value decreased as the fitness of the first point mutation increased in double mutants of both 5TU and t1 (fig. S21a). All of these trends are consistent with previously determined fitness landscapes of a yeast tRNA (26), and snoRNA (27), suggesting that they may represent general features of RNA structure and evolution.

Although our dataset does not comprehensively capture all double mutants in either ribozyme subunits, many double mutants at predicted base-pairing positions exhibit positive epistasis, (particularly within t1) lending support to our structural model (fig. S22a, Table S2-3). Moreover, at base-pairing positions predicted by our secondary structure models of 5TU and t1, point mutations that result in a wobble base pair were consistently higher in fitness compared to base pair-disrupting point mutations (fig. S22b).

### Structure of the 5TU catalytic subunit

Our structural data and fitness landscape data provides a basis for a better understanding of structure-function relationships in the TPR. The 5TU subunit is a descendant from the class I ligase (cIL) with functionally important sequence extensions at both the 5’ and 3’ ends. Comparison of the cIL crystal structure and the 5TU cryo-EM structures shows that the overall structure of the cIL core region (P3-7) is preserved despite several mutations and the lack of the substrate helices (fig. S23-24). The 5’ extension is composed of a long single strand J1/3 with a hairpin at the 5’ end, which is stabilized by the t1 subunit. The 3’ extension forms 3 stems P8-10, where P8-9 coaxially stack on P4, with P10 protruding on the side of P8-9 (Fig. 3a,b).

**Fig. 3.**
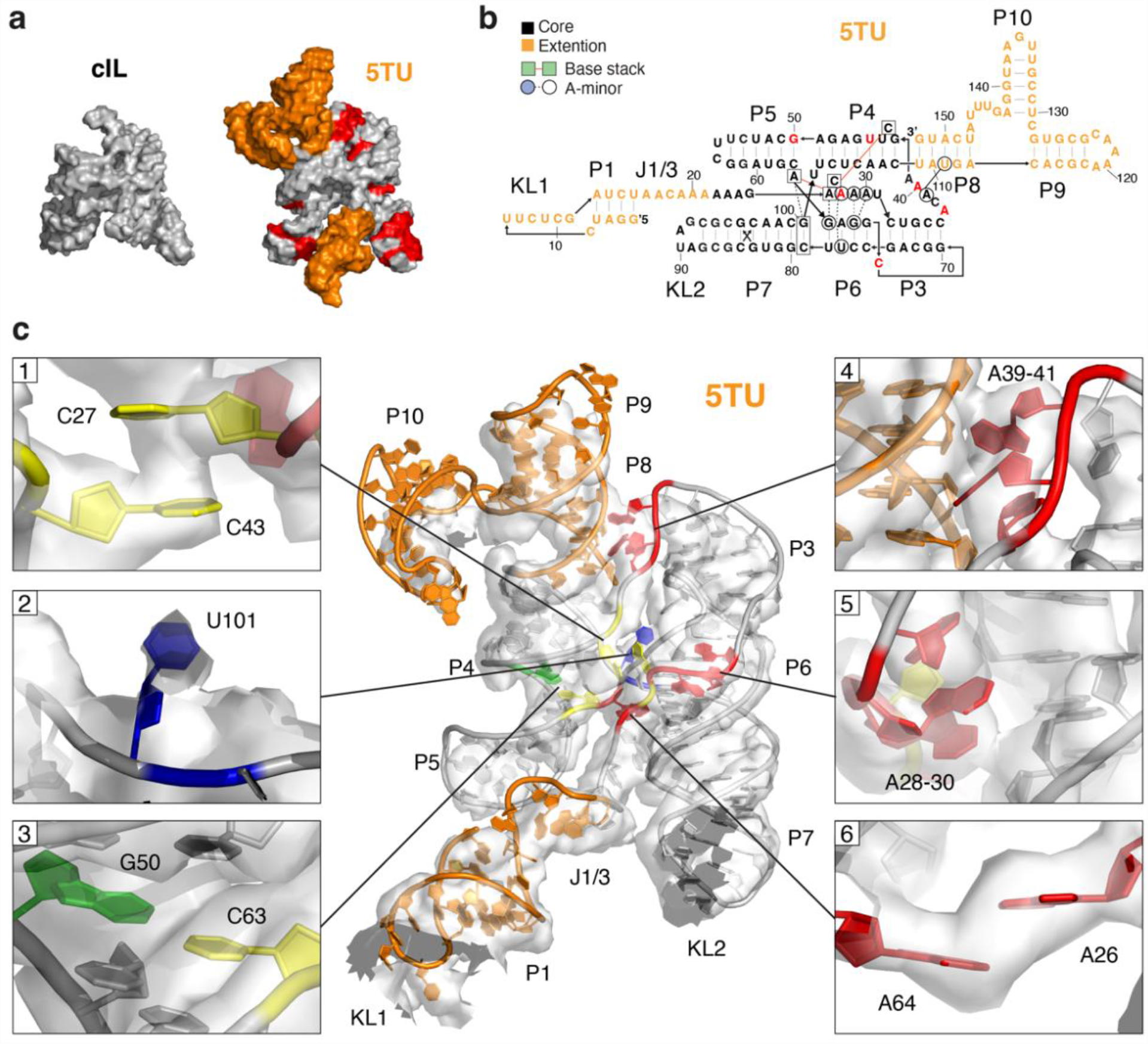
Structural features of catalytic subunit 5TU. (a) Comparison of class I ligase (cIL) and 5TU with the core domain in grey and 5’ and 3’ extensions in orange. P1 and P2 substate domains have been removed from cIL for clarity of comparison. Mutational differences are marked in red on 5TU. (b) Secondary structure model of 5TU showing core domain (black), extension domains (orange), base stacks (boxes) and A-minor interactions (circles). Mutational differences between core domain of cIL and 5TU are marked in red. (c) 5TU atomic model in cryo-EM map with zoom-ins on core features: (1) C27:C43 base stack, (2) flipped out U101, (3) G50:C63 base pair, (4) A-minor interaction of J3/4 with P8, (5) A-minor interaction of J1/3 with P6, (6) A26:A64 base stack. Selected nucleotides are colored by base type: adenine (red), uracil (blue), guanine (green), cytosine (yellow). Contour levels have been set to highlight distinctive features: full view=1; (1)=2; (2)=0.5; (3,4)=4, (5)=6; (6)=11. Full view shows sharpened global map within 5 Å of the 5TU model. Zoomed views show sharpened locally refined map for 5TU core region.

The core region (P3-7) is composed of two parallel helices connected at two positions with approximately one helical turn between connections, which creates a stable scaffold similar to the double crossover motifs known from nucleic acid nanotechnology (28). Notable differences between cIL and 5TU include an anti-parallel crossover connecting P4-P5 and P6-P7 helices, where the junction has shifted in 5TU due to a A50G mutation. The cryo-EM map supports the formation of a G50:C63 (base-pair) bp and a resulting U101 bulge (Fig. 3c, panel 2-3). Indeed, there is epistasis support for the G50:C63 when including G:U wobble pairs (fig. S22b). A second structural divergence from cIL involves the top of the P3 stem, where the cryo-EM map supports an interaction between J3/4 and the minor groove of P8 (Fig. 3c, panel 4), where A39 interacting with the minor groove is supported by mutational data. In cIL this interaction cannot happen and is instead formed by a G:U base stack, which is not present in 5TU (fig. S24).

The 5TU J1/3 single strand is connected to the core region (P3-7) by three connections in a very similar configuration as in cIL A26 of J1/3 forms a base stack with A64 of the P5-P6 junction, which is well supported by both the cryo-EM map (Fig. 3c, panel 6) and fitness data (fig. S18). C27 of J1/3 forms a base stack with the catalytic C43 bulge of the P4 stem, which is well supported by the cryo-EM map (Fig. 3c, panel 1) and fitness data showing that while C27 can be mutated to U, C43 is absolutely conserved (fig. S18). Finally, J1/3 A28-A30 form A-minor interactions with the P6 stem (Fig. 3c, panel 5) in a similar conformation as the A-minor triad in cIL. This A-triad is highly conserved (fig. S18) as is the base pairing of the P6 stem (fig. S22b). A striking feature in our data is the extension of J1/3 into a rigid, single stranded region by interaction with the t1 domain which is well resolved in the cryo-EM map. The second A-minor triad that in cIL interacts with the substrate helix is placed the same spatial position in 5TU suggesting that the template will likely be placed in a similar position.

The 5TU 5’ extension forms a hairpin that forms a kissing loop interaction with the t1 subunit, which is strongly supported by fitness data (fig. S18). The 3’ extension domain forms the P8 stem which branches into P9 and P10 hairpins. P8 forms a coaxial stack with P4 on one side and P9 on the other. P8 109-113 and 148-152 are both highly sensitive to mutations (fig. S18) and supported by epistasis when including G:U wobble pairs (fig. S22b). In contrast, P9 is less sensitive to mutations suggesting a less important role. The loop at the end of P9 is modelled as a A119:C123 trans sugar-Hogsteen bp with an A-stack at the 3’ side of the loop. P10 projects from P8-P9 with a four nt single strand on one side and a direct connection on other side, which results in a perpendicular orientation of the P10 hairpin. The hairpin is 6 bp long and ends in a 3-nt loop (position 135-137) that is modelled as a stack. However, we cannot be sure of the modeling here because the low resolution in this domain. The region around the loop of P10 and the junction around P10 is highly sensitive to mutations (fig. S18) suggesting that P10 is involved in an important functional role. Previously P8-10 has been shown to improve TPR fidelity and proposed to work through minor groove interactions with the triplet substrate (10).

### Structure of the t1 scaffolding subunit

The t1 subunit evolved as a mutualistic parasite to the catalytic domain in the original TPR selection (10) and differs from 5TU through only 7 mutations in the core domain and a distinct 3’ extension sequence (Fig. 4a). The cryo-EM structure reveals that these changes have caused a major refolding of the core domain into a long hairpin which bends back on itself to form a 70-degree angle. The base pairing interactions of the long hairpin is supported by positive epistasis values (fig. S25, Table S3). The t1 structure is composed of three main helix domains: P1 that extends at a 70-degree angle from the plane defined by the P2 and P3 helices. P2 is extended by co-axial stacking of KL1 and P1 of 5TU, whereas P3 is connected through KL2 to P7 of 5TU (Fig. 4b). The three helices together form a stable core region of the t1 subunit that, similar to 5TU, has two connections at an approximate distance of one helix turn.

**Figure 4.**
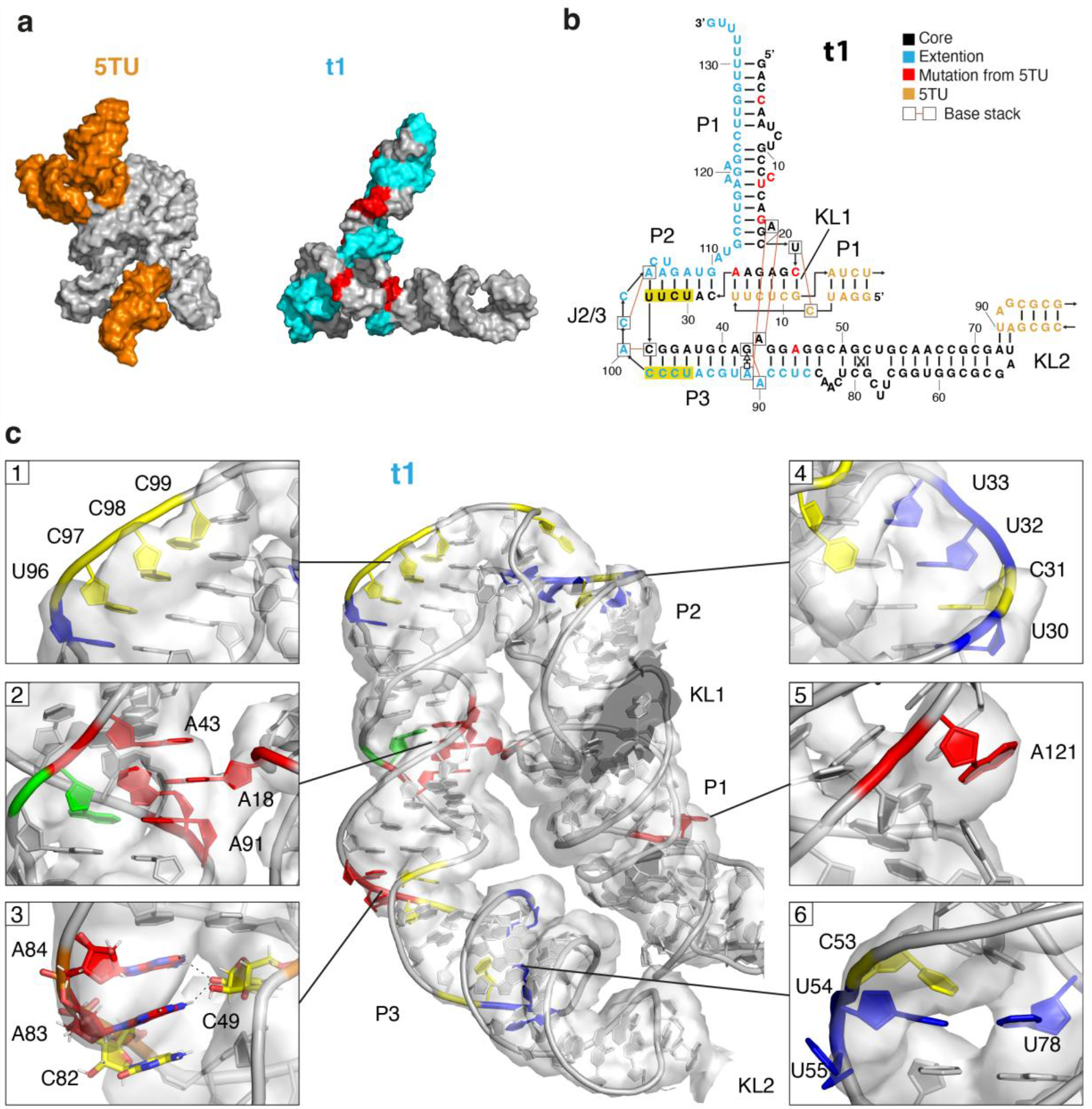
Structural features of scaffolding subunit t1. (a) Comparison of subunits 5TU and t1 in surface representation. 5TU core is colored grey and 5’ and 3’ extensions are colored orange. t1 with sequence corresponding to catalytic core colored in grey and 3’ extension colored in cyan. Mutational differences in core region are colored in red. (b) Secondary structure model of t1 showing core domain (black), extension domain (cyan), base stacks (boxes), 5TU KL elements (orange) and pyrimidine stretches (yellow). Mutational differences between core domain of 5TU and t1 are marked in red. (c) t1 atomic model in cryo-EM map with zoom-ins on core features: (1) 69-99 pyrimidine strech, (2) A18 base stacking in A43 bulge, (3) helix bend at asymmetric bulge, (4) 30-33 pyrimidine strech, (5) helix bend at asymetric bulge, (6) G58:A75 noncanonical base pair. Selected nucleotides are colored by their base type: adenine (red), uracil (blue), guanine (green), cytosine (yellow). Contour levels have been set to highlight distinctive features: full view=1; (1,3,4)=2; (2)=8; (5,6)=4. Full view shows sharpened global map within 5 Å of the t1 model. Zoomed views show sharpened locally refined map for t1 core region.

The first connection is formed by a sharp turn of the P2 and P3 helices at the J2/3 and J3/2 internal loop, which we name the “Py-turn” motif, since it is facilitated by two pyrimidine tracts (positions 30-34 and 96-99) that bend the strands towards each other (Fig. 4c, panel 1,4). The strands are further brought together by the presence of a non-canonical pyrimidine-pyrimidine bp C34:C99. The motif is capped by two bases of the J2/3 region that stacks on the P2 and P3 helices, respectively. Three bases (C102, C104, U106) remain unpaired with distinct density in the major groove of P2 and at the minor grooves of P2 and P3. The presence of this structural motif is further supported by the fitness data showing strong conservation of the pyrimidine tracts, and the non-canonical C:C bp, C102 and C104 (fig. S18a).

The second connection is formed by a bulged-out A from P1 that inserts in an internal loop of P3, which we name the “A-anchor” motif. The motif is formed by the base of the P1 helix that supports a bridge between the P2 and P3 helices that together with the Py-turn motif orients them in parallel. The A18 bulge is formed between two C:G bp of P1. A18 stacks between A43 and A91 across the minor groove, which is seen clearly in the EM map (Fig. 4c, panel 2). The alternative position of A91 is facilitated by a trans-Hogsteen-sugar bp between G42:A91. Density in the major groove further suggests that A90 stacks on G42 across the major groove. The A-anchor motif is supported by a positive epistatic interaction between A18 and A43 and for base pairs around the symmetric bulge (fig. S25) and mutations to any nucleotide causes an intermediate reduction in fitness (fig. S18a).

From the A-anchor motif, the P3 stem has two other bulge regions that causes the stem to bend approximately 120 degrees towards KL2. The first bend is formed by C82-A84 across from C49 (Fig. 4c, panel 3) and the second bend is formed by C53-U55 across from U78 (Fig. 4c, panel 6). Together these bend the P3 across the major groove in-between the motifs. Mutations to these bulge regions do not affect fitness (fig. S18a), suggesting that the precise base composition is not important, whereas the position of 3 bases across from one may suffice for the bending. Interestingly, a G51:C80 bp in the stem between the two bulges is heavily affected by mutation and can be partly rescued by mutation to wobble pair G:U (fig. S18a). In P3 there is a non-canonical G58:A75 bp, which is also preserved in the 5TU P7 as G81-A98. However, the fitness data does not suggest that this G:A bp is important for function (fig. S18a,b).

The P1 stem is well resolved in the cryo-EM map near KL1 and the A-anchor motif, but less resolved in the top part. Density is observed at A121, which seem to bulge out of the helix between two G:C bps (Fig. 4c, panel 5) in a similar fashion to the A-anchor motif. Mutations in this region do however seem to have both positive and negative effects on fitness (fig. S18a). Double mutant epistasis in general support the formation of the P1 stem with positive epistasis for position 13 and 119 indicating a base pair, while positions 8 and 124 show negative epistasis supporting the bulge (fig. S25). The t1 P1 helix appears to be supported at its base by two key tertiary interactions (the KL1 bKL and A-anchor) that form a hinge allowing the large dynamic movement of t1 P1 (Movie S1). Because of the orientation of the hinge, the movement of the t1 P1 is towards the 5TU active site with potential functional implications discussed below.

### Structure and function of bipartite kissing-loop linkage

A striking feature of the TPR structure is that the two divergent subunits are connected through two distinct kissing-loop (KL1, KL2) interactions (Fig. 5a). The geometry created by these two KL interactions enforces a rigid, extended conformation of the single stranded 5TU J1/3 segment that is clearly visible in reconstructions from all our datasets. Importantly, heterodimer formation is essential for full triplet polymerase activity (Fig. 1b) and the primer/template interaction enabling RNA synthesis activity without template tethering, which is obligatory for most other polymerase ribozymes (3).

**Fig. 5.**
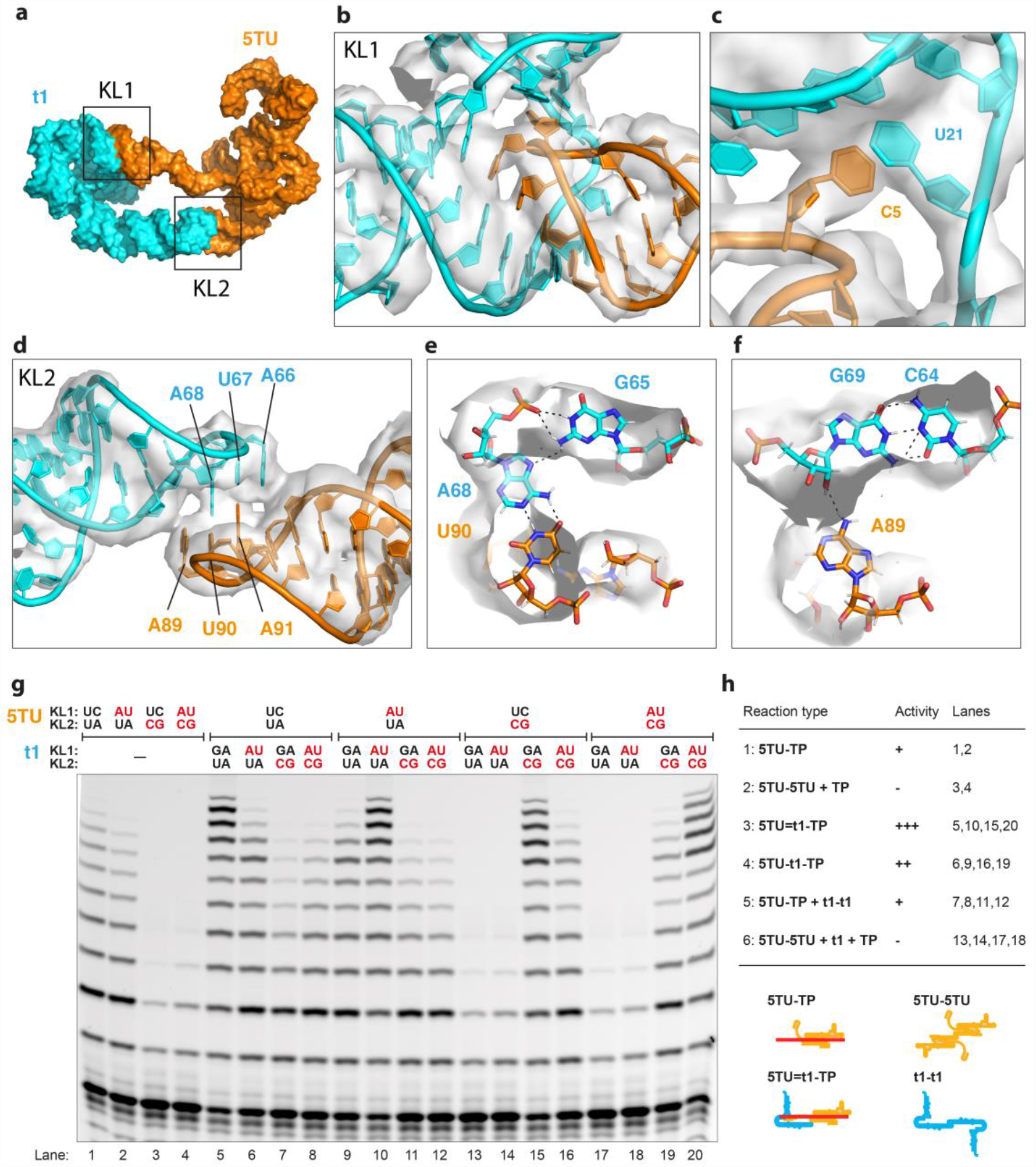
Structure and function of bipartite kissing-loop interaction. (a) TPR heterodimer model shown as surface representation with 5TU in orange and t1 in cyan. KL1 and KL2 indicated by boxes. (b) Atomic model of KL1 in sharpened locally refined EM map for t1 core region shown at contour level 8. (c) Zoom on KL1 supporting C5:U21 base stack. (d) Atomic model of KL2 in sharpened global EM map shown at contour level 4. (e) Structural detail of KL2 showing U90:A68 base pair and A68:G65 noncanonical interaction. (f) Structural detail of KL2 showing G69-C64 base pair and G69-A89 noncanonical interaction. (g) TPR primer extension activity of wild-type as well as mutant KL sequences visualized by gel electrophoresis. Mutation of 5TU KL1 and in particular KL2 reduce TPR activity, but activity can be restored by compensating mutations in cognate loops in t1. (h) Overview of reaction type, activity, and associated lanes from gel in panel g. Schematics showing the proposed complexes that can form depending on homodimer or heterodimer formation.

The structure of KL1 shows a 6-bp interaction between the loop of the 5TU P1 hairpin and the t1 J1/2 internal loop, which results in a coaxial stack of 5TU P1, the KL, and the t1 P2 stems (Fig. 5b). The interaction is reminiscent of a branched KL (29) with several similar features like the bridging over the major groove. The 5TU C5 that bridges the major groove is observed to base stack with t1 U21 (Fig. 5c), while the two single stranded bases A111 and U112 stack underneath the P1 stem. The KL1 base pairing between 5TU U6-G11 and t1 C22-A27 shows a clear functional signal, since mutation in these regions are detrimental, and mutations to the wobble G:U are less severe (fig. S26a). Mutations of the (5TU C5) : (t1 U21) base stack does not show a strong effect on fitness (fig. S26a).

The structure of KL2 is a 2-bp loop-loop interaction between the identical apical loops of 5TU P7 and t1 P3. The 5TU P7 and t1 P3 have identical terminal GAUA loops as a consequence of the shared evolutionary ancestry between 5TU and t1 (fig. S10). The KL2 structure is found to have high structural similarity to the GACG kissing loop of the retrovirus MoMuLV determined by NMR (30), which is a palindromic/symmetrical interaction involved in homodimerization of retroviral genomes. The difference is that the 2-bp interaction is formed by two A:U bps in KL2, whereas it is formed by two G:C bps in the MoMuLV KL. The central A:U base pairs are stabilized by hydrogen bonding to the first G of the tetraloop (Fig. 5e) and the second A of the tetraloop stack on the kissing base pairs and form an additional inter-strand hydrogen bond to the 2’O of the G:C bp of the stem (Fig. 5f). Because of the symmetry of the homodimerization, the fitness of 5TU and t1 point mutants in the two loops and first bp of the stems are virtually identical, with mutations to G:U being less severe (fig. S26b).

To investigate the role of the KLs on TPR activity we introduced targeted mutation into both KL1 and KL2 interactions and analyzed their functional impacts by primer extension assays (Fig. 5g). In the absence of t1, mutation of 5TU KL1 did only marginally reduce polymerase activity (Fig. 5g, compare lane 1,2), whereas mutation of 5TU KL2 strongly inhibited primer extension (lane 3,4). We hypothesize that this may be due to formation of 5TU KL2-CG homodimers that inhibit primer extension (Fig. 5h, reaction type 1,2). The interaction of wt t1 and wt 5TU provides the expected boost of primer extension activity (lane 5, reaction type 3). When KL1 is disrupted by an AU mutation, we see a reduced activity that can be explained since KL2 is still able to form the heterodimer (reaction type 4). When t1 KL2 is disrupted by mutating to CG, we see a reduction of activity to that of 5TU alone, which may be explained by t1 forming strong homodimers and does thus not contribute to the activity boost (reaction type 5). When 5TU KL2 is disrupted by mutating to CG, we see strong inhibition of activity, which may be explained by strong 5TU homodimer formation (reaction type 6). Interestingly, compensatory mutations restoring the KLs leads to wildtype activity by reconstituting the heterodimer (reaction type 3). Thus, mutational analysis confirms the importance of the cognate KL1 / 2 interactions in promoting formation of the fully active TPR heterodimer as suggested by the structure. Furthermore, it highlights the importance of the bivalent KL interaction in favoring the correct heterodimer over possible poorly active homodimers that can be formed through the symmetric KL2 interaction (Fig. 5h).

### Model of holoenzyme and effects of functional elements

To explore the functional implications of the TPR structure we sought to build a model of the holoenzyme. First, the catalytic 5TU subunit was aligned to the cIL crystal structure (21) to build a putative template-product helix by analogy to the cIL P1 substrate helix (Fig. 6a, fig. S27-28). This simple model remarkably positions the primer-template duplex, the triplet substrate 5’-triphosphate group, and incoming triplet substrate minor groove close to interacting features of the 5TU subunit (J1-3 segment, active site, and P10 domain). Additionally, it situates the elongated nascent strand/template duplex near the t1 P1 helix (Fig. 6b,c).

**Fig. 6.**
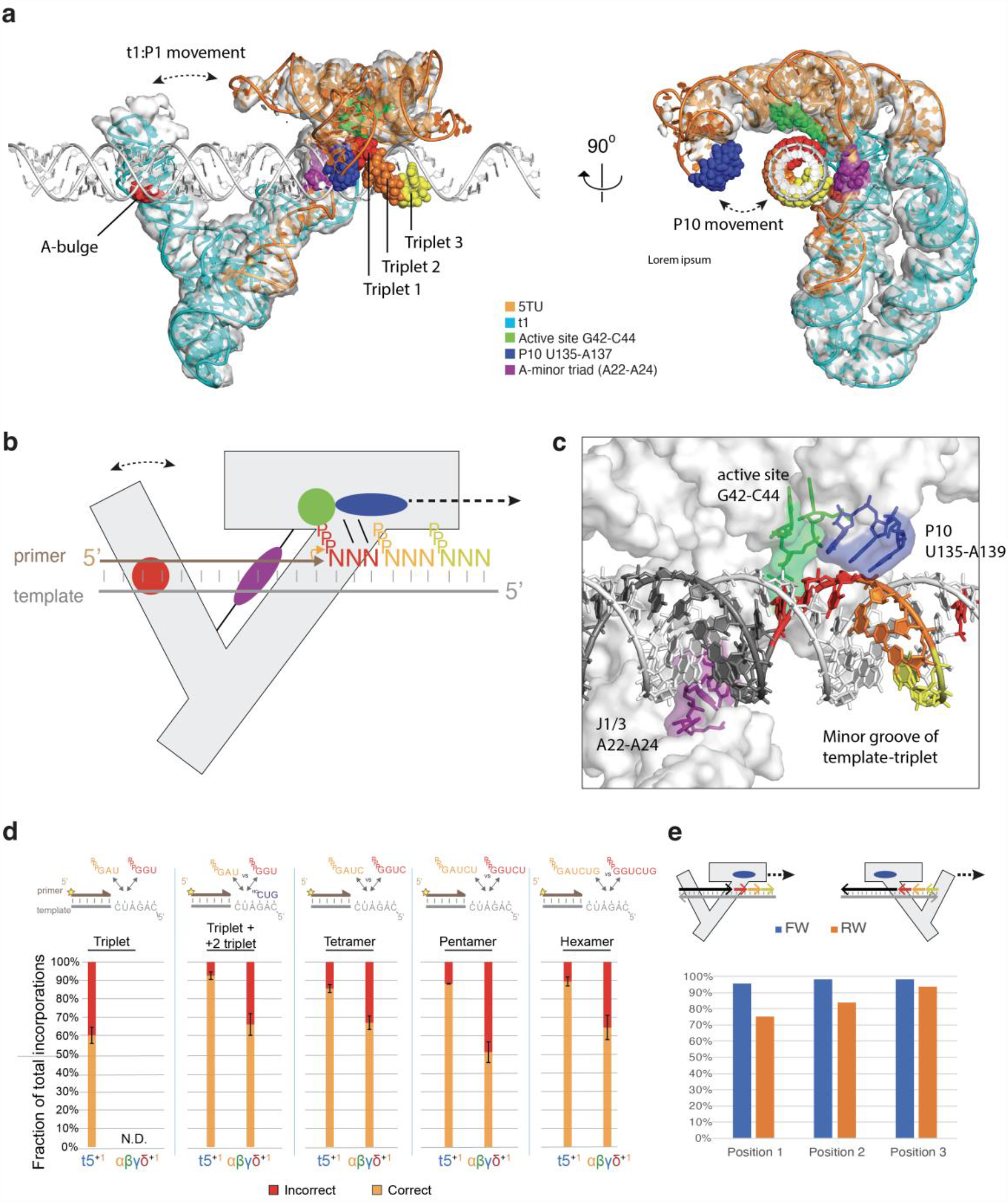
Structural model of the TPR holo-enzyme and role of P10 proof-reading. (a) TPR model (5TU (orange), t1 (cyan)) in EM contour map with an idealized double-stranded RNA template (white) shown in two perpendicular views. The model was constructed by aligning the class I ligase structure to 5TU followed by aligning the template to the class I ligase substrate helix. Core elements are highlighted as spheres: P10 (blue), J1/3 (purple), active site (green), triplets (red, orange, yellow). t1 P1 and P10 movements from 3DVA are shown as dashed double headed arrows. (b) Schematic model of TPR model with RNA template and triplets. Putative contact sites are shown: active site C43 (green), J1/3 A-minor interaction (purple), P10 minor groove interaction (blue), t1 interaction (red). (c) View of the RNA template in the active site shown at an angle where minor and major grooves are visible. Shows putative minor groove interactions with primer-template helix and template-triplet helix. Colors similar to panel b. (d) TPR substrate contacts and fidelity as fraction of correct to incorrect substrate incorporation for substrates of increasing length. abgd+1 have missing P10 domain. For “triplet +2 triplet” and pentamer n=2. For all other conditions n=3. Mean ratios are shown, with standard deviations. (e) Fidelity of different polymerisation modes. Schematic of 5’ to 3’ forward and 3’ to 5’ reverse polymerisation. +1 and -1 triplets shown in red, +2 in yellow and +3 in pale green. Fidelity profiles of forward (blue) and reverse (orange) triplet incorporations, as determined by FidelitySeq.

The extended and rigid conformation of the single-stranded J1/3 linker segment is a noteworthy and unanticipated feature of the TPR structure. J1/3 is of particular interest because the equivalent positions to 5TU A22-A24 are implicated in A-minor interaction with the substrate helix in the cIL structure (21). Indeed, our TPR model positions the template-product helix in proximity to J1/3 (Fig. 6c). Furthermore, functional data suggests that an extended A-minor triad conformation is essential for full TPR function with even 2-nt insertion or deletions in J1/3 reducing TPR activity to baseline (fig. S29). Thus, the t1 domain and its KL interactions may serve to jointly stabilize J1/3 in an out-stretched conformation. This may enhance template-primer duplex interactions by reducing J1/3 conformational freedom and secondary structure formation, reducing the entropic cost of template interaction compared to an untethered strand. Analysis of the evolution of the related 52-2 polymerase ribozyme (which used NTPs as substrates) (6) suggested the emergence of a pseudoknot structure involving P7 and the J1/3 equivalent, which might enhance PR activity via a similar restriction of the conformational freedom of this crucial sequence segment.

A notable feature of the TPR observed previously is its fidelity of 97% (per nucleotide position) (10). A significant contribution was ascribed to the P10 (formerly epsilon (10)) domain that - by H-bonding with the minor groove of the 3’ base of the incoming triplet - appears to enhance fidelity compared to 92% of a TPR variant that lacks P10 (10). We performed further functional analysis, which suggests, that P10 may make even more extensive interactions (Fig. 6d): When only a single triplet is bound to the template 3’ of the ligation junction, the P10 fidelity boost is lost, but full fidelity is regained in the presence of a second downstream triplet. Even in the absence of a downstream triplet, but using substrates of increasing length, P10-dependent fidelity gains are almost entirely restored using a quadruplet (pppN_4_) substrate, with minimal further fidelity gains with longer (pppN_5_, pppN_6_) substrates. This suggests that P10 forms functionally important contacts with the primer-template duplex extending over at least 4 nts. Indeed, our structural model positions P10 and specifically U135, G136 & A137 in close proximity, poised for interaction with the minor groove of the incoming triplet substrate (Fig. 6c).

Another remarkable feature of the TPR is its capacity to support non-canonical RNA synthesis modes such as triplet polymerization in the reverse 3’-5’ direction (10). Analysis of the 3’-5’ mode of templated RNA synthesis by the TPR using deep sequencing (FidelitySeq, fig. S30) suggests that - in contrast to the 5’-3’ reaction - fidelity is reduced to 84%, even below the baseline fidelity of 5’-3’ synthesis in the absence of the P10 domain (Fig. 6e, fig. S31). Although the measured 3’-5’ error rate may be inflated due to poor incorporation of AU-rich triplets (fig. S32), it is clear that 3’-5’ fidelity is significantly reduced (Fig. 6e). This loss of fidelity can now be rationalized in the light of our holoenzyme model as in the 3’-5’ mode (with the triplet triphosphate moiety positioned in the active site) P10 can neither interact with (nor stabilize) the substrate triplet, but instead is positioned to interact with the upstream (3’) primer with no impact on triplet incorporation (fig. S33).

### Evolution of a polymerase ribozyme heterodimer

The structure of the 5TU+t1 TPR comprising a catalytic 5TU and non-catalytic t1 subunit has interesting analogies with proteinaceous polymerases such as the HIV reverse transcriptase (RT) holoenzyme heterodimer formed by a catalytic p65 a non-catalytic p55 (derived from p65). In the heterodimer, p55 supports an extended conformation of p65 that allows positioning the primer/template duplex for optimal processive synthesis (fig. S34). It is tempting to speculate that the non-catalytic t1 RNA subunit may serve a similar function. Indeed, we have demonstrated that t1 helps position J1/3 for optimal interaction with the template and our TPR holoenzyme model indicates that RNA templates of 30 nts (or longer) would be able to interact with t1 P1.

The structure of t1 and the bipartite KL interaction offers a potential explanation for the emergence of the mutualistic interaction between the catalytic and accessory subunits during *in vitro* evolution (10). In the progenitor t1 RNA, the 3’ sequence extension triggered a wholescale reorganization of the tertiary fold, abolishing its catalytic activity. Serendipitously, this enabled a kissing loop interaction, which positioned the t1 5’ selection cassette near to the active site of a catalytically active RNA (5TU progenitor), allowing for mutualistic exploitation of its activity by t1. Over the course of the selection experiment, t1 gained further mutations to better associate and co-evolve with catalytically active subunits, and, in turn, active subunits that could exploit t1 complex formation thrived (10). From the KL mutational study, we further find that the bipartite KL interactions are not only important to facilitate heterodimerization but also to inhibit non-productive homodimerization. Thus, mutualism and eventual molecular symbiosis between the two subunits likely emerged by co-opting a parasitic t1 progenitor RNA.

## Conclusion

Our results describe a first structure and comprehensive structure-function analysis of a polymerase ribozyme, providing a framework for a better molecular understanding of templated RNA-catalyzed RNA synthesis, an enzymatic activity widely considered to be fundamental for the emergence of life’s first genetic system. The cryo-EM structural analysis revealed several novel RNA motifs (such as the Py-turn and A-anchor motifs) and RNA motifs with close resemblance to both engineered and retroviral kissing loop motifs. These discoveries highlight the intricate structural motifs that can be discovered through *in vitro* evolution of large and complex RNA molecules and these may provide input and inspiration for rational RNA nanotechnology designs like the RNA origami architecture (31).

## Data availability

Accession codes (PDB ID 8T2P, EMD-40984).

## Acknowledgements

We thank our colleague K. Nguyen (MRC LMB) for helpful comments on the manuscript. The research at iNANO AU was supported by the Independent Research Fund Denmark (9040-00425B) (EKSM, ESA), the Novo Nordisk Foundation (NNF21OC0070452) (EKSM, ELK, KH, ESA), a fellowship from the Canadian Natural Sciences and Engineering Research Council (532417) (EKSM), a Carlsberg Foundation Research Infrastructure grant (CF20-0635) (ESA), and a Lundbeck fellowship (R250-2017-1502) (ELK). The research at MRC LMB was supported by the Medical Research Council, as part of United Kingdom Research and Innovation (also known as UK Research and Innovation (UKRI)) [MC_U105178804] (CJKW, EG, IG, JFC, JA, PH), a grant from the Volkswagen Foundation (96 755) (EG), a Herchel Smith studentship (2017) (CJKW), a Marie Curie fellowship (H2020-MSCA-IF-2018-845303) (IG), a Carlsberg fellowship (CF17-0809) (ELK). For the purpose of open access, the MRC Laboratory of Molecular Biology has applied a CC BY public copyright license to any Author Accepted Manuscript version arising.

