## Supplementary Information for "Cryo-EM structure and functional landscape of an RNA polymerase ribozyme"

**This PDF file includes:**

Materials and Methods
Figs. S1 to S34
Tables S1 to S4
Captions for Movies S1 to S2

**Other Supplementary Materials for this manuscript include the following:**

Movies S1 to S2

### Materials and Methods

#### RNA preparation

*5TU+t1-anneal sample:*

Ribozyme RNA was prepared essentially as described (1). In brief, RNA was *in vitro* transcribed, gel purified by 10% Urea PAGE, and recovered by freeze and squeeze extraction (removing gel pieces with a Spin-X column (0.22 µm pore size, Costar)). Recovered RNA of 5TU and t1 was then mixed in a 1:1 ratio, precipitated in 96% EtOH+KCl, washed in 70% ice-cold EtOH, and redissolved in buffer (50 mM Tris, pH 8, 100 mM $MgCl_2$) to a final concentration of 3 mg/mL RNA dimer. All buffers and EtOH solutions were filtered (3 K cut-off, Amicon) prior to use. RNA dissolved in buffer was then fast annealed (1 min at 80 °C and then quickly moved to ice) to allow folding of the ribozyme (fast) but not aggregation due to high $MgCl_2$ (slow). Finally, the annealed RNA dimer (3 mg/mL, ~50µM) was added to grids for downstream cryo-EM analysis as described below.

*5TU+t1-cofold sample:*

RNA was prepared under native cotranscriptionally folding conditions essentially as described (2). In brief, RNA was *in vitro* transcribed, purified by size-exclusion chromatography, mixed in a 1:1 ratio (5TU and t1), concentrated and buffer exchanged (to 50 mM Tris, pH 8, 100 mM $MgCl_2$) by spin column filtering (30 K cut-off, Amicon). Finally, the annealed RNA dimer (1.1 mg/mL, ~17 µM) was added to grids for downstream cryo-EM analysis as described below.

*t5+1 sample:*

Ribozyme RNA was prepared similarly as described (1). In brief, RNA was *in vitro* transcribed, gel purified by 10% Urea PAGE, and recovered by electro-elution (Model 422, BIO RAD), and finally filtered with a Spin-X column (0.22 µm pore size, Costar). Ribozyme RNAs were then ethanol precipitated independently and resuspended in milli-pure water to stock concentrations of 50 µM (t5) and 100 µM (t1). t5 and t1 (1:1 ratio; $t5^{+1}$ as heterodimer) were prepared at 10 µM final concentration each in Tris-HCl pH 8.3 50 mM, $MgCl_2$ 25 mM and Tween-20 0.005 % (w/v) as follows: t5 and t1 were mixed and annealed by heating at 50 °C for 5 min, cooled down to 17 °C for 10 min in milli-pure water and finally placed in ice. Then, the required amount of Tris-HCl, $MgCl_2$ and milli-pure water (to reach the final volume minus Tween-20), were added and left to incubate in ice for at least 10 min. Finally, Tween-20 was added before sample application to the grid.

#### Cryo-electron microscopy data acquisition

*5TU+t1-anneal and the 5TU+t1-cofold dataset:*

Protochips 1.2/1.3 300 mesh Au-Flat grids were glow discharged in a GloQube Plus glow discharging system for 45 seconds at 15mA and used immediately after for plunge freezing. Plunge freezing was performed on a Leica GP2 with the sample chamber set to 99% humidity and 15 or 4 degrees Celsius for the annealed and cofolded datasets, respectively. Three microlitres of sample was applied onto the foil side of the grid in the sample chamber before a 4 second delay and then 6 seconds of distance-calibrated foil-side blotting against a double layer of Whatman #1 filter paper. With no delay after blotting the sample was plunged into liquid ethane set to -184 degrees Celsius. All data were acquired at 300 keV on a Titan Krios G3i (Thermo Fisher Scientific) equipped with a K3 camera (Gatan/Ametek) and energy filter operated in

EFTEM mode using a slit width of 20 eV. Data were collected over a defocus range of -0.8 to -2 micrometers with a targeted dose of 60 electrons per square angstrom (Å$^2$). Automated data collection was performed with EPU and the data was saved as gain normalized compressed tiff files with a calibrated pixel size of 0.647 Ångstrom per pixel.

*t5$^{+1}$ dataset:*
Aliquots of 3 µl of pre-annealed t5+1 were applied into C-Flat carbon CF-1.2/1.3 300 mesh grids, which were plasma cleaned for 30 seconds in a 3:1 (Argon:Oxygen) gas mixture. The grids were blotted for 12 seconds at 4 °C and 100% humidity, and plunged into liquid ethane, using a home-made manual plunger. t5$^{1+}$t1 data was collected in a Titan Krios transmission electron microscope operated at 300 kV. Zero-loss-energy images were collected on a Gatan K2-Summit detector in super-resolution counting mode (pixel size of 1.1 Å) with slit width of 20 eV on a GIF Quantum energy filter. Each image was exposed for a total of 18 s (65 electron/Å$^2$) and dose-fractionated into 72 movie frames.

**Single particle image processing and 3D reconstruction**
*5TU+t1-anneal dataset:*
Motion and CTF correction were performed in CS-Live (3, 4) and the micrographs were curated to 12,507 acceptable exposures. Micrographs were binned to a pixel size of 1.294 Å during motion correction. Initial blob picking followed by templated picking using 2D classes generated during CS-live pre-processing were used to generate the initial particle stack. Several rounds of *ab initio* reconstruction followed by heterogeneous refinement were performed before we identified a volume that refined to 8 Å GSFSC (0.143) with 69,977 particles. This volume was used to create 7 different 2D templates that were used to re-initiate templated particle picking.

Templated particle picking, from the templates generated using our *ab initio* model, resulted in 849,824 particle picks that were extracted with a box size of 256 pixels and Fourier cropped to 128 pixels. 2D classification (250 classes) was performed and contained 12 classes (78,962 particles) that were “junk”, and therefore discarded from further analysis. *Ab initio* reconstruction using 30,000 particles and 3 classes was used to generate initial volumes. Heterogeneous refinement of 770,862 particles resulted in 306,161 particles being sorted into the class that was further investigated. These particles were re-extracted with newly aligned shifts for the particle centers and subjected to another 3-class *ab initio*, this time using the entire particle stack (302,927 particles), which was then followed by heterogeneous refinement into three volumes. 126,690 particles were sorted into the best class and used for a 2-class *ab initio* job with class similarity parameter set to 0, followed by heterogeneous refinement into those two volumes, resulting in 95,114 particles being kept. The method of 2 class *ab initio* with a class similarity of 0, followed by heterogeneous refinement was repeated twice more, further reducing the particle stack to 71,746 and then to 45,589. At this point we were no longer removing junk particles but sorting different structural classes of the heterodimer that either had or did not have good alignment with the epsilon domain (P9/P10). Heterogeneous refinement into the previous 2 *ab initio* volumes was performed 3 times, reducing the particle stack to 35,633, then 31,103 then 28,000 particles. Further classification was determined to have diminishing returns in terms of improved resolution.

It was determined that we were able to achieve better particle alignments with a smaller box size and so the particles were re-extracted with a box size of 208 and not Fourier cropped. Some close particle picks were found to be preventing the FSC curve from dropping to zero and were removed, resulting in a final particle stack of 26,167 particles. A final local refinement using a full mask was performed while minimizing over per-particle scale at each iteration of the refinement (5). This resulted in a GSFSC (0.143) of 5.9 Å. The final map was sharpened with a B-factor of 275. Masks were created to locally refine the 5TU core and t1 turn motifs independently. Masks were created in ChimeraX by using the command molmap sel 15 while selecting residues from the PDB model inside the area we were refining. This map was further dilated and soft-padded (with a width of 15) in cryoSPARC's volume tools. The best alignments were attained by extending the search range to 40 Å and minimizing over the per-particle scale factor. The half maps from these local refinements were sharpened using the software DeepEMhancer (6).

*5TU+t1-cofold dataset:*

Motion and CTF correction were performed in CS-Live (3, 4) and the micrographs were curated to 14,906 acceptable exposures. Micrographs were binned to a pixel size of 1.294 Å during motion correction. Initial blob picking, 2D classification and ab-initio reconstruction generated during CS-live pre-processing were used to generate the initial particle stack leading to a volume that refined to 8.3 Å GSFSC (0.143) with 28.596 particles. Templated particle picking, from the templates generated using the initial refined volume, resulted in 516,889 particle picks that were extracted with a box size of 265 pixels and Fourier cropped to 128 pixels. 2D classification (50 classes) was performed containing 14 classes (105,116 particles) that were "junk", and therefore discarded from further analysis. Two rounds of ab-initio reconstruction with 3 classes and 2 classes, followed by heterogenous refinement after each ab-initio job using the ab-initio created volumes, were performed to sort particles, resulting in a stack of 144,118 particles, which were re-extracted with newly aligned shifts for the particle centres and no Fourier cropping. Then particle stack was subjected to another three 3-class *ab initio* reconstructions, each time taking the two best classes, discarding the works from the analysis (resulting in 99,993, 92,455, and finally 86,545 particles respectively). The final stack of 86,545 particles was non-uniformly refined with an initial lowpass resolution of 20 Å followed by local refinement using a full mask resulting in the reported GSFSC (0.143) of 5.0 Å. Masks were created to locally refine the 5TU core and t1 turn motifs independently. Masks were created in ChimeraX by using the command molmap sel 15 while selecting residues from the PDB model inside the area we were refining. This map was further dilated and soft-padded (with a width of 15) in cryoSPARC's volume tools. The best alignments were attained by extending the search range to 40 Å and minimizing over the per-particle scale factor. The half maps from the local refinements were sharpened using the software DeepEMhancer (6).

*$t5^{+1}$ dataset:*

We collected 452 movies which were imported to cryoSPARC. Whole-image drift correction of the movie frames ('Patch motion correction'), and contrast transfer function (CTF) estimation ('Patch CTF estimation') were performed using default parameters. An initial stack of ~ 37k particles were "blob picked" and extracted using a 256-pixel box (binning 4) and subjected to one round of reference-free 2D classification (150 classes). From these, 14 2D classes where selected to conduct a template-based particle picking. A total of 38276 particles were extracted using a 256-pixel box (binning 2). These particles were then used to produce 3 *ab initio* models in which one of them already had the overall structure of t5+1 at low resolution. To remove the defective particles, an initial 3D classification ('Heterogeneous refinement' in cryoSPARC) was performed using all the particles and the 3 *ab initio* models, where the 2 ill-formed models were acting as ‘sinks’. The ‘Heterogeneous refinement’ was repeated twice by using the particles of the best volume and the 3 volumes of the previous job as input. Finally, the particles (n = 5,485) of the best volume, were extracted from the micrographs using a 256-pixel box, and an ‘homogeneous refinement’ was performed giving rise to an EM-map with an overall 8.0 Å GSFSC resolution. 5TU+t1 model and EM-map from $t5^{1+}t1$ were then docked using Chimera.

**Model building**

To determine the helix placement for 5TU and t1 we used DRRAFTER (7, 8) to fit the helical fragments into our map. For 5TU we used the secondary structure diagram from Attwater et al. (1) as restraints for DRRAFTER. Taking into account that we had knowledge of a potential interaction / dimerization point between the 5’ cap region from 5TU and t1, we manually placed the 5’cap helix at the end of the region of our map— which was clearly the long single stranded 5TU:J1/3 (Fig. S7a). Using this initial helix placement, DRRAFTER was able to determine the helix positions and model the single-stranded junctions in an automated fashion. After the first round of modelling, which produced 3000 models, the top 10 models only converged to a mean pairwise RMSD of 22.3 angstrom (Fig. S7a). Upon visual inspection, most of these top models had clearly failed to fit all helical components within the volume of our map. However, the best fitted model had managed to place all 152 nucleotides within the volume reasonably well (Fig. S7a). We then used this model as a starting point for manual model building of the heterodimer.

Model building with DRRAFTER failed to produce reasonable models using the secondary structure diagram for t1, as described in Attwater et al. (1). Furthermore, it was clear from the remaining unfilled space in our map that there was an extended helical component that was longer than any of the helical components previously predicted. We used the NUPACK web application (9) to predict the secondary structure of t1 (Fig. S5c), and used this predicted structure as restraint for model building with DRRAFTER. The end of the extended helix predicted by NUPACK was manually fit in the map (Fig. S7b, column 1) and after only 2000 models generated by DRRAFTER, the top 10 models had achieved a mean pairwise RMSD of 12.2 angstrom (Fig. S7b, column 2). The top model (Fig. S7b, column 3) was selected as a template for manual model building of the heterodimer.

t1 RNA has 2 major helical components (t1:P1 and t1:P3), which are connected by a joining region (t1:J1/2), the short helix (t1:P2) and another joining region (t1:J2/3). t1:J1/2 has a 7-nucleotide loop component that our DRRAFTER model places where the 5’ P1 cap helix of 5TU is located. This 7-nucleotide loop has 6 bases which complements perfectly with the 5TU hairpin loop of the 5’ cap. We manually built this dimerization site around an idealized 6-bp double stranded helix between sequences U6-U7-C8-U9-C10-G11 from 5TU and C22-G23-A24-G25-A26-A27 from t1.

After having defined the coarse features of the 5TU+t1 dimer, we started to focus on the fine details in our model. The two GAUA loop sequences make up the second dimerization site between 5TU and t1 forming a 2-base pair kissing loop. This is reminiscent of the 2-base pair GACG kissing loop from the 5’-end dimerization signal of the Moloney murine leukemia virus (MoMuLV) RNA (10). Accordingly, we used the NMR structure (PDB: 1F5U) of this dimerization signal as a template to model this interaction site. We remodelled 5TU:J2/3 and 5TU:P6 using the crystal structure of the class I ligase ribozyme (PDB: 3IVK) (11) as a template. Finally, individual DRRAFTER runs with 7600 models were setup to rebuild t1:J3/2 (SI Fig. 6e), 5TU:J3/4 (Fig. S7c) as well as 5TU:J10/9 (Fig. S7d). These DRRAFTER runs on smaller fragments reached much better convergence than the DRRAFTER build of the entire RNA strands, with mean pairwise RMSD values of 1.7, 2.1 and 2.4 Å, respectively.

Flexible fitting with molecular dynamics, as well as general model inspection and combination was performed using ISOLDE (12) and ChimeraX (13, 14). The PDB-tools software package was utilized for renumbering, editing the sequence and merging chains from PDB models (15). The model was iteratively refined using Real-Space refinement and validation using Phenix software package (16, 17) and energy minimizations using QRNAS (18), which uses the AMBER force fields (19, 20). Validation (21) of the final model can be found in Fig. S7 and Table S1.

**3D variability analysis**

3DVA was performed in cryoSPARC (22) using the 126,690 particle from the 5TU+t1 (anneal) dataset stack as an input, which was only 3D classified to remove junk particles. A filter threshold of 8 Å was applied and 3 components were solved. The second component contained the greatest motion and is the only one presented herein. The 3DVA intermediates display job was used to sort the particles into 9 classes with no overlap (top hat windows). Three classes from each flank of the distribution were used to reconstruct volumes without alignment, followed by homogeneous and then local refinements. The leading-edge refinement contained 19,854 particles and the tailing edge refinement contained 17,799 particles. To generate the movie, the consensus model was fit into each map, individually and molecular dynamics with flexible fitting was performed using ISOLDE. The force field strength was reduced to 0.05 x 1000 kJ per mol per map units by cubic angstrom and allowed to run for 15 minutes with a 0-degree temperature factor. This allowed the model to smoothly drift into the slightly deviant conformations with minimal change to the secondary structure. The coordinate sets between the two states were then calculated in ChimeraX using the default corkscrew rigid-body transformation morph command.

**Selection library synthesis**

A library containing all possible single mutants, insertions and deletions in t5 was synthesised by a commercial supplier (Twist Bioscience). The library (0.5 ng) was used as template in a 50 µl GoTaq HotStart (Promega) PCR using forceGG and HDVrec primers. The PCR product (0.1 ng) was further mutagenised in a 50 µl error-prone PCR using GeneMorph II Random Mutagenesis Kit (Agilent) for 30 cycles using forceGG and HDVrec primers, following the manufacturer's instructions. The resulting amplicon was purified using agarose gel, and further amplified in a 50 µl GoTaq HotStart (Promega) PCR using HDVRT and t5_tri12x12 primers. The DNA from this reaction was transcribed into RNA overnight using MEGAshortscript™ T7 Transcription Kit (ThermoFisher); products of the transcription was subsequently purified using preparative-scale urea-PAGE.

***In vitro* evolution cycle**

The t5 library selection construct was annealed with equimolar 5' biotinylated primer and t1 ribozyme, as well as triplets in water (80 °C 2–4 min, 17 °C 10 min). Chilled extension buffer was added, and the reaction was then frozen and incubated at −7 °C. After the incubation the reaction was thawed on ice. Constructs were then precipitated with 0.3 M sodium acetate in isopropanol (55%) before treatment with polynucleotide kinase (NEB) followed by denaturation to resolve the HDV-derived 2', 3'-cyclic phosphates and allow later adaptor ligation.

Constructs were urea-PAGE separated alongside FITC-labelled RNA markers equivalent to successfully ligated constructs. The marker-adjacent gel region in the construct lane was excised. Biotinylated (primer-linked) constructs were eluted overnight into BB with 100 µg MyOne C1 beads. After 30 µm filtering (Partec Celltrics(Wolflabs (York, UK))) of the supernatant to remove gel fragments, the beads were washed in BB then 0.1 M NaOH to confirm covalent linkage of construct to primer, before further BB washing and transfer to a fresh microcentrifuge tube to minimize downstream contamination. 3' adaptors were subsequently ligated to bead-bound constructs for 2 hr (with buffer/enzyme added after bead resuspension in other reaction components including 0.04% Tween-20). Beads were BB washed.

Bead-bound constructs were put into a 50 µl RT-PCR using HDVRec and forceGG primers, and SuperScript III One-Step RT-PCR System with Platinum Taq DNA Polymerase (ThermoFisher). Resulting products were further amplified using HDVRT and t5_tri12x12 primers, in a 50 µl GoTaq HotStart (Promega) PCR to generate the DNA template for the subsequent round of selection. The DNA was subsequently transcribed into RNA overnight using MEGAshortscript™ T7 Transcription Kit (ThermoFisher); products of the transcription was subsequently purified using preparative-scale urea-PAGE. At the end of the selection, t5 libraries were amplified by P51HDVba and P7forceGG primers in a 50 µl GoTaq HotStart (Promega) PCR; products were purified by agarose gel, quantified using a KAPA SYBR FAST qPCR kit (KAPA Systems) and then sequenced on a HiSeq 2500 (Illumina).

**Comparing t5 and 5TU TPR activity**

0.5 µM of t5 or 5TU was mixed with 0.5 µM of t6F10mix RNA template, 0.5 µM of F10 primer, 5 µM each of $^{ppp}$GCG, $^{ppp}$ACC, $^{ppp}$UUC, $^{ppp}$GAA, $^{ppp}$CGC, $^{ppp}$AUA, $^{ppp}$GGU, $^{ppp}$CCA, with or without 0.5 µM of t1 (as described within SI Fig 1C), made up to 3.75 µL in water. Reactions were annealed to 80 °C for 2 minutes, followed by 17 °C for 10 minutes. Reaction buffer was added to a final concentration of 200 mM $MgCl_2$ and 50 mM Tris pH 8.4; reactions were subsequently frozen in dry ice, and incubated at -7 °C for 12 hours. Reactions were then thawed, to which 50 pmol of a competing oligo (t6F10mix-comp) added. Products of the reaction were separated on analytical urea-PAGE, and detected on a Typhoon Trio scanner (GE Healthcare (GE) (Chicago, USA)).

**Determination of 5TU+t1 adaptive landscape**

For construction of the 5TU library, 5TU-F1, 5TU-F2, 5TU-R1 and 5TU-R2 were mixed equimolar, and a small amount (around 0.05 pmol of each) is used as a template for a 50 µl PCR using Q5 DNA Polymerase (NEB), with forceGG and HDVrec as primers. For construction of the t1 library, t1-F1 and t1-R1 were mixed equimolar, and a small amount (around 0.05 pmol of each) is used as a template for a 50 µl PCR using Q5 DNA Polymerase, with t1rec and HDVrec as primers. For both subunits, the amplification products were purified and diluted, such that 106 molecules were subsequently used as templates in a PCR using Q5 DNA Polymerase (primers 6AUA-6AACA-fGG and HDVRT for 5TU, and 6AUA-6AACA-t1rec and HDVRT for t1). The resulting PCR products were transcribed using using MEGAshortscript T7 Transcription Kit (ThermoFisher); products of the transcription were subsequently purified using preparative-scale urea-PAGE.

Both 5TU and t1 libraries were subjected to 1 round of *in vitro* evolution in triplicates, as described above. Pre-selection and post-selection libraries were amplified using primers that introduce indexed adaptors for Illumina sequencing (P51HDVba, P52HDVba, P53HDVba and

P510HDVba for the forward primer and P7forceGG for the reverse primer for the 5TU libraries, and P51t1rec, P52t1rec, P53t1rec and P54t1rec for the reverse primer and P7HDVba for the reverse primer for the t1 libraries). PCR products were subsequently quantified using a KAPA SYBR FAST qPCR kit (KAPA Systems) and then sequenced on a HiSeq 2500 (Illumina).

**Calculating fitness associated with each genotype**

Reads from the HiSeq run were merged using PEAR (23) and demultiplexed into their respective libraries (input and 3 output libraries for both 5TU and t1) using a custom Python script, according to 6-nucleotide barcodes at the 5' end of each read. Using FASTX-toolkit, reads were trimmed to only contain the 5TU or t1 gene, and quality filtered such that each read contains only bases with Q-score 30 or above. Remaining reads were aligned to the wild-type 5TU or t1 sequences and mutations were called using alignparse (24). Genotypes containing 10 reads or more in the input libraries, as well as at least 1 read in each of the 3 output libraries, were retained for downstream analysis; the rest of the genotypes were discarded, as their fitness could not be accurately calculated.

To calculate the fitness of each genotype, as well as the error of fitness measurement, we took into account sampling error associated with a given read count (25). We first calculated the fraction of each library occupied by each genotype in the input ($f_{in}$) and 3 output ($f_{out}$):

$$f_{in_g} = \frac{counts_{in_g}}{\sum_{g=1}^{L} counts_{in_g}}$$

$$f_{out_{gi}} = \frac{counts_{out_{gi}}}{\sum_{g=1}^{L} counts_{out_{gi}}}$$

where g is the genotype in question (from 1 to L where L is the total number of genotypes), and i is the output replicate (1, 2, or 3).

Each input and output frequency is modelled with a Poisson variance (σ) associated with the number of reads for that genotype and the total number of reads in that library:

$$\sigma_{in_g} = \sqrt{\frac{1}{counts_{in_g}} + \frac{1}{\sum_{g=1}^{L} counts_{in_g}}}$$

$$\sigma_{out_{gi}} = \sqrt{\frac{1}{counts_{out_{gi}}} + \frac{1}{\sum_{g=1}^{L} counts_{out_{gi}}}}$$

For each genotype, we merged the 3 output fractions as an average, weighted by the inverse of the variance of the genotype:

$$f_{out_g} = \frac{\sum_{i=1}^{3} f_{out_{gi}} * \sigma_{out_{gi}}^{-2}}{\sum_{i=1}^{3} \sigma_{out_{gi}}^{-2}}$$

The associated error is:

$$\sigma_{out_g} = \sqrt{\frac{1}{\sum_{i=1}^{3} \sigma_{out_{gi}}^{-2}}}$$

We calculated the fitness (F) of each genotype as the log2 ratio of the enrichment of the genotype during selection, and the enrichment of the wild-type sequence during selection:

$$F_g = log_2\left(\frac{f_{out_g}/f_{in_g}}{f_{out_{WT}}/f_{in_{WT}}}\right)$$

The associated error with this fitness is:

$$\sigma_{F_g} = \frac{1}{ln(2)} * \sqrt{(\sigma_{out_g})^2 + (\sigma_{in_g})^2}$$

Due to normalisation to wild-type and subsequent transformation into log2 space, wild-type sequences would have a fitness of 0, while less or more functional mutants would have fitness <0 or >0, respectively.

For a given double mutant consisting of point mutations A and B, we defined epistasis (Ɛ) as:

$$\mathcal{E} = F_{AB} - F_A - F_B$$

To check whether a given epistasis value was significant, we performed a one-sample t-test using Ɛ and its propagated error. For double mutants within 5TU and t1, the false discovery rate was adjusted using the Benjamini–Hochberg method.(26)

**Fidelity assay for substrate lengths**

Reactions were carried out with 0.1 µM ribozyme and 0.5 µM template (TγGA2SU) and 5' FITC-labelled primer (Fs5γ7) in 200 mM $MgCl_2$, 50 mM tris•HCl pH 8.3. As described(1), ribozyme and template/primer/substrate mixes were pre-annealed in water independently (80 °C 2 min, 17 °C 10 min) before buffer addition and freezing on dry ice (10 min) followed by incubation at -7 °C for 16 hours. Substrates were at 5 µM each (triplets), 2 µM each (tetramers), 1 uµM each (pentamers) and 0.5 µM each (hexamers). $^{HO}$CUG was used as the +2 downstream triplet, to prevent incorporation of the +2 triplet to the reaction products to simplify analysis. Reactions were stopped by addition to 1 µl 0.5 M EDTA (pH 7.4) after thawing. Samples were denatured in 66.6 mM EDTA (pH 7.4), 6 M urea (94 °C 5 min), before separation on a 35 cm 30% 19:1 acrylamide:bis-acrylamide 3 M urea tris-borate gel. This is sufficient to separate correct and incorrect products due to the differential migration rates of adenine vs guanine bases. FITC fluorescence were detected using a Typhoon trio scanner (GE) and quantitated using ImageQuant software (GE).

**FidelitySeq assay**

FidelitySeq assay reactions were carried out using 4 pmol template (UP1NNN), 2 pmol $t5^{+1}$, 40 pmol $^{ppp}$NNN in 8 µl reaction. Reactions were stopped after 48 hr and separated by urea-PAGE. Empty template and bands resulting from triplet incorporation were excised, eluted in 10 mM tris•HCl pH 7.4 overnight, precipitated in 73% ethanol with 1.5 µl 1 % glycogen carrier, and resuspended in water. These products were annealed to 1 µM BiouploopRT primer in 26.8 µl water (72 °C 3 min, ice 3 min) and reverse transcribed (40 µl Superscript IV reaction supplemented with 0.02% Tween-20, 30 min 65 °C).

RT products were bound to MyOne C1 (Invitrogen) streptavidin-coated paramagnetic microbeads (8 µl thrice-BWBT washed beads per reaction) in 160 µl BWBT (200 mM NaCl, 10 mM tris•HCl pH 7.4, 1 mM EDTA, 0.1 % Tween-20) supplemented with 10 mM EDTA for 30 min. Beads were washed in BWBT, incubated for 1 min in 25 mM NaOH, 1 mM EDTA, 0.05 % Tween-20 to denature any duplex(27), washed again in BWBT before adaptor ligation (10 µl App DNA/RNA ligase reaction: 1X NEB buffer 1, 5 mM $MnCl_2$, 0.04 % Tween-20, 2 µM adenylated-HDVlig adaptor, 65 °C, 2 hr). Beads were washed twice in BWBT before biotinylated species were eluted by heating in 95 % formamide, 10 mM EDTA (94 °C, 5 min) and separated by urea-PAGE.

Ligation products were excised and eluted into BWBT before binding to 3 µl thrice-washed beads per reaction and resuspension in 10 µl water. Samples were PCR amplified with GoTaq HotStart master mix (Promega) (0.5 µM primers P3HDV and barcoded P5Xuplooprt, 40 °C anneal, 25 cycles), agarose gel purified and pooled for sequencing (Illumina HiSeq).

Sequencing reads were split by barcode (yielding over 250,000 reads per sample), and 3' adaptor sequences trimmed off using the Galaxy web platform, at the public server usegalaxy.org (28). Custom python code was used to count the number and type of incorrect and correct incorporation events per template, correcting for cross-template priming (which was negligible). Base position fidelity and overall extension fidelity were calculated as geometric means as described (1). Extension likelihood was calculated by dividing extended count by total reads, normalised by fractional gel intensities of the extended and unextended bands. Likelihood of correct extension was calculated by multiplying a given triplet's extension likelihood by its fidelity.

**Oligonucleotide syntheses**

RNA templates for the substrate length and FIdelitySeq assays were prepared by *in vitro* transcription and urea-PAGE purification. These were transcribed as described (1), using MegaShortScript enzyme and buffer (ThermoFisher), from dsDNA templates. Triplets and substrates up to hexamers in length were synthesised and purified as described (1). Briefly, T7 RNA polymerase transcription of short overhanging templates produced triplets and dinucleotides (30 µl reactions, 72 nmol each NTP required, 15 pmol template, 37 °C overnight). Products were separated by urea-PAGE, identified by UV shadowing and relative migration rates. Correct products were eluted and precipitated in 85% ethanol.RNA templates for the substrate length and FIdelitySeq assays were prepared by *in vitro* transcription and urea-PAGE purification. These were transcribed as described (1), using MegaShortScript enzyme and buffer (ThermoFisher), from dsDNA templates. Triplets and substrates up to hexamers in length were synthesised and purified as described (1). Briefly, T7 RNA polymerase transcription of short overhanging templates produced triplets and dinucleotides (30 µl reactions, 72 nmol each NTP required, 15 pmol template, 37 °C overnight). Products were separated by urea-PAGE, identified by UV shadowing and relative migration rates. Correct products were eluted and precipitated in 85% ethanol.

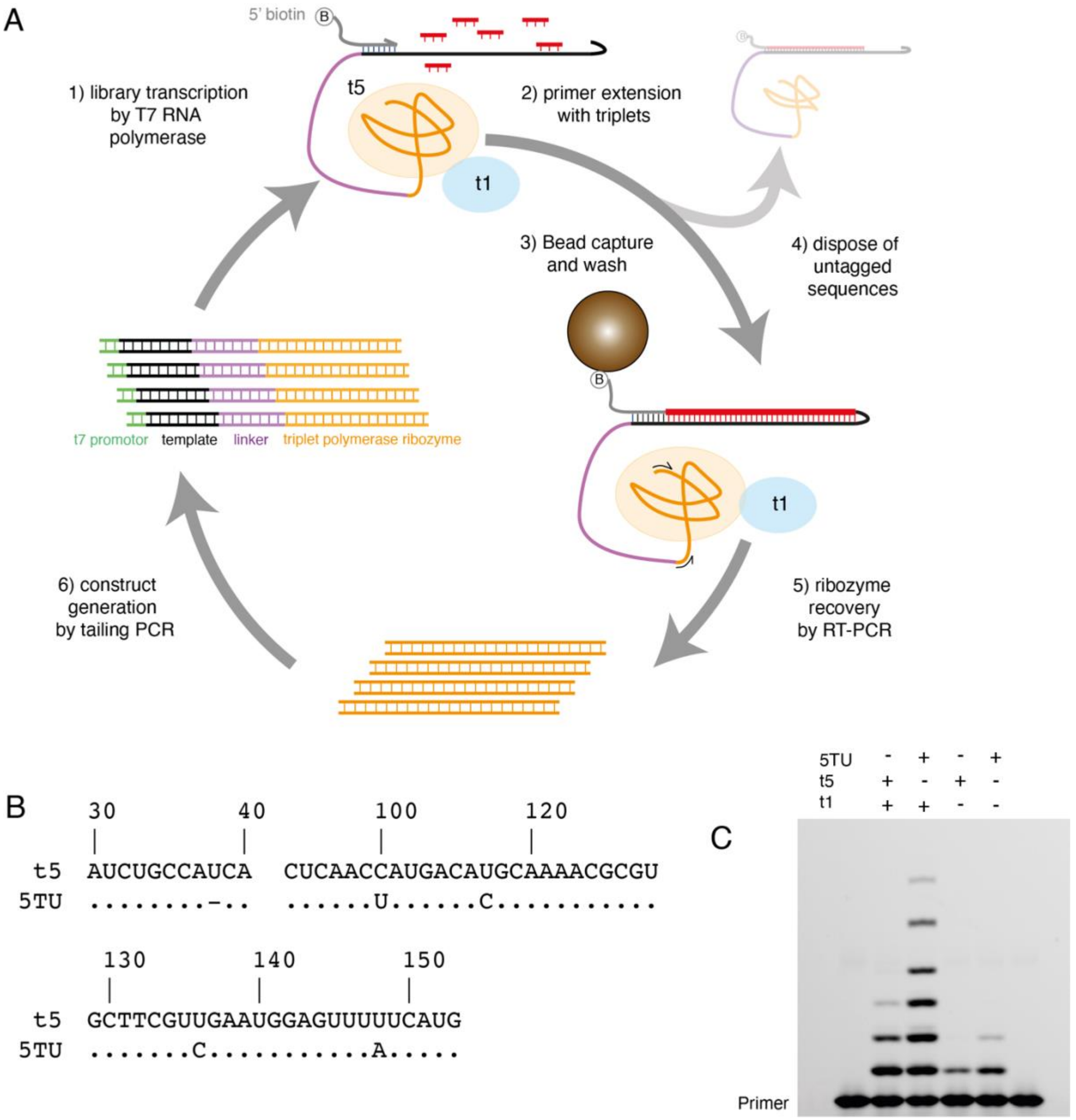

**Fig. S1. TPR evolution.**

(**a**) Diagram of the TPR activity selection strategy. Ribozyme sequences are coloured in orange, linkers in purple, biotinylated primers in grey, templates in black, streptavidin-coated magnetic bead in brown. B - biotin. (**b**) Multiple sequence alignment of t5 against variant 5TU. Sequences of the ribozymes outside of the displayed regions are identical to t5. Numbering corresponds to positions in t5. **c**) Activity of 5TU alone or in combination with wild-type t1 or top selected variant t1.5 on mixed sequence template, encoding for successive incorporations of GCG, ACC, UUC, GAA, CGC, AUA, GGU and CCA.

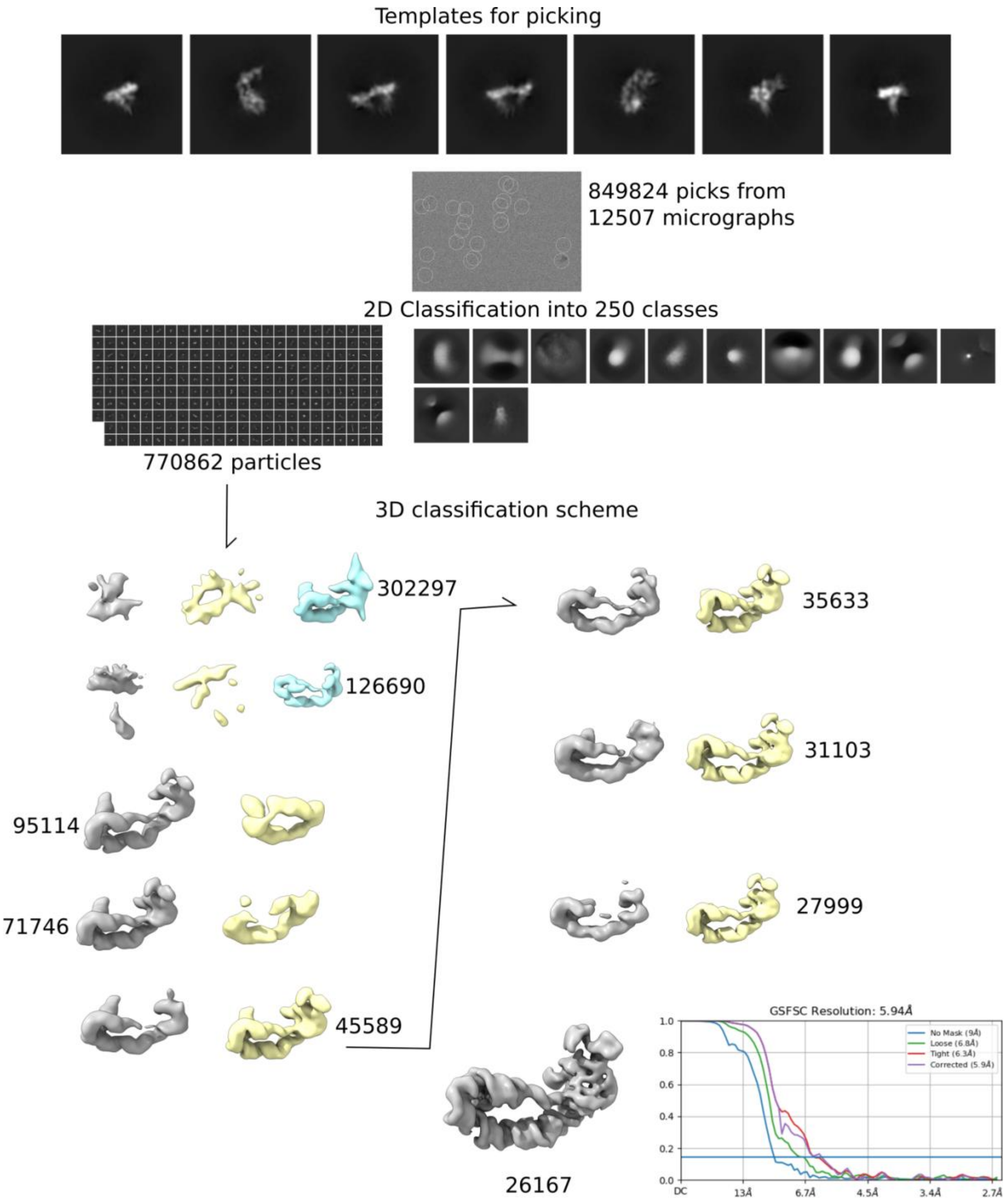

**Fig. S2. Cryo-EM single particle analysis workflow for 5TU+t1-anneal sample.** Numbers listed in the 3D classification scheme are the number of particles left in the class they are adjacent to and that were used for the subsequent round of classification.

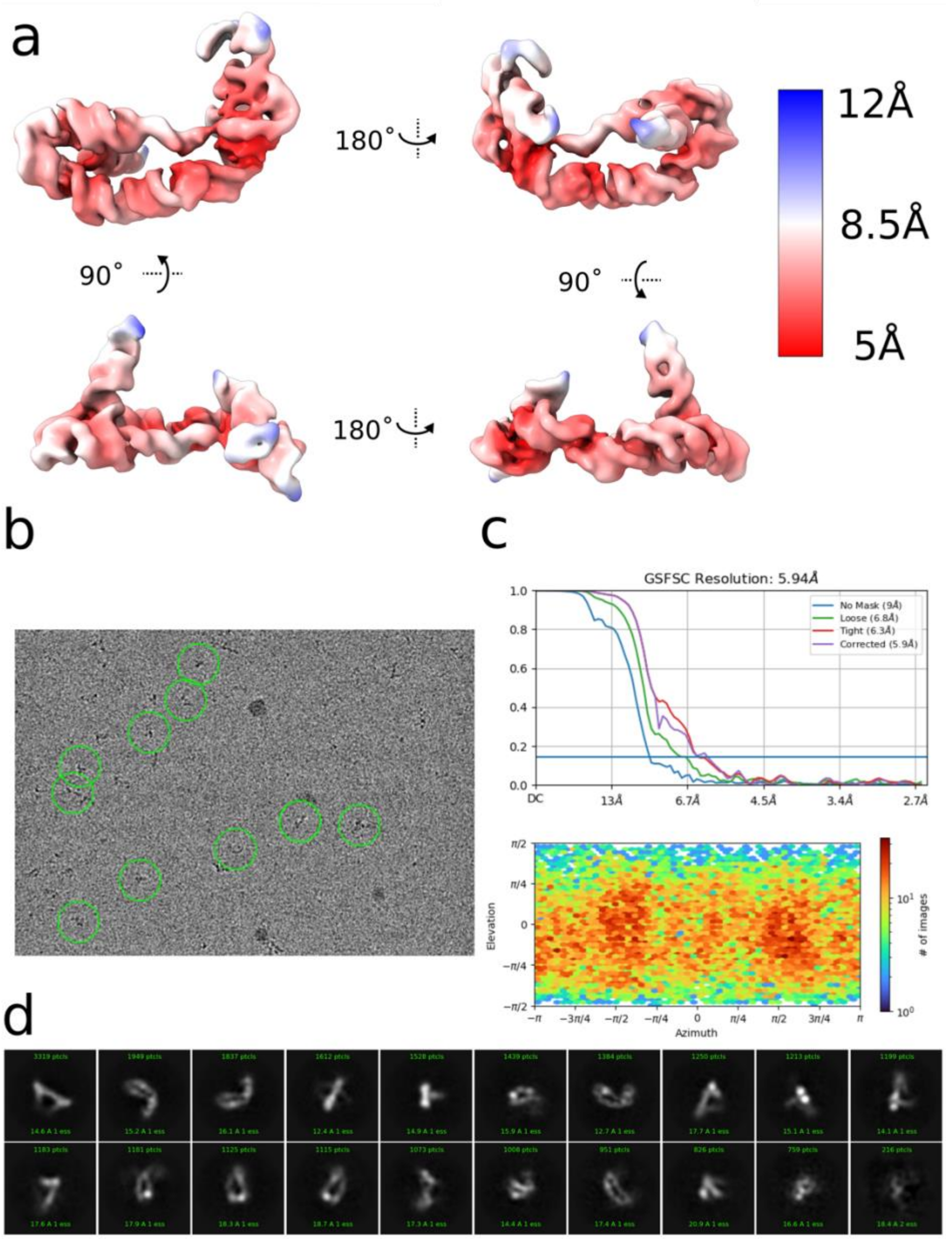

**Fig. S3. Features of the final particle stack for the 5TU+t1-anneal sample.**
(**a**) Local resolution mapped onto the locally refined 5TU+t1 heterodimer cryo-EM map by color. (**b**) Exemplary micrograph showing 10 particle picks were retained in the final particle stack. (**c**) Gold-Standard FSC curve and viewing angle distribution from the final local refinement run in cryoSPARC. (**d**) 20 2D class averages generated from the final particle stack.

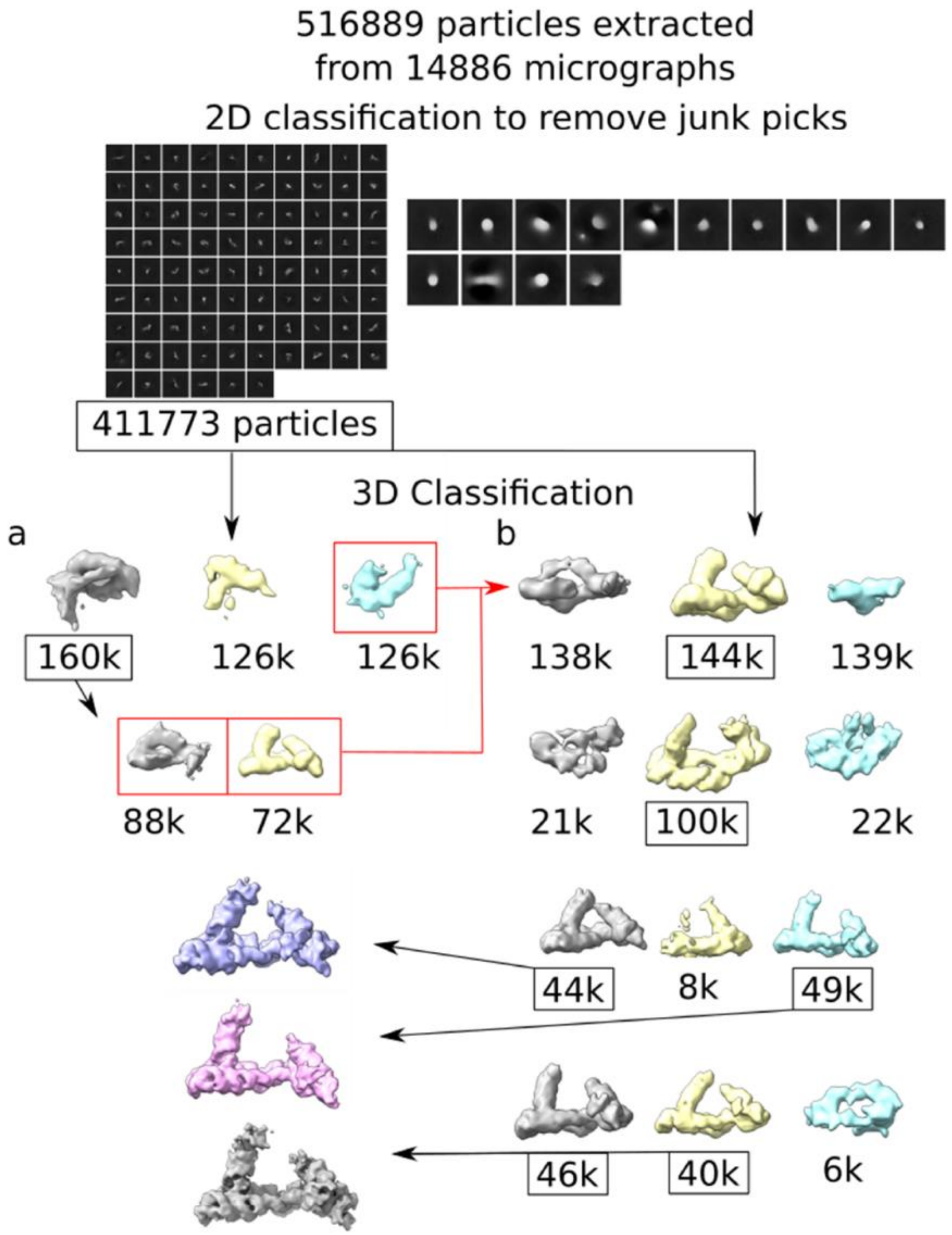

**Fig. S4. Cryo-EM single particle analysis workflow for the 5TU+t1-cofold sample.**
Numbers listed in the 3D classification scheme are the number of particles left in the class they are adjacent to and that were used for the subsequent round of classification. All 3D classification was performed as a 3-class *ab initio* reconstruction job, with the exception of the first job in column **b** where we used the "good" volume and two "junk" volumes from column **a** as inputs for 3D heterogeneous refinement with the full particle stack after 2D classification.

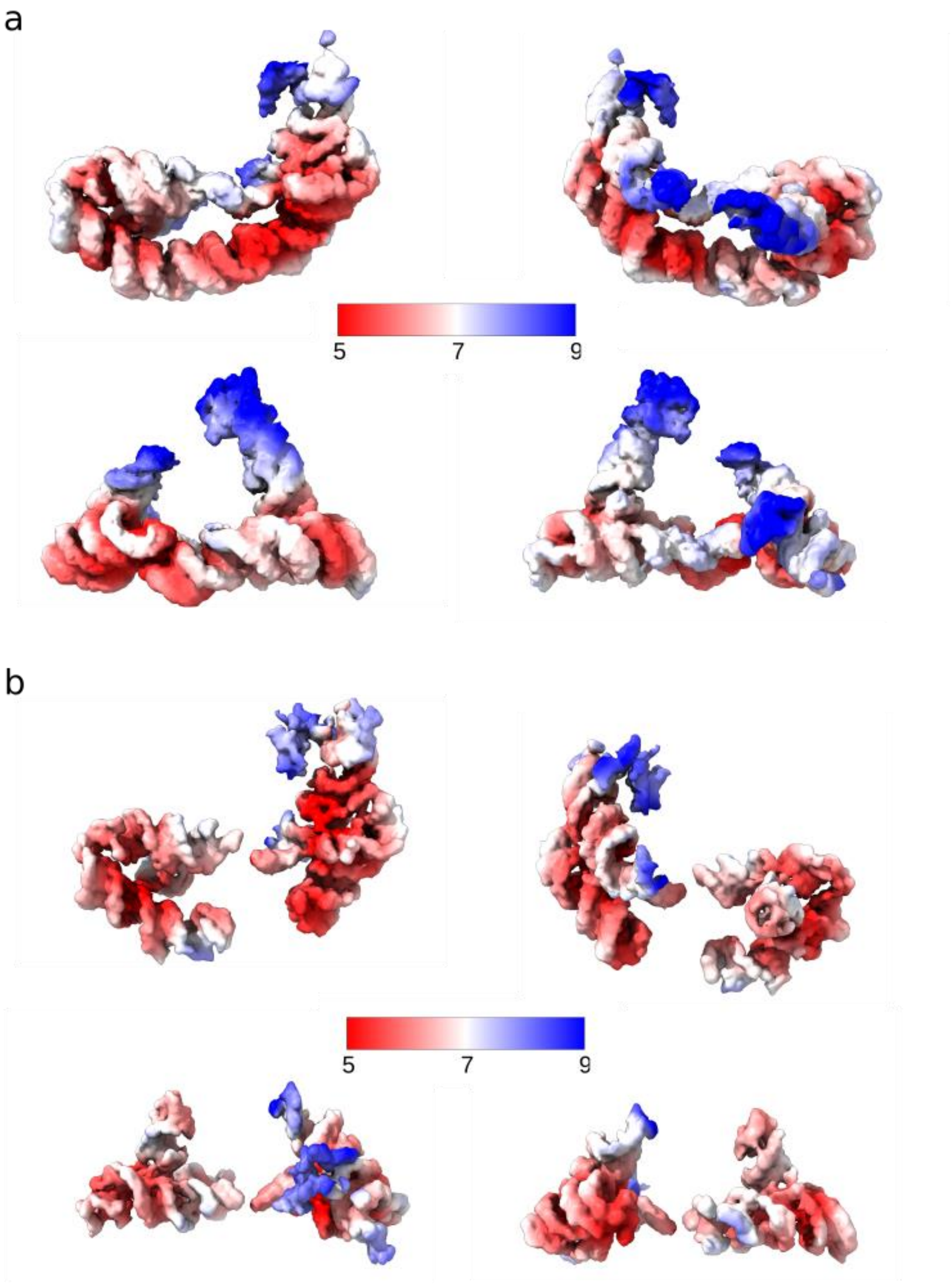

**Fig. S5. Local resolution of the final sharpened maps for the 5TU+t1-cofold sample.** Local resolution mapped as surface color onto the deepEMhancer sharpened 5TU+t1 heterodimer cryo-EM maps from the consensus refinement **(a)** and two individually locally refined regions **(b)**.

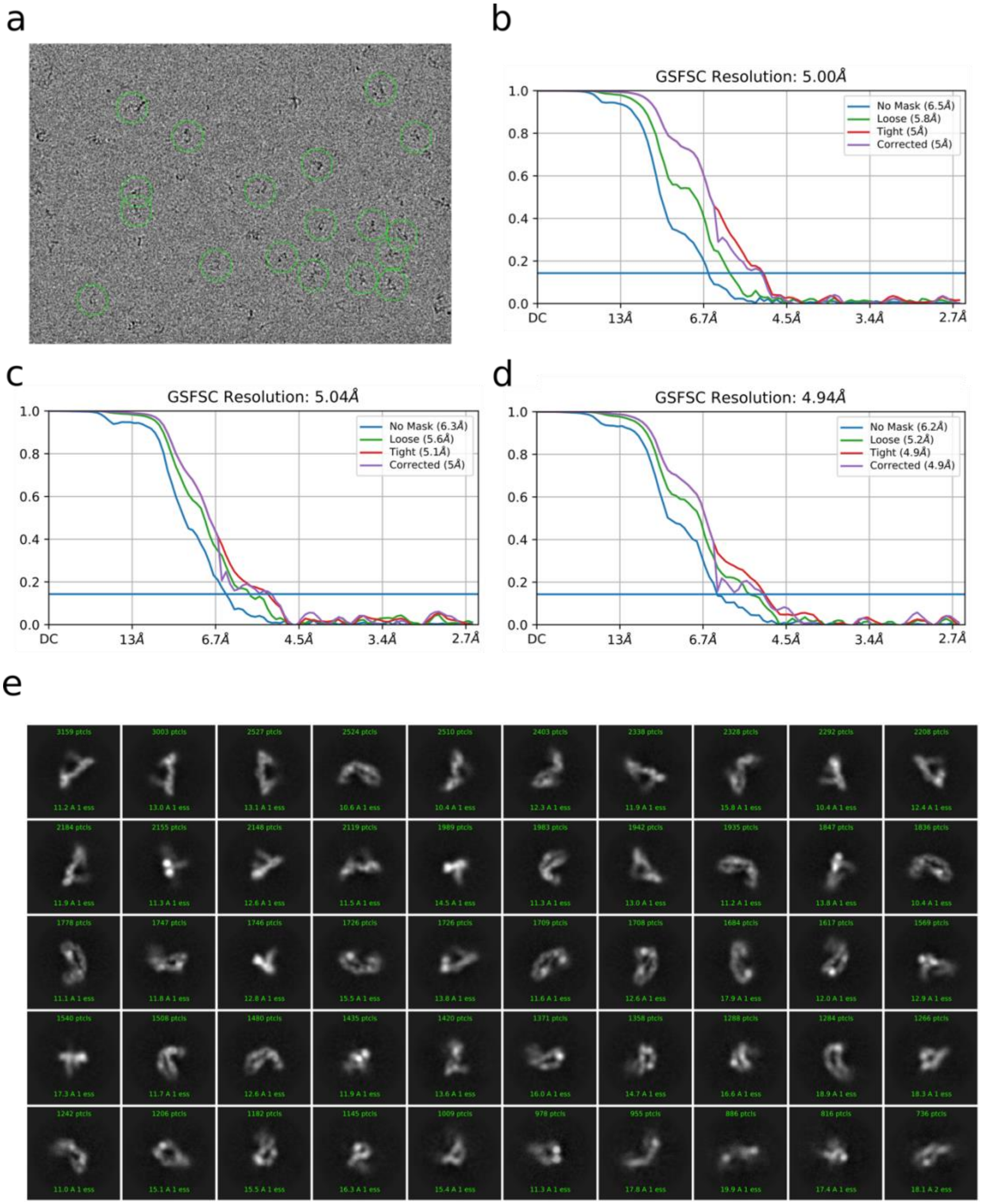

**Fig. S6. Reconstruction validation for the 5TU+t1-cofold sample.**

(**a**) An example micrograph is shown depicting the picked particles that remained in the final particle stack. The GSFSC curves are shown for the consensus refinement (**b**), and the masked local refinements of the t1 turn (**c**) and 5TU core (**d**). 2D classes generated from the final particle stack.

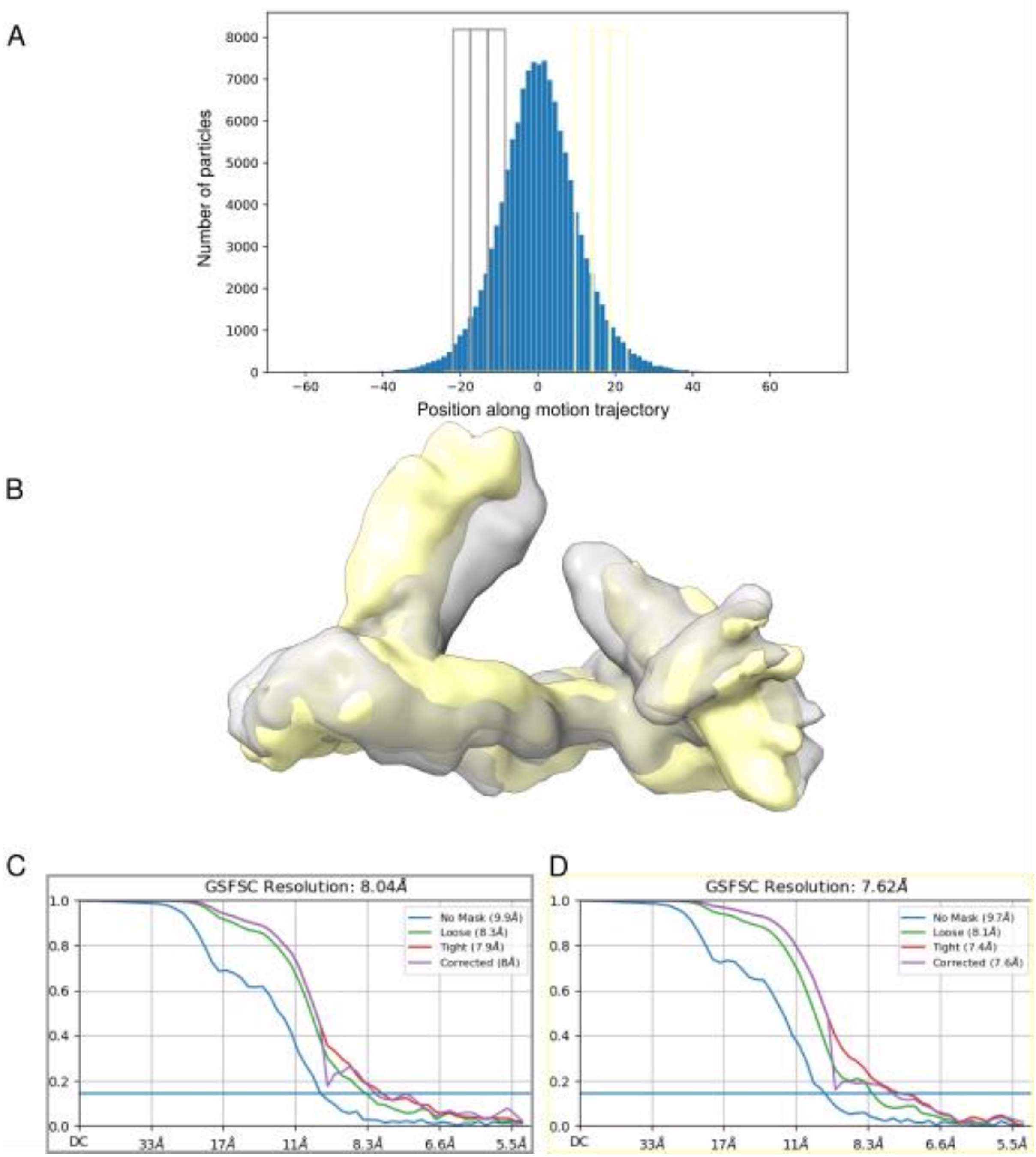

**Fig. S7. Flexibility of TPR structure for the 5TU+t1-anneal sample.**

3D variability analysis of the TPR from the 126,690-particle stack shown in Fig. S2. The distribution of particles along the motion trajectory in (**a**) and the coloured boxes represent the particles used for the independent reconstruction and refinement of the volumes in (**b**). Gold-standard Fourier Shell Correlation curves for the reconstructed volumes are shown in (**c**) & (**d**).

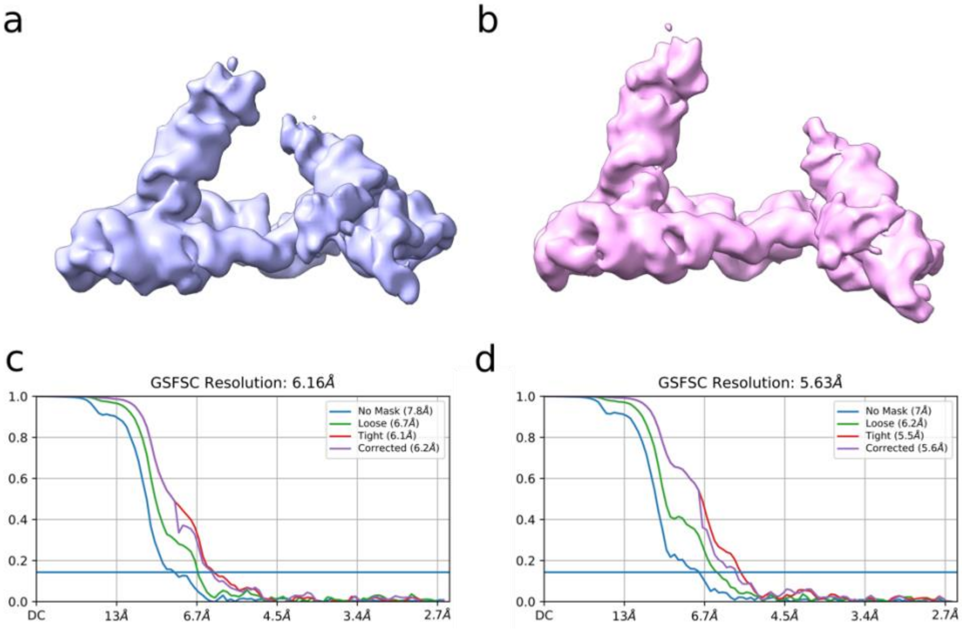

**Fig. S8. Sub-populations of the t1-P1 arm isolated during 3D classification for the 5TU+t1-cofold sample.**

The 44k **(a)** and 49k **(b)** classes from Figure S2 were subjected to non-uniform refinement and local refinement resulting in two distinct conformers whose GSFSC curves are shown in **(c)** and **(d)**.

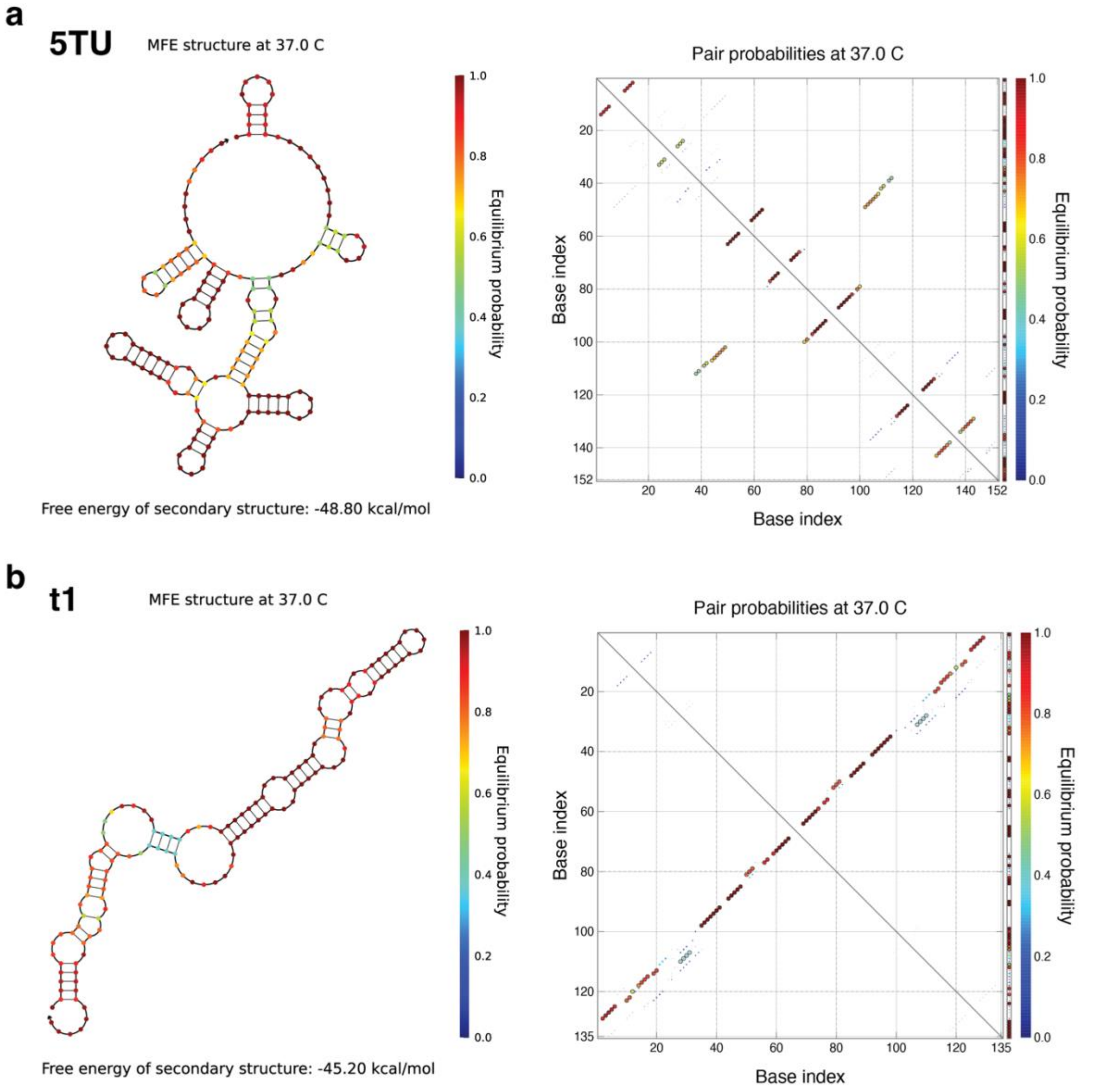

**Fig. S9. Secondary structure prediction for 5TU and t1.**

(**a**,**b**) Secondary structure prediction for 5TU and t1 using NUPACK. Left shows secondary structure coloured with equilibrium probabilities. Right shows dotplot with pair probabilities.

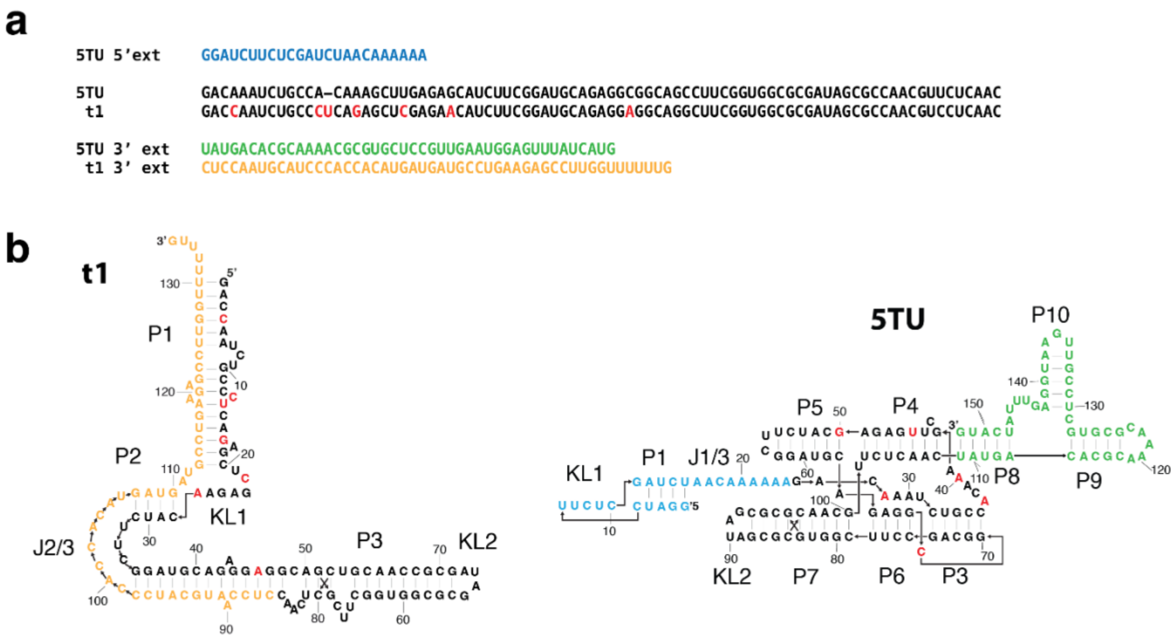

**Fig. S10. Primary and secondary structure comparison between 5TU and t1 subunits.** (**a**) Sequences for 5TU and t1. 5TU has a unique 5' extension (blue). The core domain has 7/83 mutations (red) between 5TU and t1. 5TU and t1 has unique 3' extensions coloured green and orange, respectively. (**b**) Secondary structure of 5TU and t1 subunits colored according to the domains and mutations annotated in panel a.

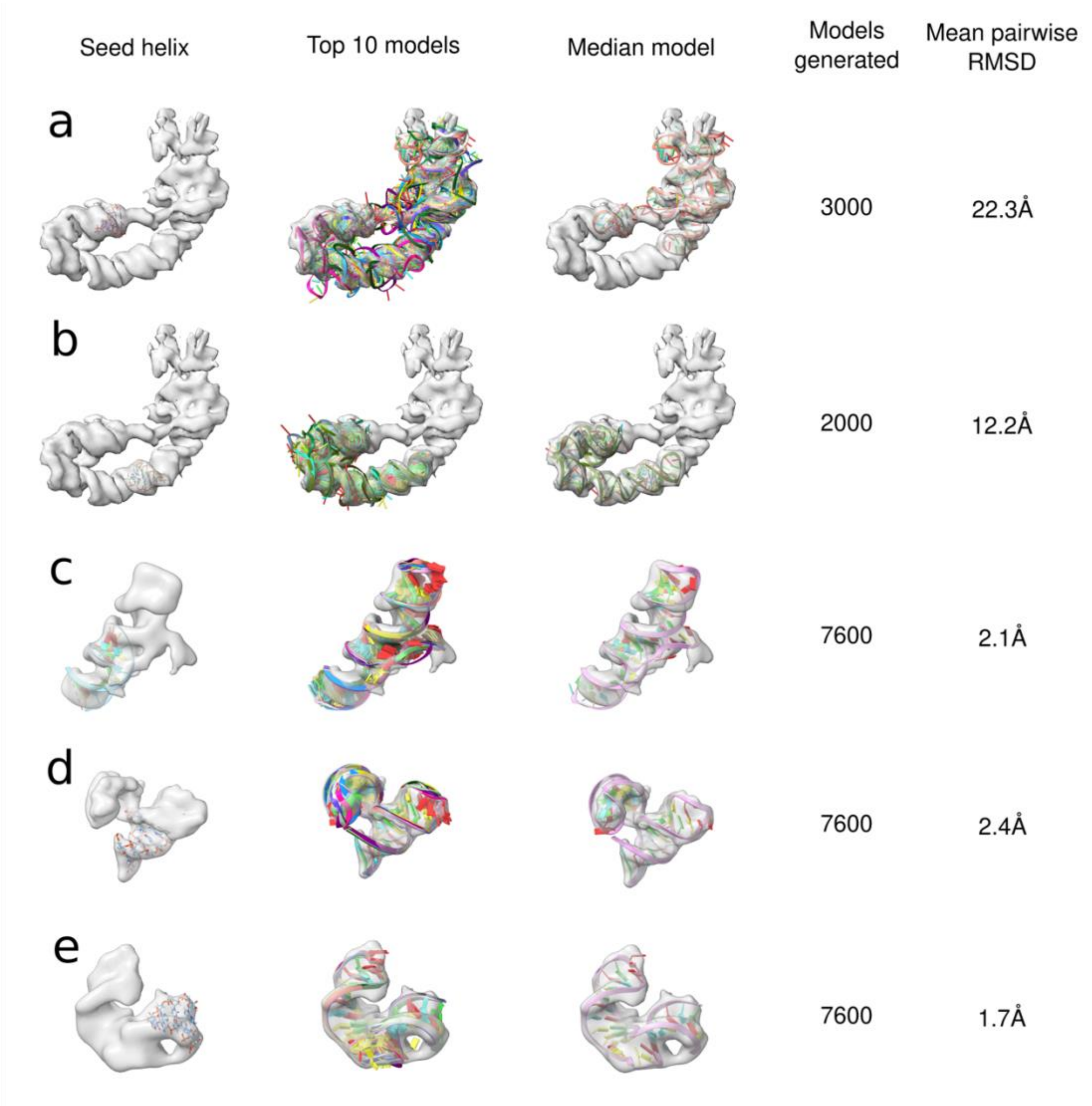

**Fig. S11. DRRAFTER modelling of 5TU and t1 using the 5TU+t1-anneal map.** From left to right the initial helix placement is shown, then the top 10 models from the DRRAFTER runs, then the top model. DRRAFTER runs were performed for (**a**) the entire 5TU, (**b**) the entire t1, (**c**) 5TU:J3/4 with 5TU:P4-P5-P8-P9, (**d**) 5TU:J10/9 with P8-P9-P10, (**e**) t1:J3/2 with P2 & P3.

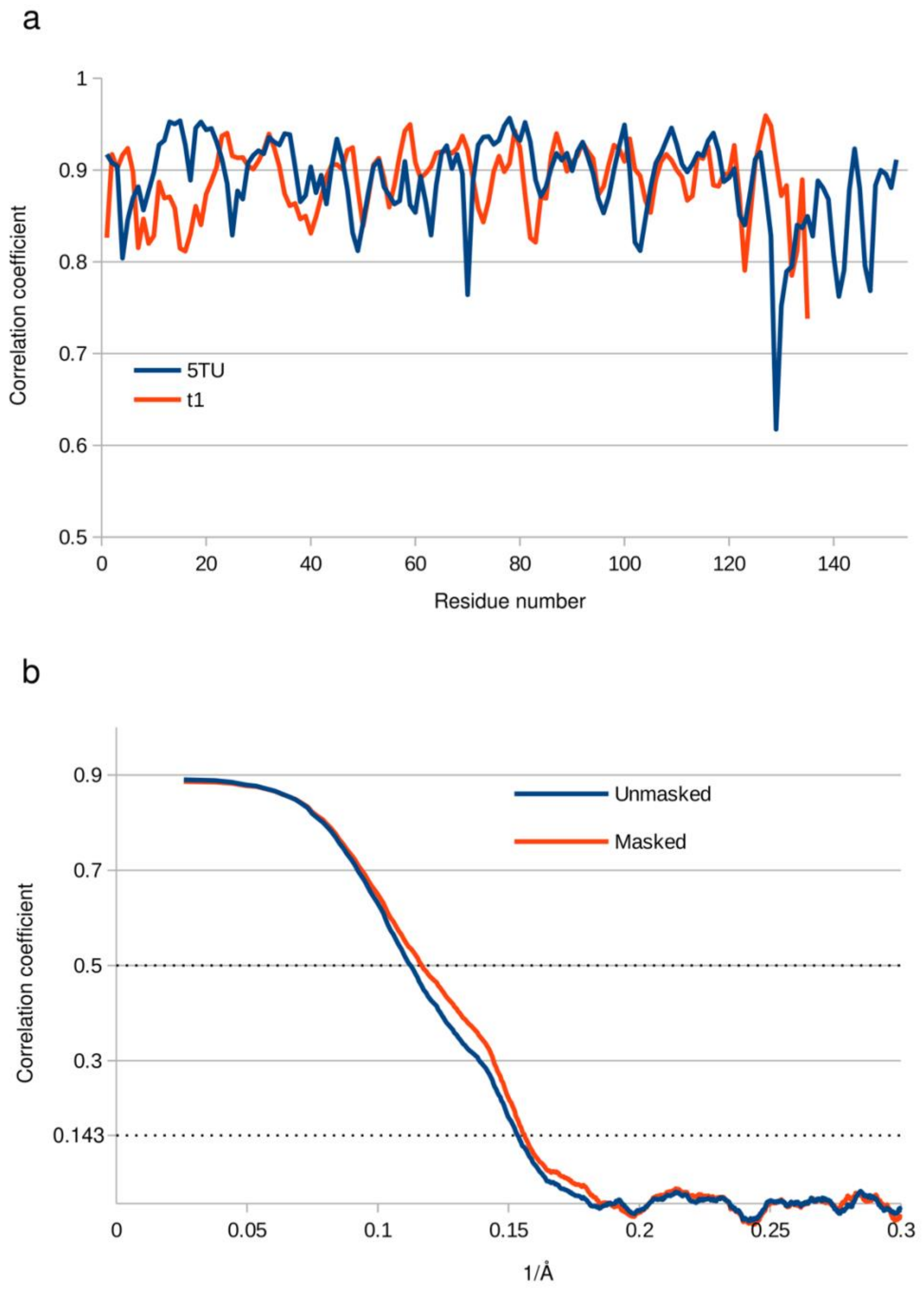

**Fig. S12. Map to model comparison for the 5TU+t1-anneal sample.**

(**a**) Correlation coefficient for 5TU and t1 for each residue. (**b**) Correlation coefficient for the 5TU+t1 heterodimer at different resolutions. A soft mask was generated from the atomic model(17).

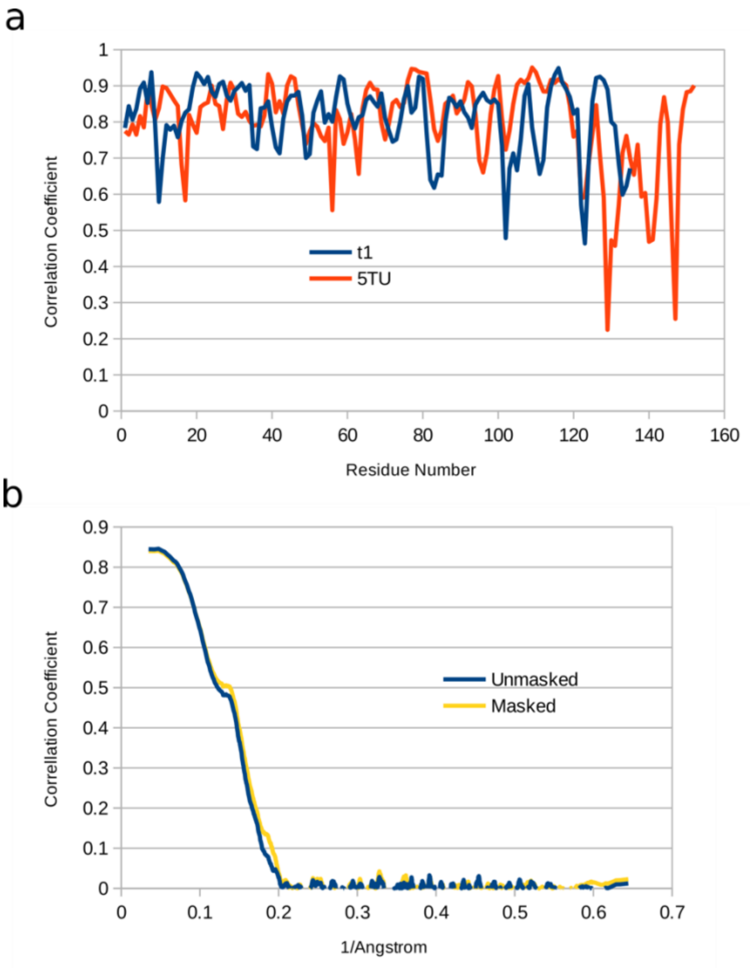

**Fig. S13. Map to model comparison using Fourier shell correlation for the 5TU+t1-cofold sample.**

(**a**) Correlation coefficient for 5TU and t1 for each residue. (**b**) Correlation coefficient for the 5TU+t1 heterodimer at different resolutions. A soft mask was generated from the atomic model (17).

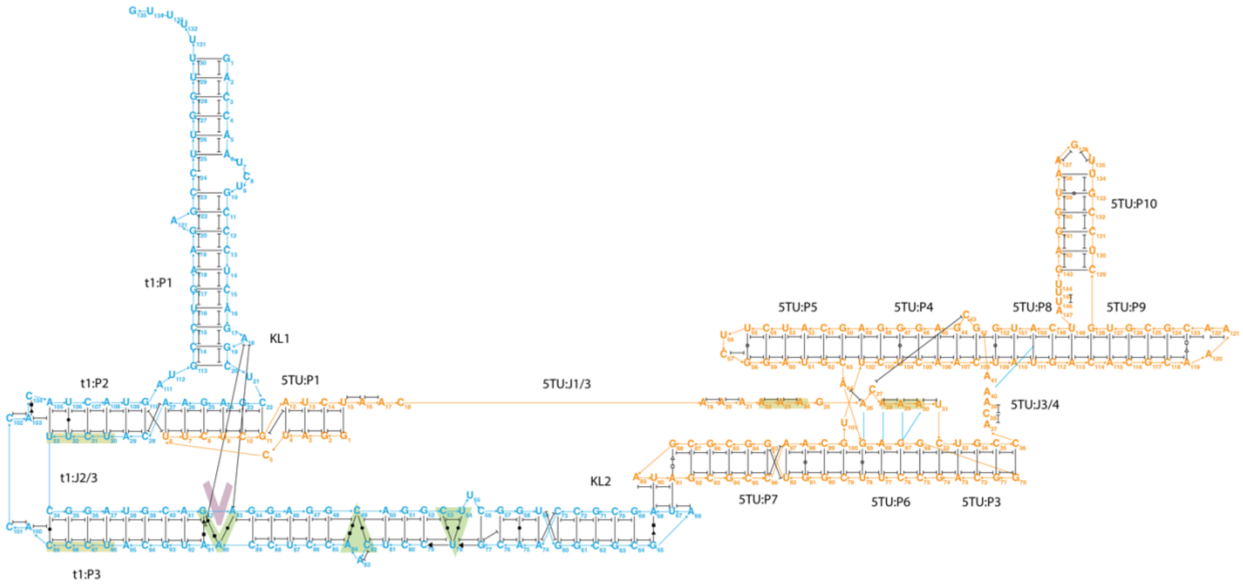

**Fig. S14. Map of the secondary and tertiary interactions of TPR model.**

Secondary structures of 5TU (orange) and t1 (cyan) are shown with annotation of base pairs (black lines), base stacks (black capped lines) and A-minor interactions (cyan lines). Selected tertiary motifs are annotated by green and purple symbols: V shape for A18 intercalation in G42-A43-A90 motif. Triangles for 120-degree bending motif involving C53-U54-U78 and C49, A84-C82. Primary sequence motifs are marked in yellow: C96-C99, U30-U33, A22-24, A26-A30.

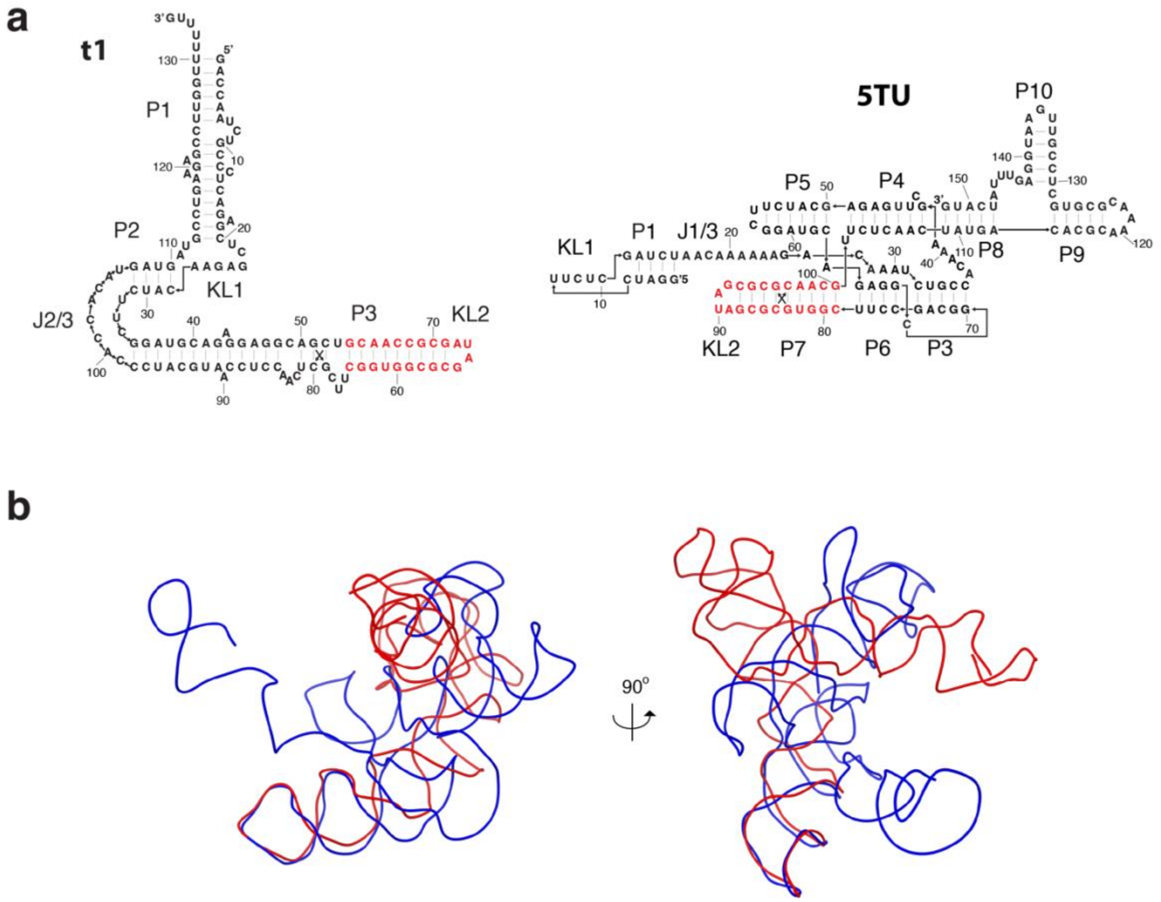

**Fig. S15. Structural comparison between 5TU and t1.**

(**a**) Secondary structure of 5TU and t1 with the only similar hairpin of 5TU:P7 and t1:P3 shown in red. (**b**) Structural alignment of the 5TU:P7 and t1:P3 hairpins to highlight the structural difference between the 5TU (blue) and t1 (red) subunits shown as ribbon diagrams at 90 degree rotated views.

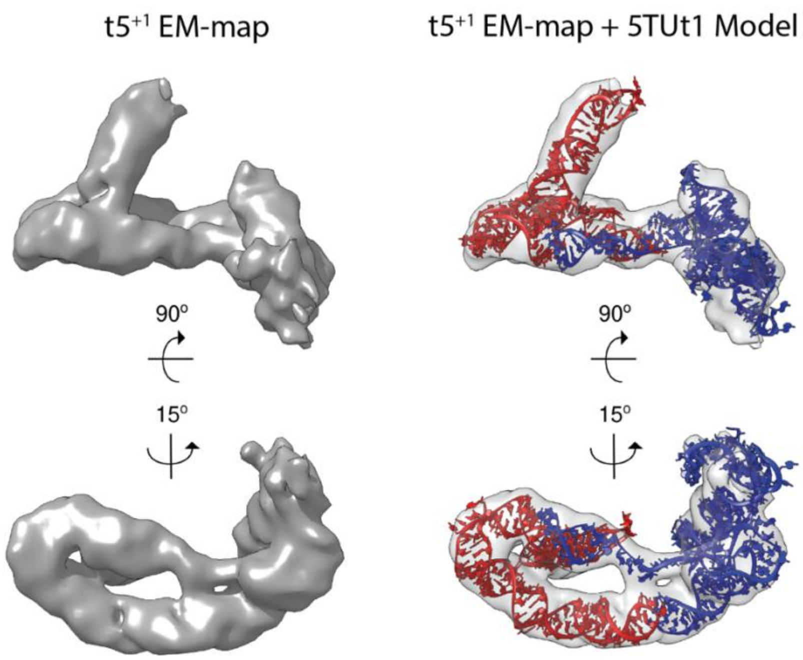

**Fig. S16. Comparison between t5$^{+1}$ EM-map and 5TU+t1 model.**

Left panels are the t5+1 EM-map. Right panels are the 5TU+t1 model docked into t5+1 EM-map (7.99 Å resolution). The docking was performed in Chimera.

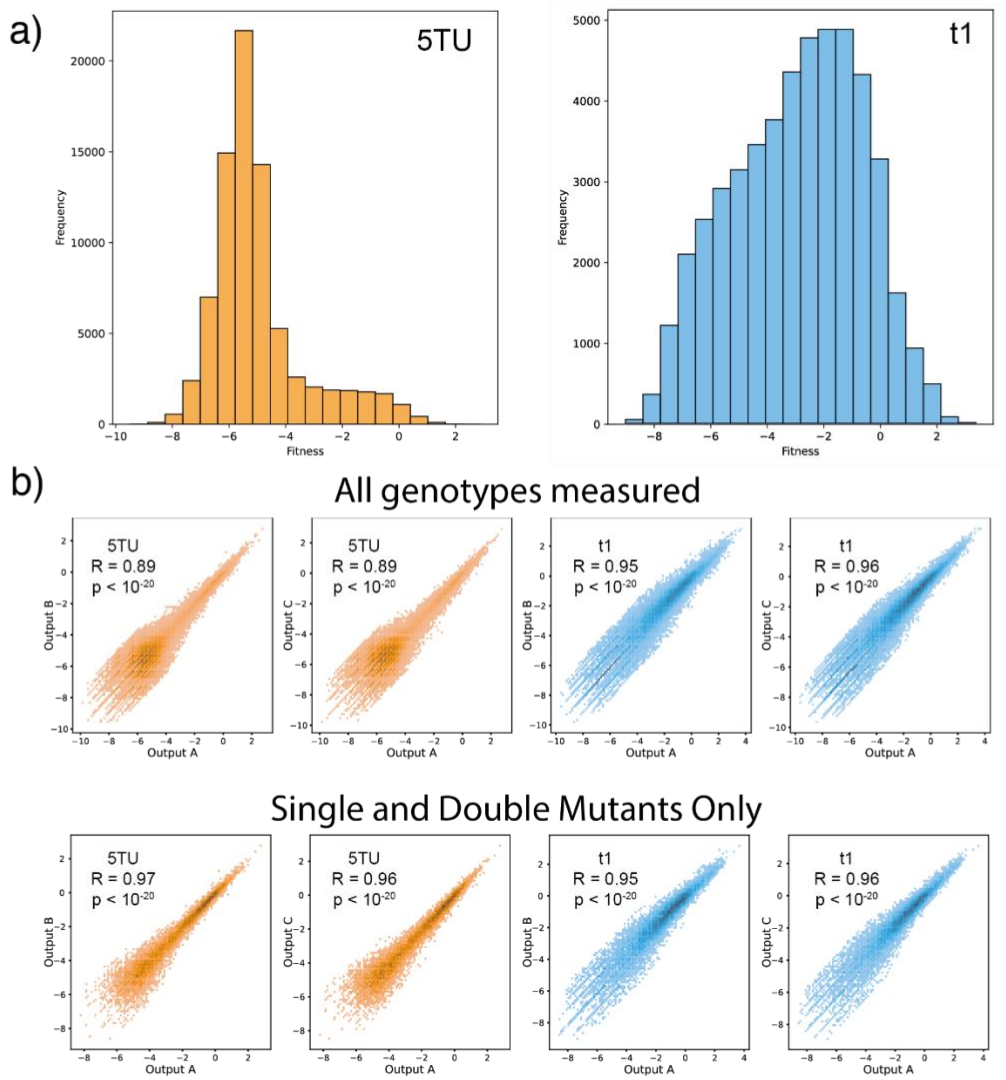

**Fig. S17. Adaptive landscape of TPR.**

(**a**) Distribution of fitness values of 5TU and t1 genotypes. Distribution of 5TU fitness values is much sharper than that of t1 genotypes. (**b**) Top row: correlation between calculated log-transformed fitness values of all ribozyme genotypes in different replicates. R = Pearson correlation coefficient, n = 79,702 for 5TU, n = 49,006 for t1. Bottom row: correlation between calculated log-transformed fitness values of single and double mutant genotypes in different replicates. R = Pearson correlation coefficient, n = 10,806 for 5TU, n = 17,086 for t1.

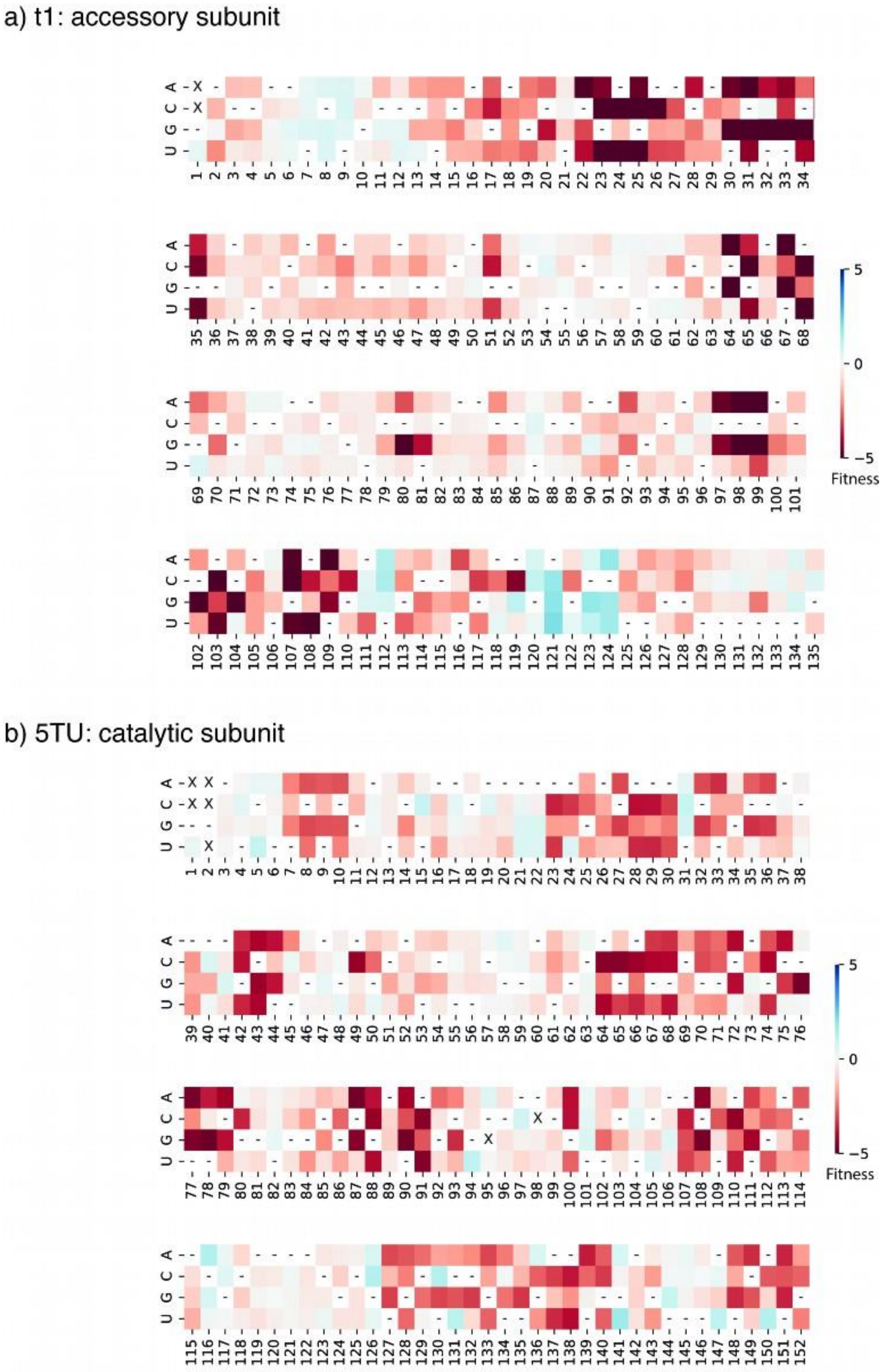

**Fig. S18. Fitness landscape of TPR.**

(**a**) Fitness values of t1 point mutants. "-" indicate wild-type base; "X" indicate that fitness of genotype could not be calculated. (**b**) Fitness values of 5TU point mutants. "-" indicate wild-type base; "X" indicate that fitness of genotype could not be calculated. In 5TU, G1 and G2 were kept unmutagenised due to the recovery primer (forceGG) used for RT-PCR; in t1, G1 was kept unmutagenised for the same reason (t1rec primer used). Hence, the fitness of mutants at this position were not measured.

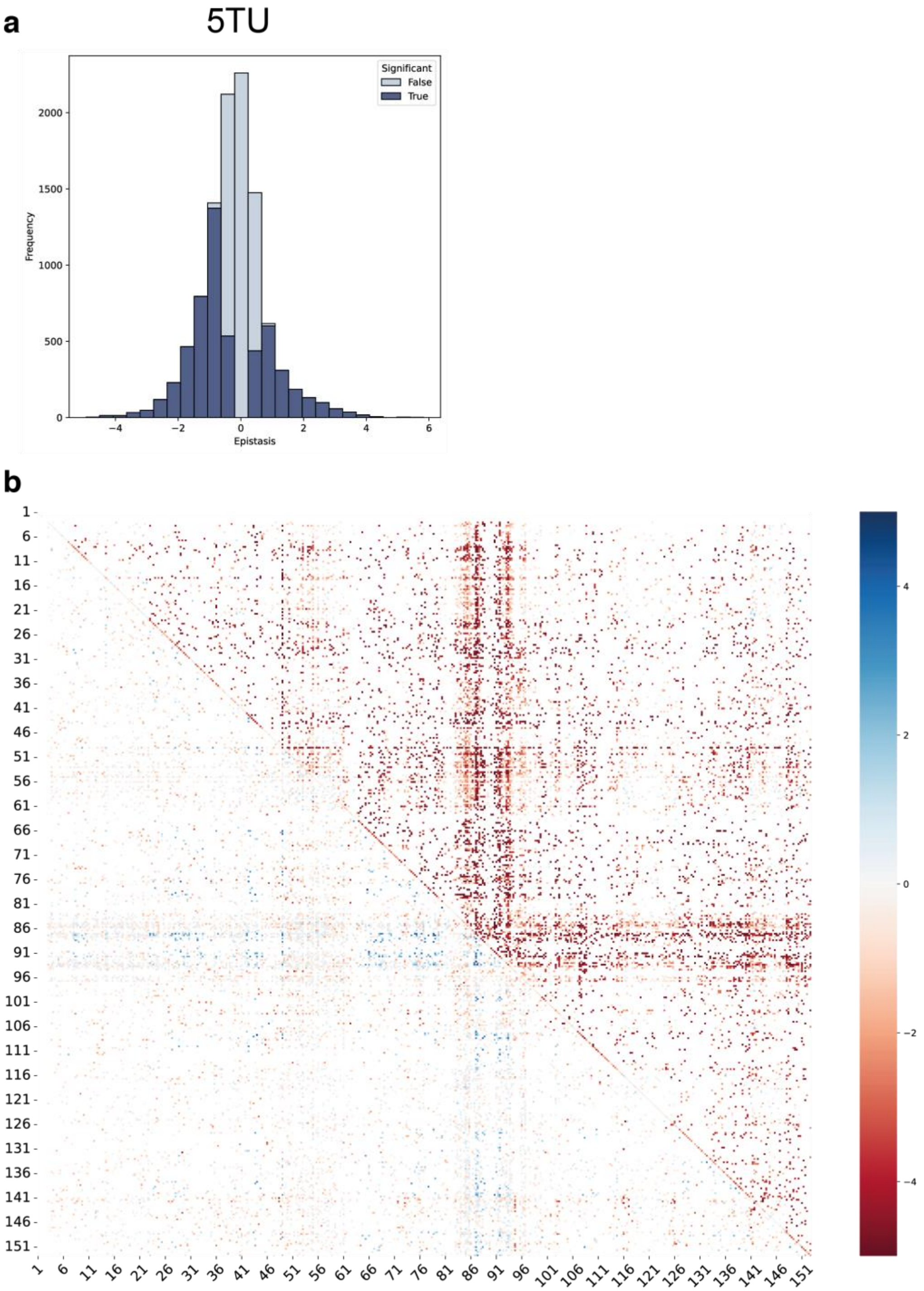

**Fig. S19. 5TU double fitness matrix.**

(**a**) Distribution of epistasis in 5TU double mutants. Significant epistasis values coloured in dark blue (False Discovery Rate: 10.1%), non-significant epistasis in light blue. In both subunits, epistasis is negatively biased. (**b**) Upper right triangle shows estimated fitness of all 5TU double mutants present in dataset. Lower left triangle shows estimated epistasis of double mutants. Scale bar refers to both fitness and epistasis, depending on the sector of the figure in question.

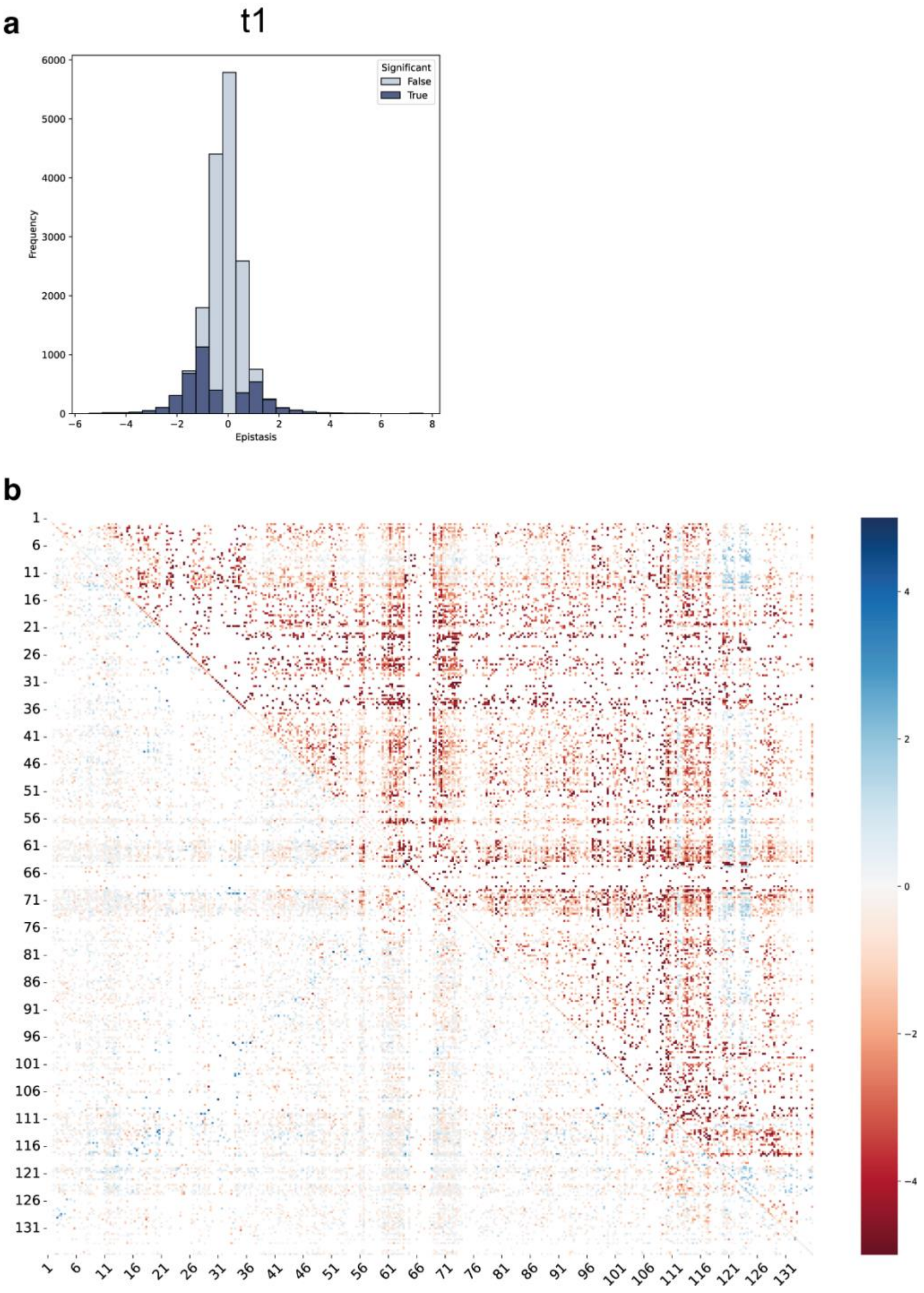

**Fig. S20. t1 double fitness matrix.**

(**a**) Distribution of epistasis in t1 double mutants. Significant epistasis values coloured in dark blue (False Discovery Rate: 16%), non-significant epistasis in light blue. In both subunits, epistasis is negatively biased. (**b**) Upper right triangle shows estimated fitness of all g1 double mutants present in dataset. Lower left triangle shows estimated epistasis of double mutants. Scale bar refers to both fitness and epistasis, depending on the sector of the figure in question.

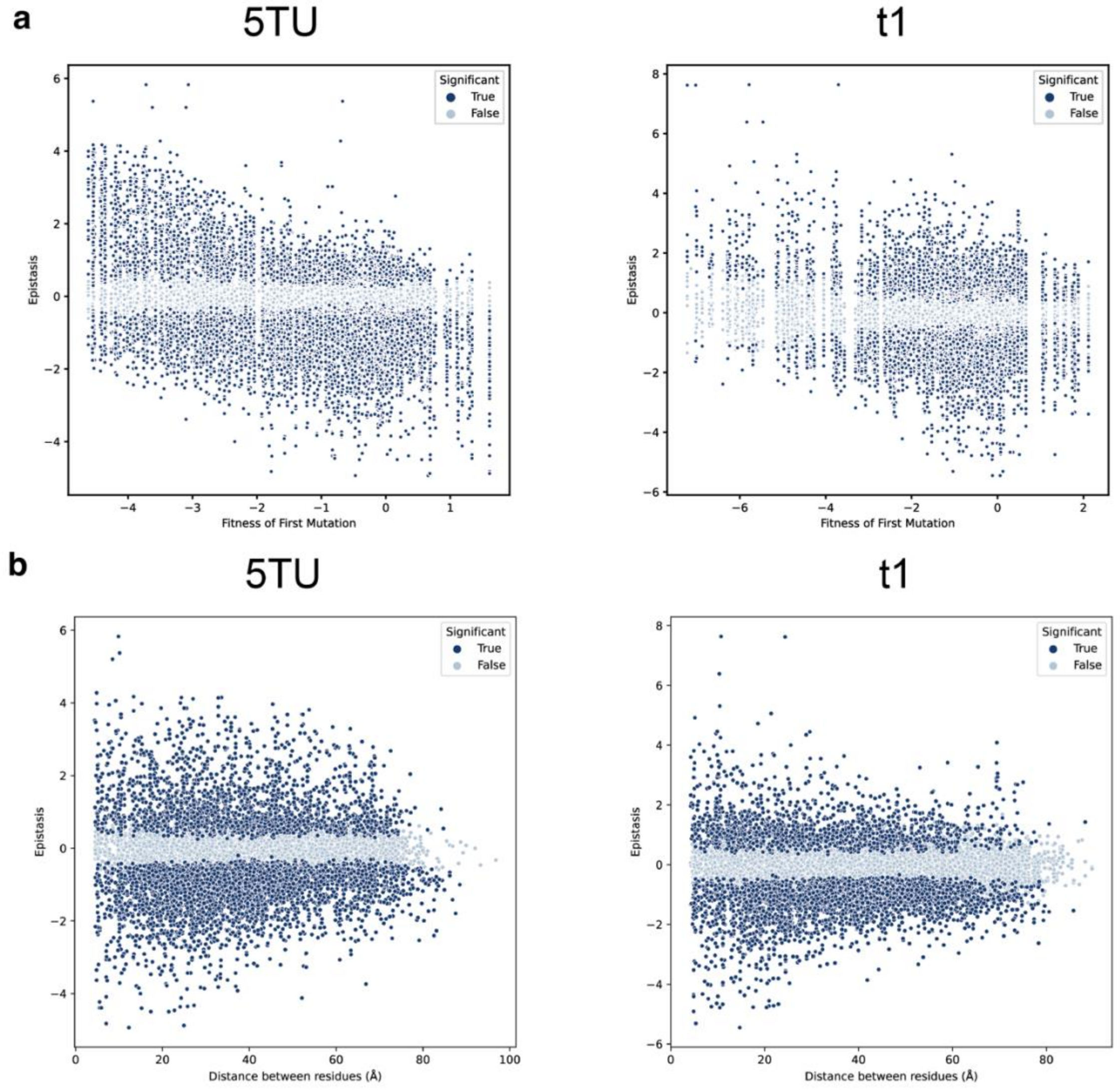

**Fig. S21. Epistasis of TPR double mutants.**

(**a**) Mean epistasis value between first and second mutations decreases with the fitness of the first mutation. Significant epistasis values coloured in dark blue (False Discovery Rate: 10.1% for 5TU, 16% for t1), non-significant epistasis in light blue. (**b**) For both 5TU and t1, the magnitude and significance of epistasis decreases as residues become more distant. Distances between residues are measured between the 1’ carbons in the ribose ring. Significant epistasis values coloured in dark blue (False Discovery Rate: 10.1% for 5TU, 16% for t1), non-significant epistasis in light blue.

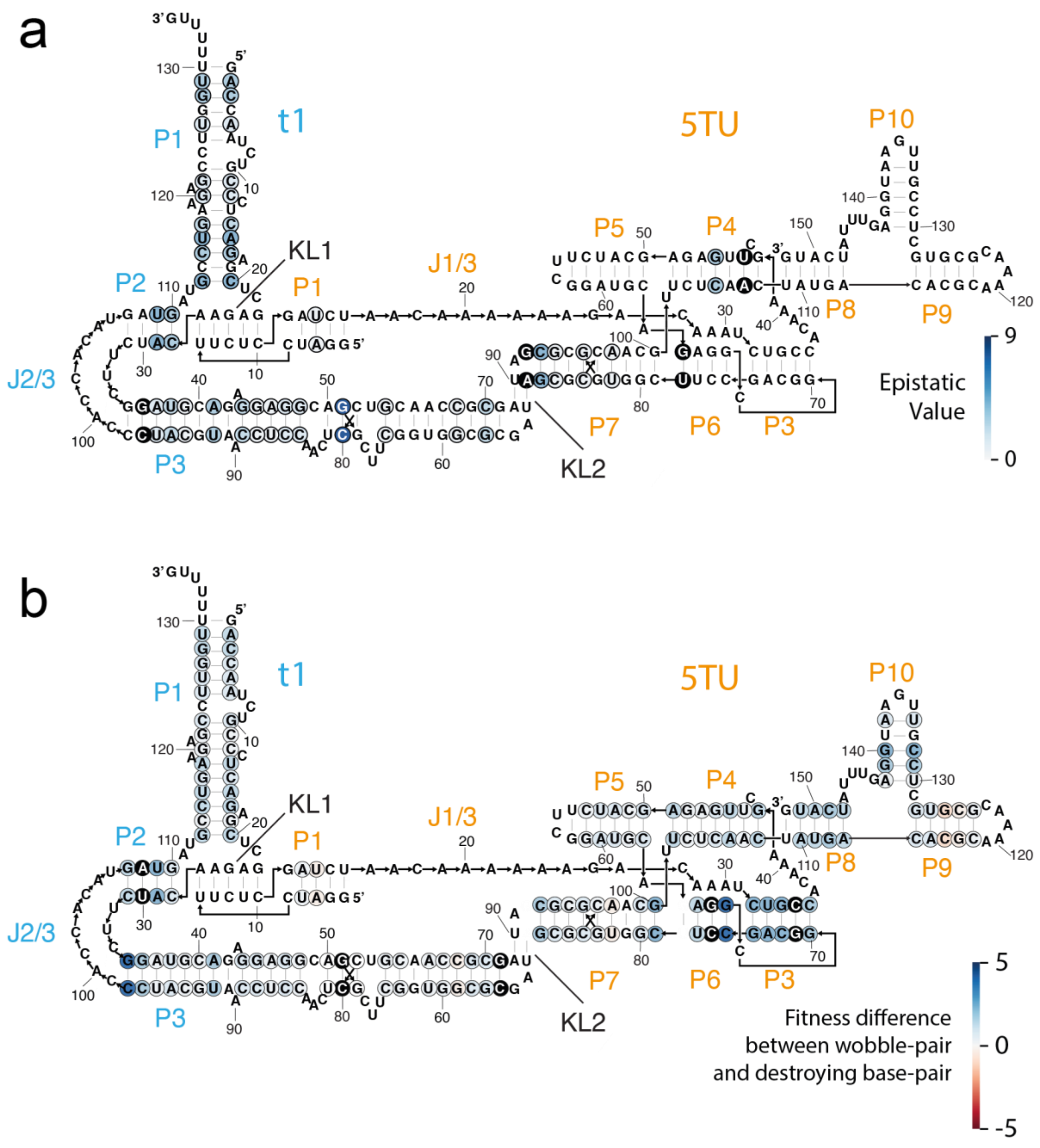

**Fig. S22. Epistasis of TPR double mutants at base-pairing positions.**

(**a**) Secondary structure of the 5TU+t1 TPR. Base-pairs are circled and coloured by epistatic value if basepair-preserving double mutants which exhibit positive epistasis can be found within the dataset. (**b**) Secondary structure of the 5TU+t1 TPR. Base-pairs are circled and coloured by the difference between fitness of point mutant that generates wobble pair, and the average fitness of point mutants which disrupt base pairing at the given position.

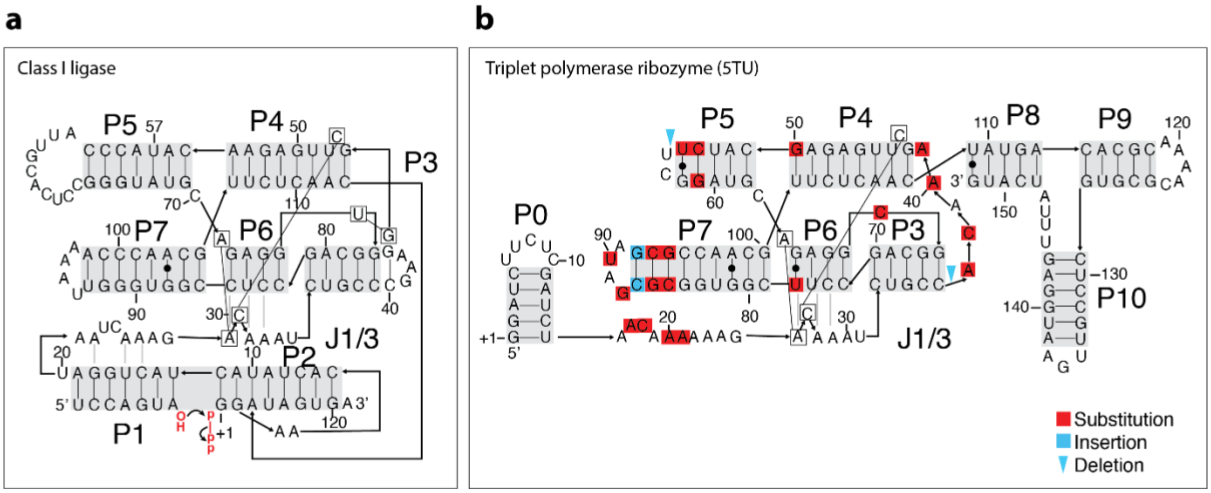

**Fig. S23. Structural comparison of class I ligase and 5TU.**
(**a**) Secondary structure model of cIL showing stem regions (P1-P7) and central base stacks (connected boxes) and A-minor interactions (grey lines). (**b**) Secondary structure model of TPR showing stem regions (P1-P10) with similar positioning of helices and annotation as in panel a. Mutational differences are indicated: substitution (red), inserts (blue), deletions (blue arrows).

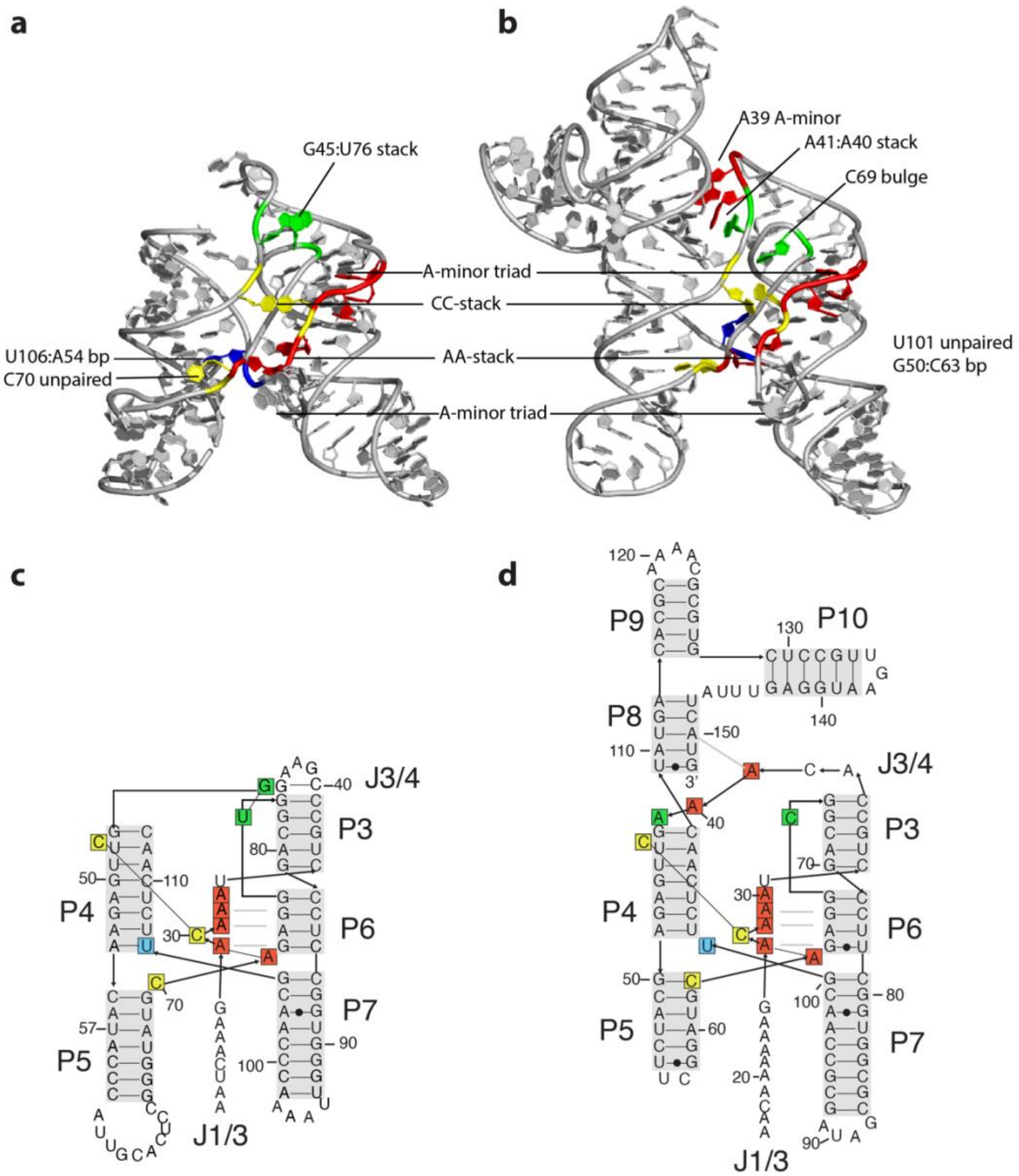

**Fig. S24. Structural comparison of class I ligase and 5TU.**

(**a**) Tertiary structure model of cIL showing stem regions (J1/3-P7) and central base stacks (connected boxes) and A-minor interactions (grey lines). (**b**) Tertiary structure model of TPR showing stem regions (J1/3-P10) with similar positioning of helices and annotation as in panel a. (**c,d**) Corresponding secondary structure models of cIL and TPR as shown in panel a,b. Bases are colored as follows: C (yellow), U (cyan), A (red). G45:U76 stack in cIL colored green to compare to similar positions A41 and C69 in TPR to highlight broken base stack.

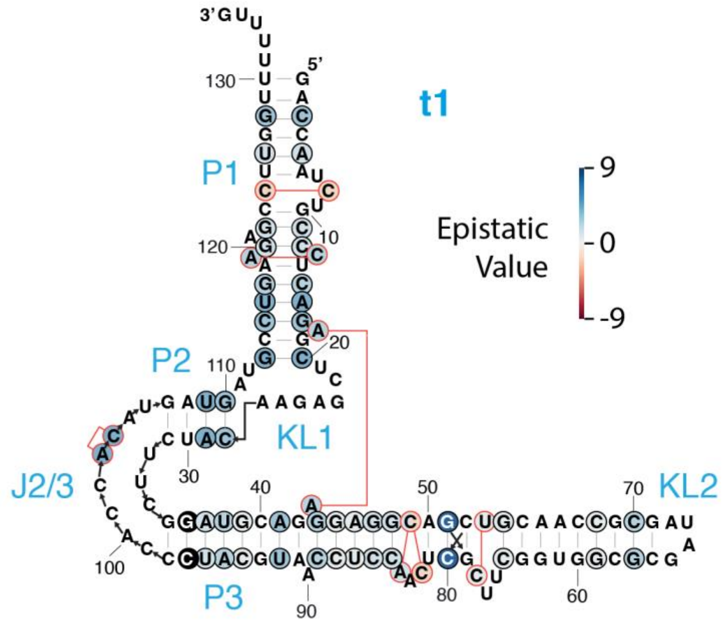

**Fig. S25. Secondary structure of t1 with epistasis values annotated for standard and non-canonical base pairs.**

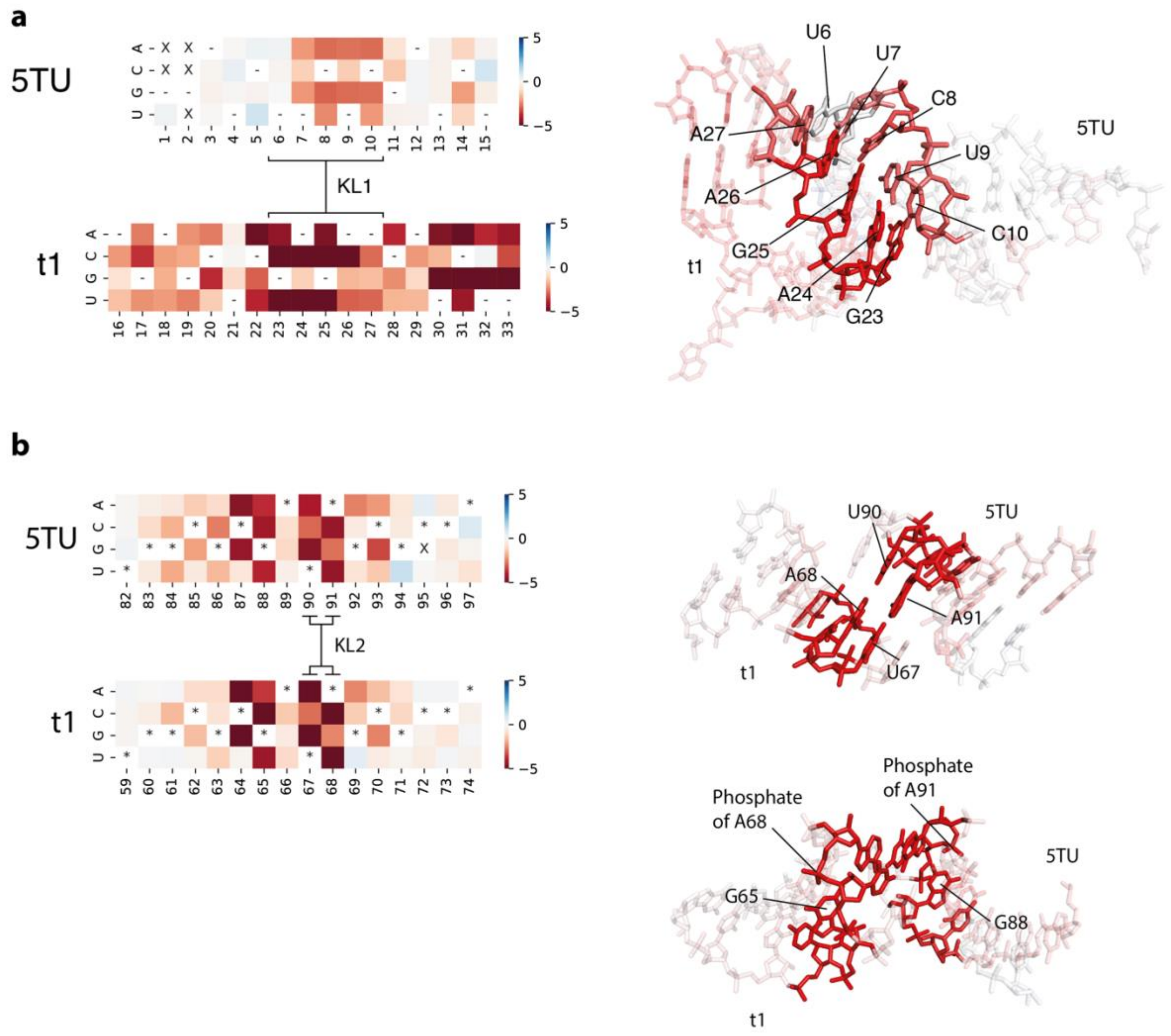

**Fig. S26. Adaptive landscape of kissing-loops KL1 & 2.**

(**a**) Left: Fitness of point mutants at positions within KL1 (G1-U15 in 5TU; A16-U33 in t1). “-“ indicate wild-type base; “X” indicate fitness of genotype could not be calculated. Right: 3D structure of KL1. Nucleotides are coloured to reflect the average fitness of point mutants at each position. For clarity, positions U6 – C10 in 5TU, and G23 – A27 in t1 are opaque; all other positions are displayed with 50% transparency. (**b**) Left: Fitness of point mutants at positions within KL2 (U82-A97 in 5TU; U59-A74 in t1). “-“ indicate wild-type base; “X” indicate fitness of genotype could not be calculated. Right: 3D structure of KL2. Nucleotides are coloured to reflect the average fitness of point mutants at each position. For clarity, positions C87, G88, U90, A91 in 5TU, and C64, G65, U67, A68 in t1 are opaque; all other positions are displayed with 50% transparency.

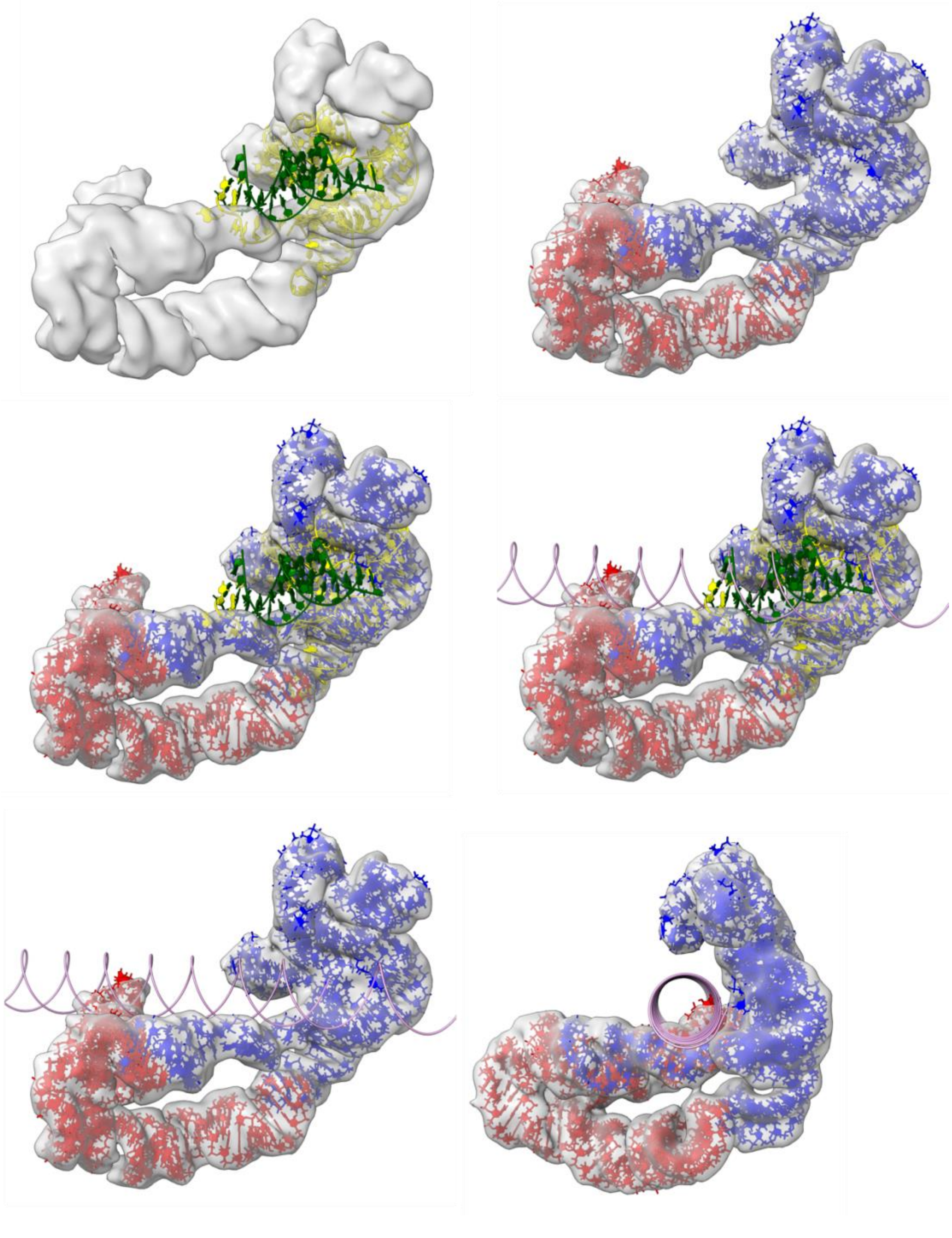

**Fig. S27. Alignment of the TPR and Class 1 ligase structures.**

Top left panel shows our TPR volume with the class 1 ligase structure colored yellow with green template. Top right panel shows the fit of our model to the volume. The middle panel shows an overlay of the class 1 ligase and our model with an extended template shown on the right. The bottom panel shows only our model with the extended template from two views.

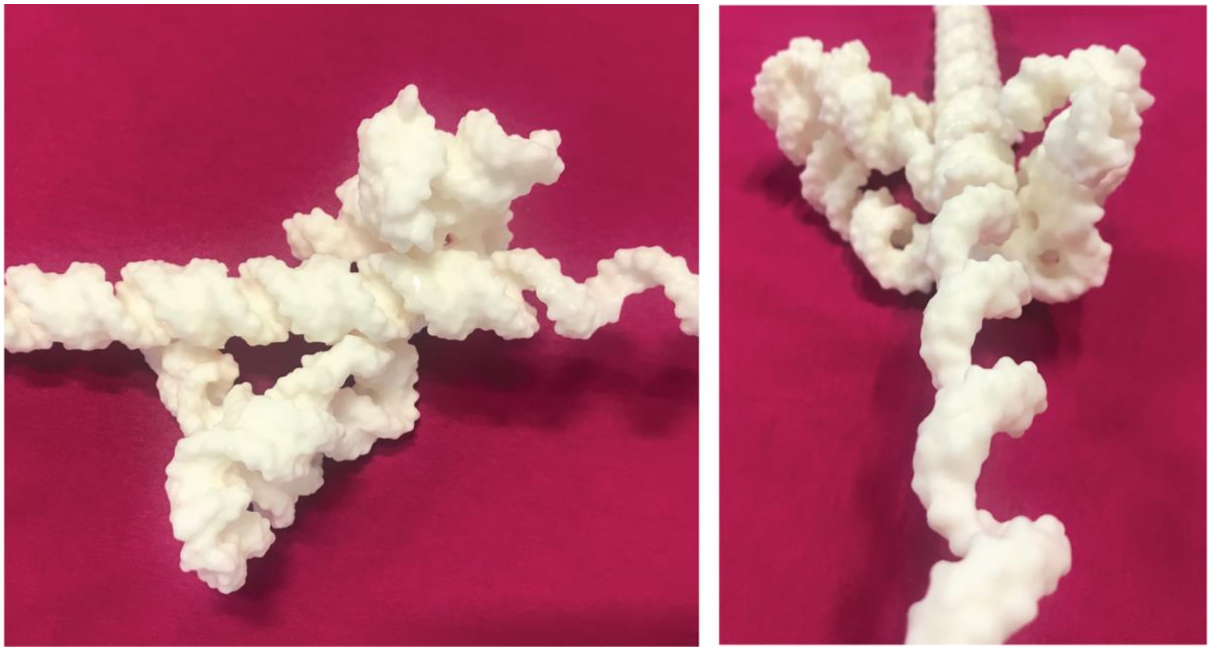

**Fig. S28. 3D model of TPR and primer-template duplex.**

The TPR and a primer-template duplex were 3D printed separately and were fitted together in accordance with the 3D modelling (Fig. S22). The model shows that the minor groove of the primer-template duplex can contact the J1/3, P10 and t1:P1 while being in close proximity to the active site of P4. The model is shown in side view and from the perspective of the yet uncopied template.

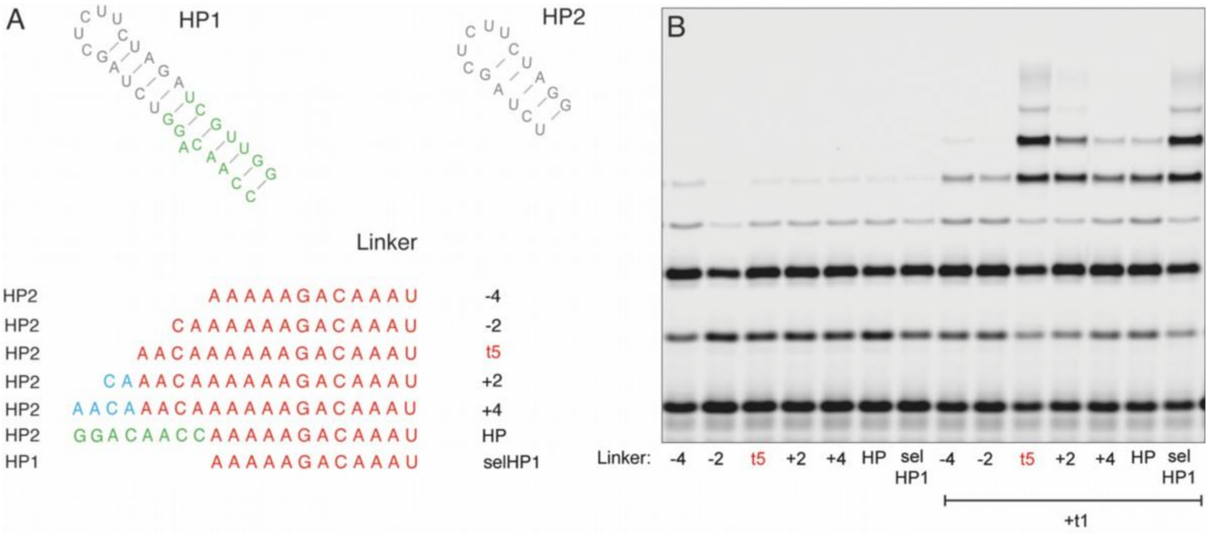

**Fig. S29. J1/3 linker length and TPR activity.**

(**a**) TPR activity as a function of J1/3 linker length, preceded either with 5TU 5’ hairpin (HP2) or 5’ hairpin used in original t5 selection (HP1) showing different constructs (HP2-4, HP2-2, t5wt, HP2+2, HP2+4, HP2HP, selHP1) and (**b**), primer extension activity of different constructs either in the absence (left) or presence (+ t1, right) of the t1 accessory subunit. Only the correct J1/3 spacing (t5wt) or a shorter J1/3 in combination with a larger (HP1) 5’- hairpin shows full triplet polymerase activity.

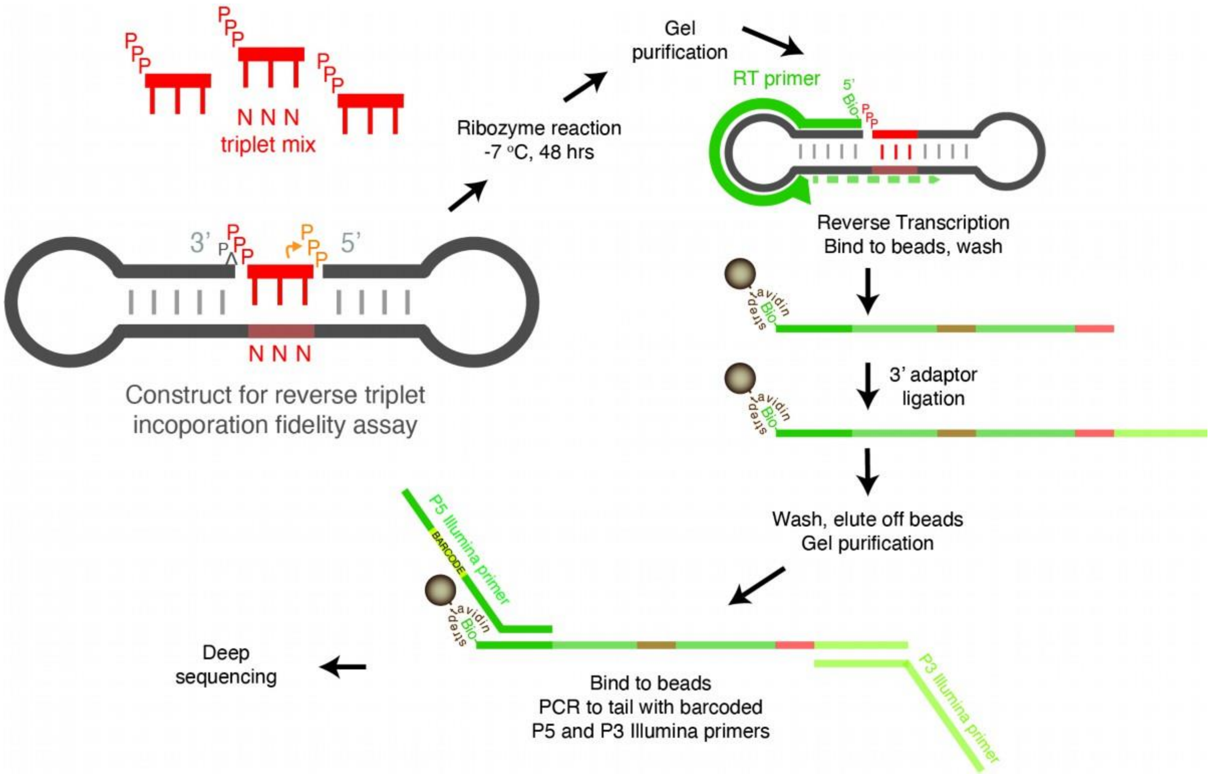

**Fig. S30. FidelitySeq assay.**

Schematic of FidelitySeq assay to assess triplet incorporation fidelity in the 3’-5’ direction. ^ character indicates a 2’,3’-cyclic phosphate generated by HDV ribozyme cleavage, which is not ligatable by the ribozyme. This workflow enabled sequence reads to include both the template (brown) and incorporated triplet (red). DNA species depicted in green, RNA species in grey.

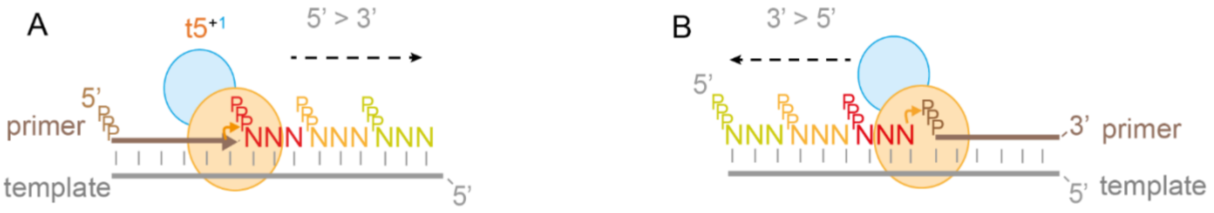

**A** — % incorporated:

| Base encoded | A | C | G | U |
|---|---|---|---|---|
| A | 85.5% | 0.6% | 13.7% | 0.2% |
| C | 0.1% | 98.7% | 0.3% | 1.0% |
| G | 0.1% | 0.2% | 99.6% | 0.1% |
| U | 0.1% | 0.3% | 0.2% | 99.4% |
| 1st position fidelity: | | | | 95.6% |

| Base encoded | A | C | G | U |
|---|---|---|---|---|
| A | 96.6% | 0.2% | 2.2% | 1.0% |
| C | 0.3% | 98.1% | 0.3% | 1.3% |
| G | 0.2% | 0.5% | 98.8% | 0.4% |
| U | 0.1% | 0.3% | 0.2% | 99.4% |
| 2nd position fidelity: | | | | 98.2% |

| Base encoded | A | C | G | U |
|---|---|---|---|---|
| A | 97.4% | 1.0% | 1.4% | 0.2% |
| C | 0.6% | 98.0% | 0.3% | 1.1% |
| G | 0.3% | 0.5% | 98.8% | 0.3% |
| U | 0.2% | 0.4% | 0.3% | 99.1% |
| 3rd position fidelity: | | | | 98.3% |

Overall fidelity: 97.4%

**B** — % incorporated:

| Base encoded | A | C | G | U |
|---|---|---|---|---|
| A | 44.8% | 6.1% | 42.5% | 6.7% |
| C | 0.8% | 96.4% | 1.6% | 1.2% |
| G | 0.7% | 3.3% | 94.5% | 1.5% |
| U | 1.5% | 5.7% | 14.6% | 78.1% |
| 1st position fidelity: | | | | 75.1% |

| Base encoded | A | C | G | U |
|---|---|---|---|---|
| A | 57.6% | 3.8% | 36.5% | 2.1% |
| C | 1.1% | 95.2% | 1.9% | 1.8% |
| G | 0.8% | 2.2% | 96.0% | 0.9% |
| U | 1.0% | 3.7% | 2.2% | 93.1% |
| 2nd position fidelity: | | | | 83.7% |

| Base encoded | A | C | G | U |
|---|---|---|---|---|
| A | 93.8% | 2.0% | 3.2% | 0.9% |
| C | 2.6% | 92.4% | 1.5% | 3.4% |
| G | 2.0% | 1.9% | 95.2% | 0.9% |
| U | 2.2% | 2.6% | 2.2% | 93.0% |
| 3rd position fidelity: | | | | 93.6% |

Overall fidelity: 83.8%

**Fig. S31. Fidelity of different polymerisation modes.**

Schematic of 5' to 3' forward (**a**) and 3' to 5' reverse (**b**) polymerisation. +1 and -1 triplets shown in red, +2 in yellow and +3 in pale green and associated fidelity profiles of forward triplet (a) and reverse (b) incorporations, as determined by FidelitySeq (extra SI figure 30), revealing high overall fidelity, with a tolerance for G:U wobble pairs at the first position for forward synthesis and a lower overall fidelity and broader error profile for reverse synthesis (b). Position and overall fidelities are calculated as geometric means.

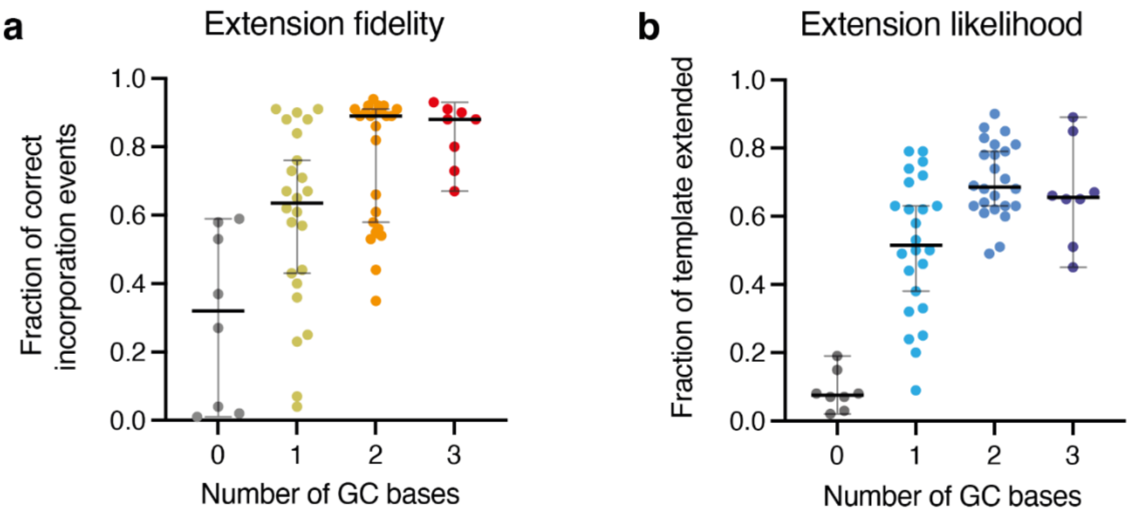

**Fig. S32. 3’-5’ triplet extension and triplet GC content.**

Non-canonical 3’-5’ triplet incorporation fidelity and extension correlate with triplet GC content. (**a**) Comparison of the fidelity of triplet incorporation and the number of GC bases in the triplet. Fidelity scores for each of the 64 triplet combinations were determined using the number of sequencing-reads from correct incorporation events as a fraction of all incorporation events for that template sequence. (**b**) Comparison of the likelihood of triplet incorporation and the number of GC bases in the triplet. The likelihood of extension was determined using the number of reads for each template which have had a triplet incorporated as a fraction of the total number of reads for that template. Error bars represent median values and 95% confidence intervals.

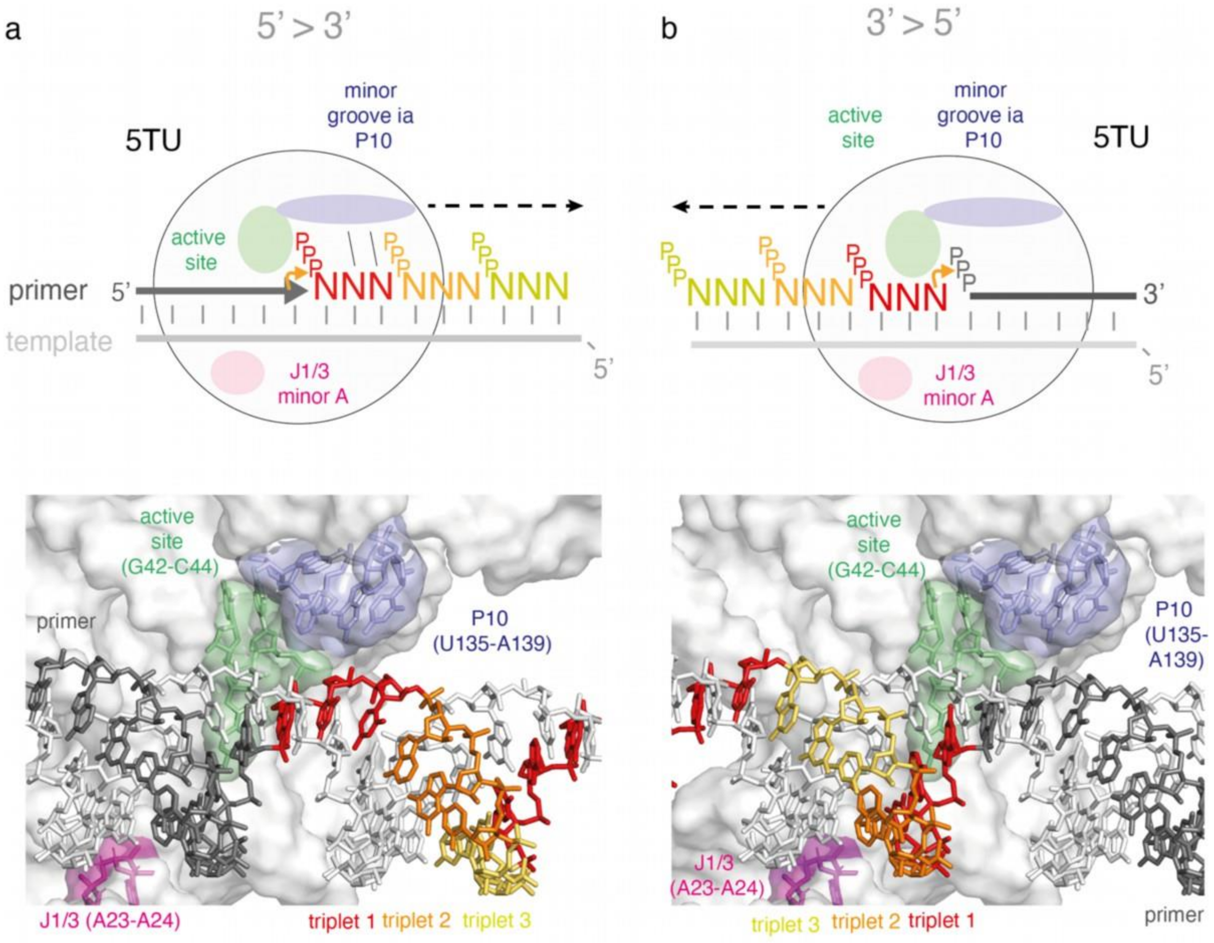

**Fig. S33. Structural context of different polymerisation modes.**

(**a**, **b**) Cartoon (top) and local TPR holoenzyme structure model showing the two RNA synthesis modes of the TPR, in the canonical 5’-3’ direction (a) and the non-canonical reverse mode (3’-5 direction), with primer (dark grey), template (light grey), 1st triplet to be incorporated in respective modes (triplet 1 (red)), triplet 2 (orange), triplet 3 (yellow) (next triplet 4 (again red)). Also shown P10 (light blue), active site (light green) and J1/3 (magenta).

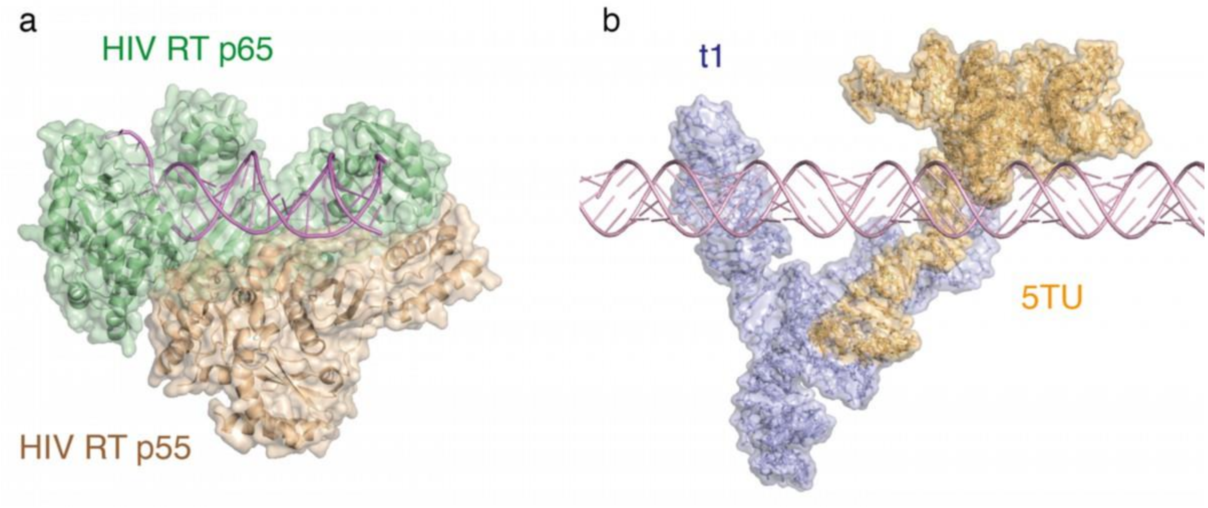

**Fig. S34. Heterodimeric polymerases HIV RT and 5TU+t1 TPR.**

Side by side comparison of heterodimeric HIV RT structure (5TXM.pdb)(29) (left) with 2 subunits (catalytic subunit p65 (green) and accessory subunit p55 (wheat)) and DNA-primer template duplex (pink) and heterodimeric all-RNA TPR structure holoenzyme model (right) with 5TU catalytic subunit (orange) and accessory subunit t1 (light blue) and model RNA primer - template duplex (light pink).

**Table S1. Cryo-EM data collection, refinement, and validation statistics.**

| | 5TU+t1-anneal | 5TU+t1-cofold | $t5^{+1}$ |
|---|---|---|---|
| **Data collection and processing** | | | |
| Magnification | 130000 | 130000 | 105000 |
| Voltage (kV) | 300 | 300 | 300 |
| Electron exposure (e–/Å$^2$) | ~60 | ~60 | 65 |
| Defocus range (μm) | -0.5 to -2.2 | -0.5 to -2.2 | -1.2 to -2.6 |
| Pixel size (Å) | 0.647 | 0.647 | 1.1 |
| Symmetry imposed | None | None | None |
| Initial particle images (no.) | | | |
| Final particle images (no.) | 26167 | 86545 | 5485 |
| Map resolution (Å)<br>FSC threshold (0.143) | 5.94 | 5.00 | 7.99 |
| **Refinement** | | | |
| Initial model used (PDB code) | | PDB:3IVK,1F5U | |
| Model resolution (Å)<br>FSC threshold (0.143) | | 5.6 | |
| Map sharpening *B* factor (Å$^2$) | | 288 | |
| Model composition | | 9206 atoms<br>3097 hydrogens<br>287 nucleotides | |
| R.m.s. deviations<br>Bond lengths (Å)<br>Bond angles (°) | | <br>0.006 (0)<br>0.797 (0) | |
| Validation<br>MolProbity score<br>Clashscore | | <br>1.65<br>0 | |

**Table S2. Epistasis of 5TU bp positions.**

**Double mutants at canonical basepairing positions in 5TU with statistically significant epistasis**

| Genotype | Restores basepair? | Fitness of genotype | Epistasis |
|---|---|---|---|
| T33G A73G | False | -0.513 | 1.756 |
| C35T G71T | False | -3.531 | -1.510 |
| T44A A107T | True | -0.956 | 5.829 |
| G46C C105G | True | 1.294 | 3.023 |
| G65T T78G | True | -5.645 | 2.073 |
| G65T T78A | True | -1.522 | 5.203 |
| G86T C93A | True | -0.194 | 3.600 |
| G86T C93G | True | -1.278 | 3.687 |
| C87A G92T | True | 0.159 | 5.373 |
| T134A A138C | False | -4.680 | 1.237 |

**Double mutants at noncanonical basepairing positions in 5TU with statistically significant epistasis**

| Genotype | Fitness of genotype | Epistasis |
|---|---|---|
| G50C T101G | -2.637 | -0.679 |
| G88T A91G | -5.391 | 1.264 |
| G88A A91C | -4.851 | 3.244 |
| G88A A91G | -4.398 | 2.115 |

**Table S3. Epistasis of t1 bp positions.**

**Double mutants at canonical basepairing positions in t1 with statistically significant epistasis**

| Genotype | Restores basepair? | Fitness of genotype | Epistasis |
|---|---|---|---|
| A2T T129A | True | 0.107 | 3.290 |
| C3A G128T | True | -1.030 | 1.761 |
| C3G G128T | True | -1.243 | 1.913 |
| C3G G128C | True | -0.539 | 3.268 |
| A5T T126A | True | -0.780 | 1.042 |
| C11A G122T | True | 0.073 | 1.024 |
| C12A G120T | True | 1.566 | 1.784 |
| C12T G120A | True | 1.630 | 0.580 |
| C15G G117C | True | -3.409 | 1.910 |
| A16G T116C | True | 1.596 | 2.750 |
| A16T T116G | True | 0.557 | 3.963 |
| G17T C115G | True | -0.423 | 3.460 |
| G19A C114A | False | -1.546 | 2.124 |
| C20G G113C | True | -1.319 | 4.456 |
| C20T G113A | True | 1.398 | 3.670 |
| C28T G110A | True | 0.874 | 3.356 |
| C28G G110A | False | -1.493 | 1.606 |
| C28A G110A | False | -0.364 | 4.252 |
| C28T G110T | False | -1.442 | 1.330 |
| A29G T109C | True | 0.997 | 4.391 |
| C31A G107C | False | -4.910 | 6.387 |
| G35A C98G | False | -6.500 | 2.929 |
| G36A C97A | False | -3.861 | 1.670 |
| G36A C97T | True | -0.157 | 1.230 |
| G36T C97A | True | -0.428 | 5.309 |
| A37T T96G | True | -0.312 | 0.575 |
| A37G T96A | False | -3.443 | -2.363 |
| A37C T96G | True | 0.787 | 1.750 |
| T38G A95C | True | 0.185 | 1.848 |
| A41T T92C | False | -1.212 | 0.777 |
| A41C T92A | False | -2.214 | 1.537 |
| A41T T92A | True | -0.903 | 3.034 |
| G44T C89A | True | 0.219 | 2.654 |
| G44T C89G | True | -0.509 | 1.839 |
| G44A C89A | False | -1.045 | 1.023 |

| | | | |
|---|---|---|---|
| G45T C88T | False | -0.272 | 1.081 |
| G45C C88T | False | -0.355 | 1.264 |
| A46T T87A | True | -0.324 | 0.774 |
| G47A C86A | False | -0.754 | 0.697 |
| G47C C86A | False | -0.437 | 1.848 |
| G47T C86G | True | -0.954 | 1.586 |
| G48A C85T | True | -0.302 | 1.323 |
| G48A C85A | False | -0.739 | 1.876 |
| A50T C79T | False | -1.760 | -0.702 |
| G51C C80G | True | -1.849 | 7.637 |
| G52T T81A | True | -2.787 | -0.955 |
| C56G G77C | True | -0.126 | 0.763 |
| T59A A74T | True | -0.959 | -0.794 |
| G60C C73T | False | -1.409 | -1.136 |
| G61C C72T | False | 0.273 | 2.301 |
| G61A C72A | False | -0.462 | -0.660 |
| G63T C70A | True | -0.179 | 2.505 |

**Double mutants at noncanonical basepairing positions in t1 with statistically significant epistasis**

| Genotype | Fitness of genotype | Epistasis |
|---|---|---|
| C8G C124T | 0.949 | -1.168 |
| C13G A119T | 0.114 | 1.994 |
| C13A A119T | 0.145 | 2.100 |
| A103C C104A | -2.667 | 3.788 |
| C49T A84C | -1.900 | -1.315 |
| C49T C82G | -0.241 | 0.916 |
| C53A T78A | 0.532 | 0.730 |

**Table S4. Oligonucleotide sequences.**

All sequences are written in a 5'-to-3' direction. DNA sequences are coloured grey. RNA sequences are coloured black. All RNAs were denaturing PAGE-purified, and DNAs were not, unless otherwise noted ('GP').

| Application | Oligonucleotide | Sequence (5'-3')<br>& Origin |
|---|---|---|
| Fill-in | 5T7 | GATCGATCTCGCCCGCGAAATTAATACGACTCAC<br>TATA<br>Sigma |
| | HDVrt | CTTCTCCCTTAGCCTACCGAAGTAGCCCAGGTCGG<br>ACCGCGAGGAGGTGGAGATGCCATGCCGACCC<br>Sigma, GP |
| Transcription of 5TU | 5TU | pGGAUCUUCUCGAUCUAACAAAAAGACAAAUC<br>UGCCACAAAGCUUGAGAGCAUCUUCGGAUGCAG<br>AGGCGGCAGCCUUCGGUGGCGCGAUAGCGCCAA<br>CGUUCUCAACUAUGACACGCAAAACGCGUGCUC<br>CGUUGAAUGGAGUUUAUCAUG<br><br>GMP transcribed from DNA template constructed from GoTaq PCR using (5TU-5T7-f and 5TU-HDVrec-r) as PCR template, and 5T7 and HDVrt as primers. |
| | 5TU-5T7-f | GATCGATCTCGCCCGCGAAATTAATACGACTCAC<br>TATAGGATCTTCTCGATCTAACAAAAAGACAAA<br>TCTGCCACAAAGCTTGAGAGCATCTTCGGATGCA<br>GAGGCGGCAGCCTTCGG<br>Sigma |
| | 5TU-HDVrec-r | GATGCCATGCCGACCCCATGATAAACTCCATTCA<br>ACGGAGCACGCGTTTTGCGTGTCATAGTTGAGAA<br>CGTTGGCGCTATCGCGCCACCGAAGGCTGCCGCC<br>Sigma |
| Transcription of t1 | t1 | pGACCAAUCUGCCCUCAGAGCUCGAGAACAUCU<br>UCGGAUGCAGAGGAGGCAGGCUUCGGUGGCGCG<br>AUAGCGCCAACGUCCUCAACCUCCAAUGCAUCC<br>CACCACAUGAUGAUGCCUGAAGAGCCUUGGUUU<br>UUUG<br><br>GMP transcribed from DNA template constructed from GoTaq PCR using (t1-5T7-f and t1-HDVrec-r) as PCR template, and 5T7 and HDVrt as primers. |
| | t1-5T7-f | CCCGCGAAATTAATACGACTCACTATAGACCAAT<br>CTGCCCTCAGAGCTCGAGAACATCTTCGGATGCA |

| | | |
|---|---|---|
| | | GAGGAGGCAGGCTTCGGTGGCGCGATAGCGCCAA CGT<br>Sigma |
| | t1-HDVrec-r | GATGCCATGCCGACCCCAAAAAACCAAGGCTCTT CAGGCATCATCATGTGGTGGGATGCATTGGAGGT TGAGGACGTTGGCGCTATCGCGCCACCG<br>Sigma |
| Testing ribozyme activity (Fig 1b, SI Fig 1c, Fig 3e) | Biocy3P10 | Biotin-cy3-CUGCCAACCG<br>IDT<br><br>(used as primer in ribozyme-mediated extensions) |
| Template for testing ribozyme activity (Fig 1b) | t6FP10GAA18 | pppGGUCCAUUCUUCUUCUUCUUCUUCUUCUUCU UCUUCUUCUUCUUCUUCUUCUUCUUCUUCCGGU UGGCAG<br><br>Transcribed using fill-in of 5T7 with:<br>CTGCCAACCGGAAGAAGAAGAAGAAGAAGAAGA AGAAGAAGAAGAAGAAGAAGAAGAAGAAGAAT GGACCTATAGTGAGTCGTATTAATTTCGCGGGCG AGATCGATC<br>Sigma |
| Template for testing ribozyme activity (SI Fig 1c) | t6FP10mix | pGGUCCAUGGACCUAUGCGUUCGAAGGUCGCCG GUUGGCAG<br><br>GMP transcribed using fill-in of 5T7 with:<br>CTGCCAACCGGCGACCTTCGAACGCATAGGTCCA TGGACCTATAGTGAGTCGTATTAATTTCGCGGGC GAGATCGATC<br>Sigma |
| Template for testing ribozyme activity (Fig 3e) | tP10CGU11 | pGGACGACGACGACGACGACGACGACGACGACG ACGCGGUUGGCAG<br><br>GMP transcribed using fill-in of 5T7 with:<br>CTGCCAACCGCGTCGTCGTCGTCGTCGTCGTCGTC GTCGTCGTCCTATAGTGAGTCGTATTAATTTCGCG GGCGAGATCGATC<br>Sigma |
| Ribozyme evolution and fitness landscape measurement | ForceGG | AACAAACAACAAAACAAACAAACAGG<br>Sigma |
| | t1rec | AACAAACAAACAAACAAAAAAG<br>Sigma |
| | HDVrec | GATGCCATGCCGACCC<br>Sigma |
| | t5_tri12x12 | GATCGATCTCGCCCGCGAAATTAATACGACTCAC TATAGGTCCGAAAGGACCTATTATTATTATTATTA |

| | | |
|---|---|---|
| | | TTATTATTATTATTATTATCGGTTGGCAGAACAAA<br>CAAACAAACAAACAAACAAACAAACAACAAAAC<br>AAACAAACAGG<br>IDT, GP |
| | AdeHDVLig | $A^{p\text{-}p}$GGGTCGGCATGGCATC-$C_3$ spacer<br>Treatment of 20 μM HDVLig with 5’ DNA adenylation kit (NEB) 65˚C 2 h, neutral phenol/chloroform extracted and precipitated in 72% ethanol. |
| | HDVLig | $^{p}$GGGTCGGCATGGCATC-$C_3$ spacer<br>IDT, GP |
| | P51HDVba | AATGATACGGCGACCACCGAGATCTACACTCTTT<br>CCCTACACGACGCTCTTCCGATCTNNNATCAGAT<br>GCCATGCCGACCC<br>IDT |
| | P52HDVba | AATGATACGGCGACCACCGAGATCTACACTCTTT<br>CCCTACACGACGCTCTTCCGATCTNNNCGATGAT<br>GCCATGCCGACCC<br>IDT |
| | P53HDVba | AATGATACGGCGACCACCGAGATCTACACTCTTT<br>CCCTACACGACGCTCTTCCGATCTNNNTTAGGATG<br>CCATGCCGACCC<br>IDT |
| | P510HDVba | AATGATACGGCGACCACCGAGATCTACACTCTTT<br>CCCTACACGACGCTCTTCCGATCTNNNTGAAGAT<br>GCCATGCCGACCC<br>IDT |
| | P7forceGG | CAAGCAGAAGACGGCATACGAGATGTGACTGGA<br>GTTCAGACGTGTGCTCTTCCGATCNNNAACAAAC<br>AACAAAACAAACAAACAGG |
| | 5TU-F1 | AACAAACAACAAAACAAACAAACAGGATCTTCTC<br>GATCTAACAAAAAAGACAAATCTGCCACAAAGCT<br>TGAGAGCATC<br>IDT<br><br>Underlined bases were spiked with 1% incorrect bases each (97% correct bases) |
| | 5TU-F2 | GGATGCAGAGGCGGCAGCCTTCGGTGGCGCGATA<br>GCGCCAACGTTCTCAACTATGACACGCAAAACGC<br>GTGCTCC<br>IDT<br><br>Underlined bases were spiked with 1% incorrect bases each (97% correct bases) |
| | 5TU-R1 | GGCTGCCGCCTCTGCATCCGAAGATGCTCTCAAG<br>CTTTGTGGCAGA |

| | | |
|---|---|---|
| | | IDT<br><br>Underlined bases were spiked with 1% incorrect bases each (97% correct bases) |
| | 5TU-R2 | GATGCCATGCCGACCCCATGATAAACTCCATTCA<br>ACGGAGCACGCGTTTTGCGTGTCATAG<br>IDT<br><br>Underlined bases were spiked with 1% incorrect bases each (97% correct bases) |
| | t1-F1 | AACAAACAAACAAACAAAAAAGACCAATCTGCC<br>CTCAGAGCTCGAGAACATCTTCGGATGCAGAGGA<br>GGCAGGCTTCGGTGGCGCGATAGCGCCAACGT<br>IDT<br><br>Underlined bases were spiked with 1% incorrect bases each (97% correct bases) |
| | t1-R1 | GATGCCATGCCGACCCCAAAAAACCAAGGCTCTT<br>CAGGCATCATCATGTGGTGGGATGCATTGGAGGT<br>TGAGGACGTTGGCGCTATCGCGCCACCG<br>IDT<br><br>Underlined bases were spiked with 1% incorrect bases each (97% correct bases) |
| | P51t1rec | AATGATACGGCGACCACCGAGATCTACACTCTTT<br>CCCTACACGACGCTCTTCCGATCTNNNAACGAAC<br>AAACAAACAAACAAAAAAG<br>IDT |
| | P52t1rec | AATGATACGGCGACCACCGAGATCTACACTCTTT<br>CCCTACACGACGCTCTTCCGATCTNNNCGTGAAC<br>AAACAAACAAACAAAAAAG<br>IDT |
| | P53t1rec | AATGATACGGCGACCACCGAGATCTACACTCTTT<br>CCCTACACGACGCTCTTCCGATCTNNNGAACAAC<br>AAACAAACAAACAAAAAAG<br>IDT |
| | P54t1rec | AATGATACGGCGACCACCGAGATCTACACTCTTT<br>CCCTACACGACGCTCTTCCGATCTNNNTCGAAAC<br>AAACAAACAAACAAAAAAG<br>IDT |

| | | |
|---|---|---|
| | P7HDVba | CAAGCAGAAGACGGCATACGAGATGTGACTGGA GTTCAGACGTGTGCTCTTCCGATCNNNGATGCCAT GCCGACCC<br>IDT |

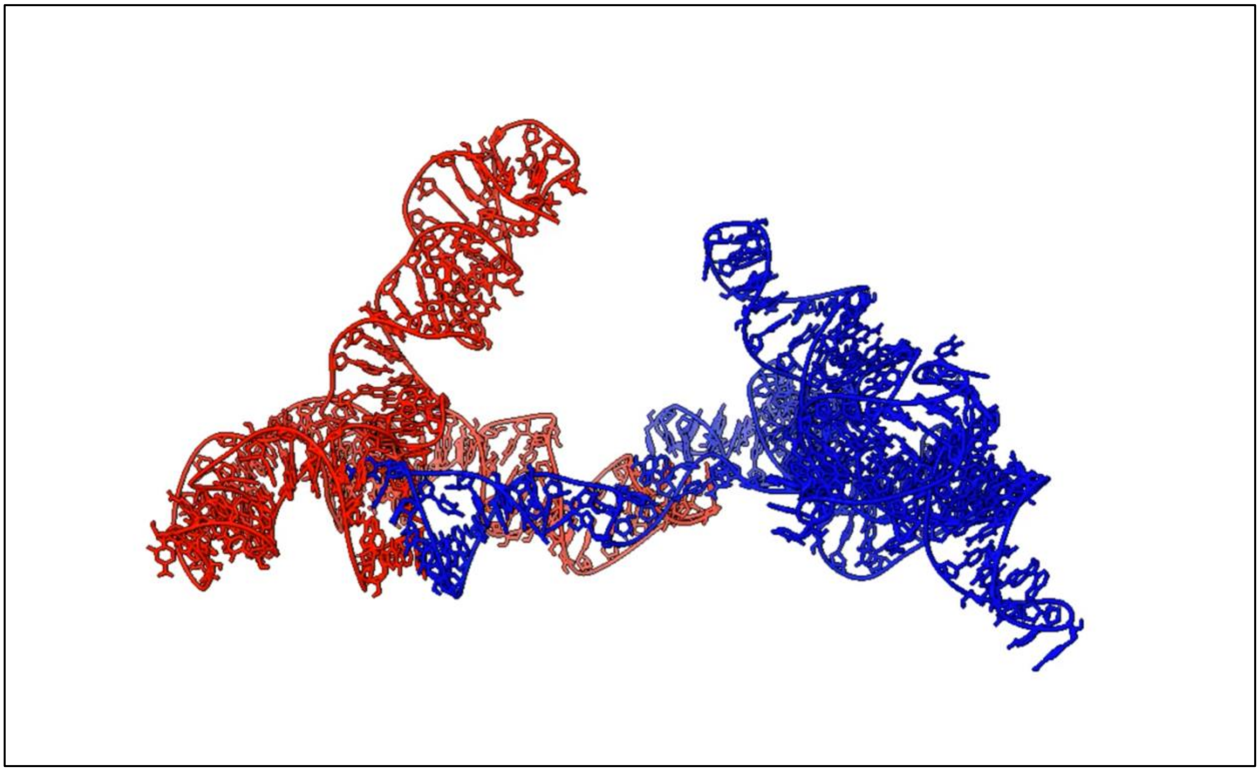

**Movie S1. 3D variability analysis of the TPR.**

Movie showing 3D variability analysis of the TPR from the 126,690-particle stack shown in Fig. S2.

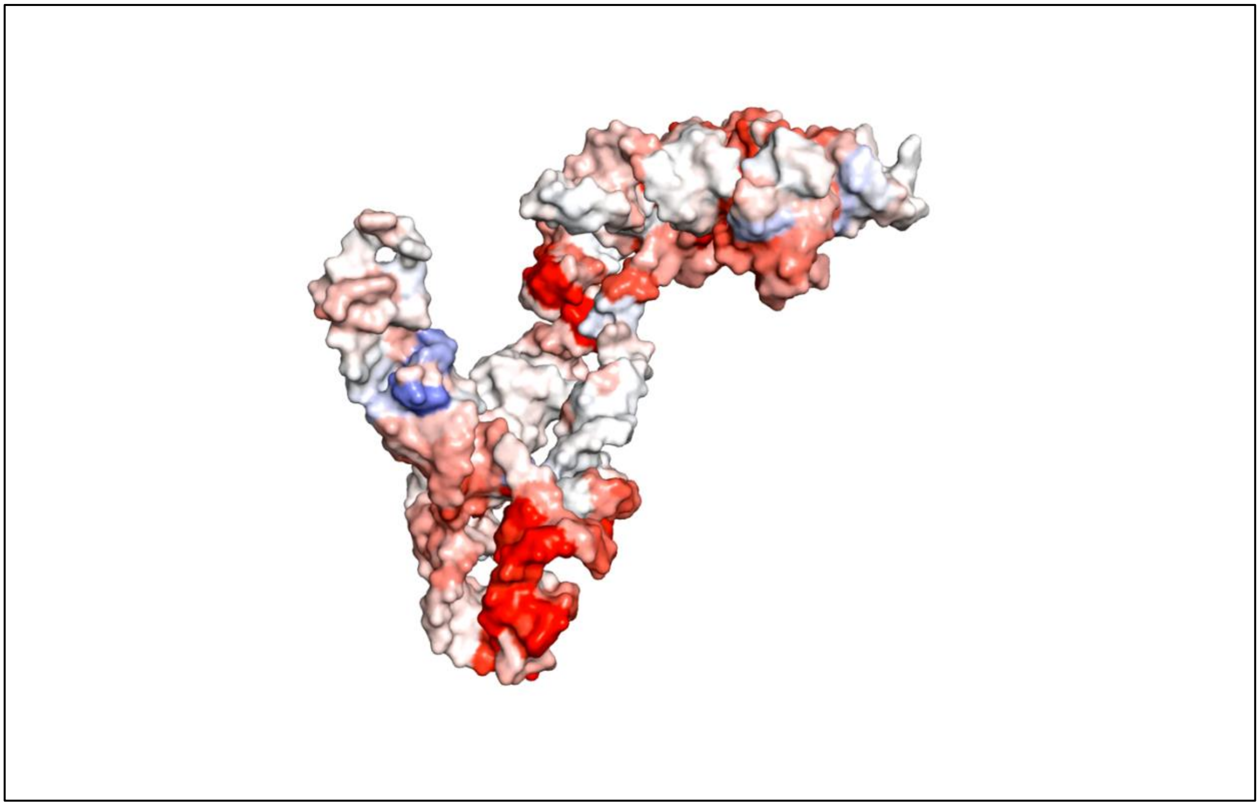

**Movie S2. Average fitness values on the TPR tertiary structure.**

Movie showing the average fitness values for a given nucleotide position on the TPR tertiary structure.
